# RNAquarium: an archive-scale atlas of zebrafish gene expression coupled with pan-taxonomic profiling reveals diverse viral drivers of transcriptomic states

**DOI:** 10.64898/2026.06.17.732950

**Authors:** Yttria Aniseia, Eric Waltari, Hejin Huang, Leandro Lima, Gibraan Rahman, Max Frank, Andy Zhou, Yang-Joon Kim, Jacob Paras, Steven Baker, Yasin Şenbabaoğlu, Duo Peng, Keir Balla

## Abstract

Zebrafish RNA-seq studies span diverse developmental, physiological, and disease contexts, yet most analyses remain confined to individual experiments and disregard the non-zebrafish component of the data. We present RNAquarium, a scalable framework for joint transcriptomic and metatranscriptomic analysis of RNA-seq data and apply it to all publicly available zebrafish RNA-seq datasets in the Sequence Read Archive. RNAquarium captures transcriptomic structure across development and tissues, reveals diverse microbial and viral associations, and identifies previously undescribed zebrafish viruses including a close relative of human influenza B virus linked to distinct host transcriptional states. We further demonstrate that archive-scale transcriptomes can support foundation-model training and prediction of infection-associated transcriptomic signatures. RNAquarium provides an open framework and interactive portal for exploring the breadth of zebrafish gene expression patterns and associated taxa profiled across a large research community. Moreover, our framework establishes a generalizable strategy for integrating transcriptomic and metatranscriptomic analyses across the diversity of life represented in public sequencing archives.

## INTRODUCTION

RNA sequencing (RNA-seq) is one of the primary tools for profiling biological systems due to its versatility in yielding significant information across diverse contexts and scales [1]. The colossal collection of publicly available sequencing data hosted by the Sequence Read Archive [2] (SRA) is a testament to the usefulness of RNA-seq in thousands of studies, and in aggregate provides the potential for revealing latent features that are uniquely apparent at large scales or through alternative analytical approaches.

Advances in computing infrastructure and analysis tools have enabled important discoveries made through massive-scale investigations of RNA-seq data stored in the SRA and other compendia. For example, systematic processing and analysis of thousands of RNA-seq datasets from human, mice, and other research organisms broadly improved annotation of gene structures and networks, illuminated cross-species similarities, and revealed novel transcript usage [3–6]. Furthermore, output from these studies has been used to train models with broad inference capabilities including functional predictions for understudied genes and characterization of novel disease states [7, 8]. One limitation of these studies is that they disregard all sequences that do not align to the subject’s reference transcriptome. Parallel repository-scale work has revealed previously undescribed viruses across the SRA and transformed the archive into longer assemblies of sequences to potentially enable expanded taxonomic annotation of each sequence in every dataset [9, 10] (**Supplemental Table S1**). Linking the taxonomic characterization of all sequences in a dataset with gene expression patterns at compendium scale holds the prospect of illuminating drivers of transcriptomic states in research subjects that were profiled across diverse contexts often along with viruses and other organisms.

The zebrafish (*Danio rerio*) is an ideal subject for coupling gene expression quantification with comprehensive taxonomic profiling at repository scale. As a widely used vertebrate model for foundational biology and human disease research, zebrafish studies have generated vast and diverse RNA-seq datasets across the lifespan in a rich variety of biological contexts from cellular to organismal scales [11]. The recent publication of the first telomere-to-telomere zebrafish genome assembly together with improved gene annotations [12] has further enhanced the accuracy of RNA-seq read alignment and transcript quantification. Many RNA-seq datasets also likely contain reads derived from organisms that naturally associate with zebrafish in research settings, including microbiome-associated bacteria [13], eukaryotic parasites [14], and viruses [15]. Although these associations are often overlooked, they can substantially influence zebrafish transcriptional programs and physiological states [15–17]. However, the diversity, prevalence, and transcriptomic impact of these associated organisms remain largely uncharted.

Here we present RNAquarium and a global characterization of zebrafish transcriptomes from across the SRA. RNAquarium is a distributed and generalizable workflow that processes RNA-seq data at scale to produce gene expression quantification for a subject organism paired with *de novo* assembled sequences that are taxonomically classified. We applied RNAquarium to the public zebrafish RNA-seq archive and used representation learning to provide contextual information for every gene in the genome along with an atlas of associated taxa. We identify organisms from across the tree of life and detect viruses in thousands of datasets that are associated with altered transcriptomic signatures. Beyond supporting infection biology discovery, such organism-scale compendia provide the substrate for a variety of strategies aimed at modeling the diverse transcriptomic states of complex biological systems.

## RESULTS

### RNAquarium enables scalable processing of RNA-seq data from zebrafish across the public archive

We developed RNAquarium, an open-source pipeline for repository-scale RNA-seq processing that outputs gene expression quantification for a reference organism (referred to as host) paired with *de novo* assembly and taxonomic classification of non-host transcripts. The first stage of the workflow retrieves raw sequencing reads from the SRA and pre-processes them for host gene expression analysis. In parallel, a host-filtering step efficiently screens all reads against a collection of host genome variants. Non-host reads are then passed to the second stage of the workflow, where they are assembled into transcript contigs, taxonomically identified, and quantified across all sequencing runs (**Figure 1A; Supplemental Figure S1**).

**Figure 1.**
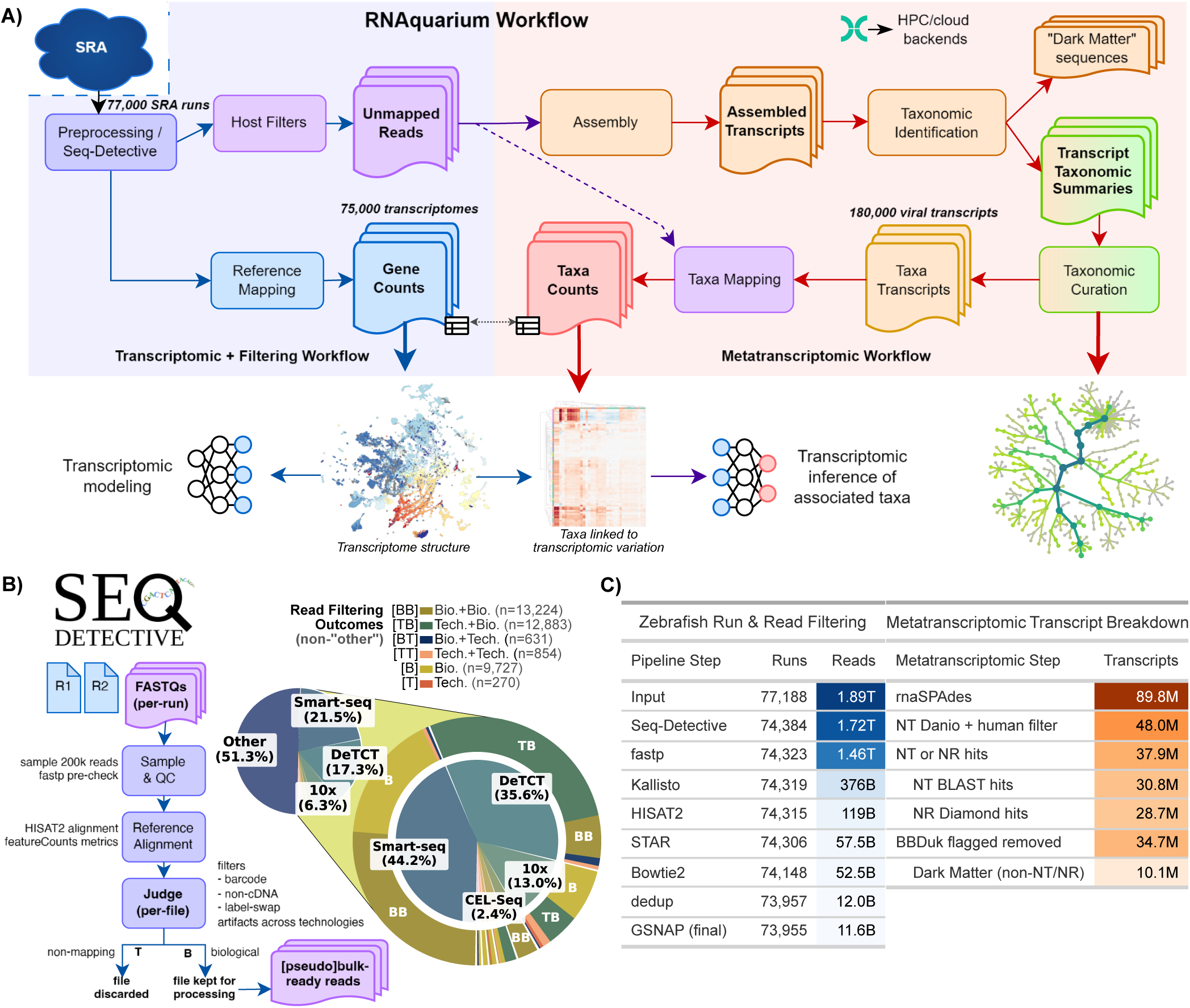
Gene expression quantification and pan-taxonomic profiling of all zebrafish RNA-seq accessions in the SRA. (A) Schematic overview of the RNAquarium computational workflow. The workflow is divided into two main components: a transcriptomic and filtering workflow, which downloads sequencing reads from the SRA, preprocesses reads, maps them to the reference host genome, and filters out host-derived reads; and a metatranscriptomic workflow, which performs transcript assembly, taxonomic identification, and abundance estimation. Downstream outputs are discussed in later figures. (B) Seq-Detective identifies and filters technical mate reads that fail to map to the reference species. Manually annotated sequencing technologies are shown as pie charts. (C) Pipeline outcome summaries and assembled-transcript counts per step, or per category for indented rows.

A key challenge in the read pre-processing step is accurately determining read configuration, as the biological (B) and technical (T) labels in SRA metadata do not reliably distinguish single-end from paired-end reads for host genome alignment. To address this, we developed Seq-Detective, which classifies each sequencing run’s read mate configuration as biological (B) or technical (T) using heuristics based on alignment rate, expression sparsity, and read length (**Figure 1B**). We validated Seq-Detective’s classification performance against 150 human-curated filtering decisions representative of diverse configurations (see STAR Methods for details; **Supplemental Figure S2**). Applied across the full accession set, Seq-Detective rejected 2,800 unmappable accessions and recovered 14,825 accessions that would otherwise have been mis-processed, thus minimizing the influence of technical sequences in further analyses.

Both stages of the RNAquarium workflow are implemented in Nextflow and support distributed execution with thousands of concurrent jobs, dynamic resource scaling, and fault-tolerant caching. These design choices enabled end-to-end processing of the full zebrafish SRA archive, comprising 77,188 sequencing runs and 1.89 trillion reads (**Figure 1C**; see STAR Methods for complete pipeline details including steps, component packages, and databases). Alignment to the zebrafish genome assembly GRCz12tu [12] yielded non-zero counts for 43,091 out of 49,285 annotated genes across 74,275 runs. In the metatranscriptomic workflow, 11.6 billion reads (0.7% of input) across 73,955 runs were retained as putative non-zebrafish sequence, which were pooled at the BioProject level and assembled into 89.8 million transcripts. Of the 48 million transcripts that were not classified as human or zebrafish, 37.9 million sequences were annotated as deriving from one of 108,836 prospective taxa (**Supplemental Figure S1**). Completion of the pipeline from download to transcriptomic and annotated metatranscriptomic outputs took *≈*30,000 CPU hours, enabling analysis of gene expression across the zebrafish RNA-seq archive in parallel with associated taxa.

### Constructing a pan-archive atlas of zebrafish transcriptomes and metatranscriptomes spanning all life stages and tissues

We parsed metadata from the SRA and the Gene Expression Omnibus for all zebrafish accessions and assigned each a developmental stage and anatomy ontology [18] (see STAR Methods for details). Projecting all 74,275 transcriptomes overlaid with curated metadata labels highlighted clustering that corresponded to developmental stage (**Figure 2A**).

**Figure 2.**
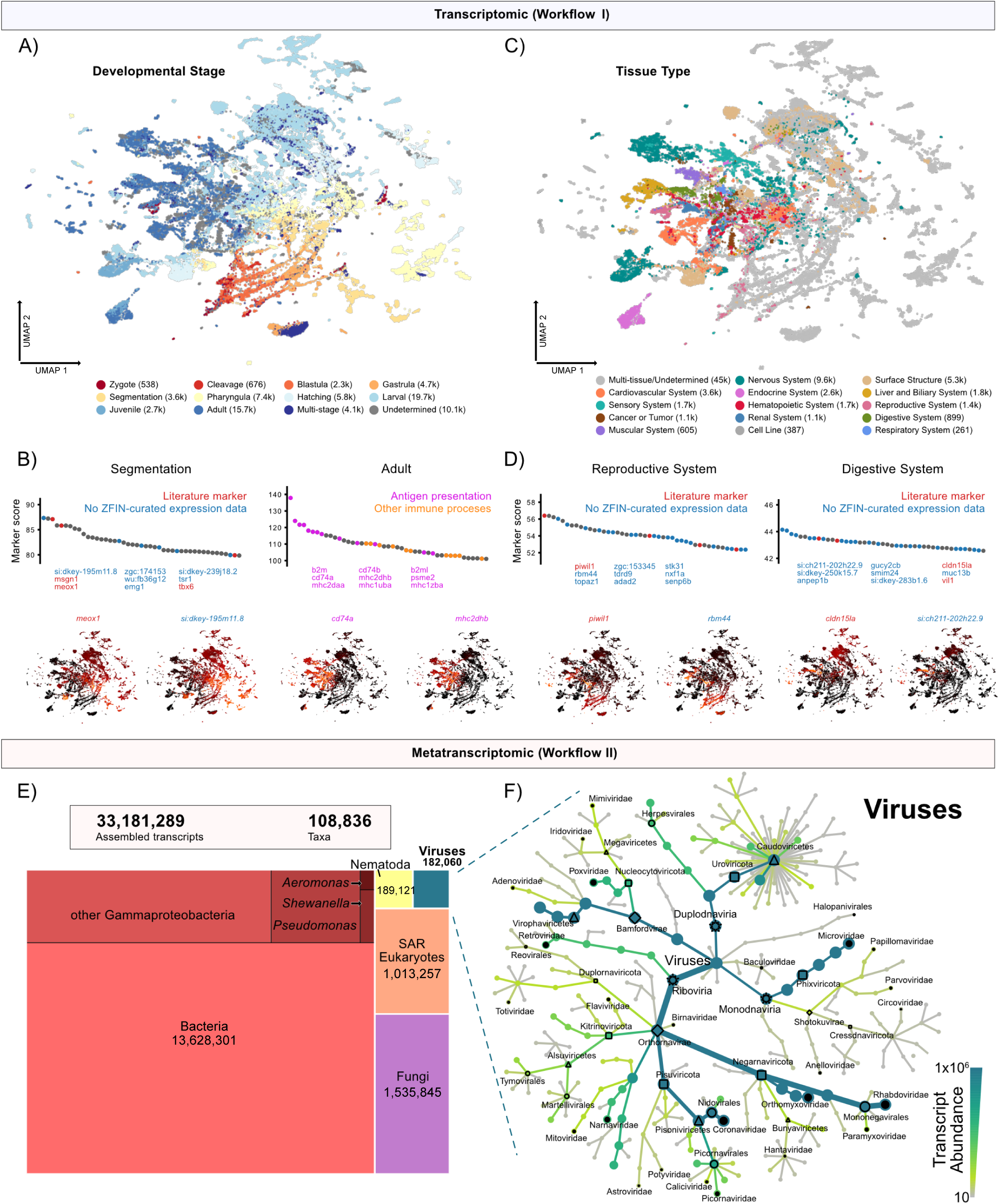
Diversity of zebrafish transcriptomes and associated taxa across the SRA. (A) UMAP representation of zebrafish expression from 74,275 SRA accessions, colored by developmental stage. Numbers in parentheses indicate number of SRA datasets corresponding to each stage. (B) Top genes found to be enriched in segmentation (left) or adult (right) stages compared to other stages. For the segmentation-enriched genes, colors indicate whether they have been previously described (red) or not found with expression data in ZFIN (blue), while in adult-enriched genes colors indicate whether they are involved in antigen presentation (cyan) or other immune processes (orange). Two example UMAPs for both segmentation-enriched and adult-enriched genes are also shown (*meox1* & *si:dkey-195m11.8*, lower left; *cd74a* & *mhc2dhb*, lower right). See **Supplemental Figure S3A** for full gene lists across additional developmental stages. (C) UMAP representation of zebrafish expression from 74,275 SRA accessions, colored by broad tissue category. Numbers in parentheses indicate number of SRA accessions corresponding to each tissue. For interactivity and to see tissue types found in fewer than 250 SRA accessions, see the portal UMAP viewer. (D) Top genes found to be enriched in reproductive system tissues (left) or digestive system tissues (right) compared to other tissues. For both gene panels, colors indicate whether they have been previously described (red) or not found with expression data in ZFIN (blue). Two example UMAPs for both reproductive system-enriched and digestive system-enriched genes are also shown (*piwil1* & *rbm44*, lower left; *cldn15la* & *si:ch211-202h22.9*, lower right). See **Supplemental Figure S3B** for full gene lists across additional tissue categories. (E) Treemap indicating numbers of selected non-host transcripts across taxonomic categories, with total assembled transcripts and taxa at top. (F) Metacoder heat tree of virus transcripts, showing classification to families. Shapes on heat tree indicate taxonomic level: stars = viral realms; diamonds = kingdoms; squares = phylum; open circles = orders; black circles = families. Both width of tree branches and shade are scaled by cumulative search result transcript length x coverage ‘abundance’ values across all transcripts in each clade.

Comparing transcriptomes between developmental stages revealed enrichment for genes that are classical markers of each life stage in addition to genes without previously reported expression data in the Zebrafish Information Network (ZFIN) [19] (**Figure 2B; Supplemental Figure S3)**. For example, we observed enrichment for somite-related genes during the somite-forming segmentation stage of development [20], including *msgn1* and *tbx6* that have been used to transgenically drive expression in somites [21, 22] (**Figure 2B**). Transcriptomes from adult zebrafish were enriched for genes involved in antigen presentation and other immune functions, highlighting adaptive immunity and other immune processes as the predominant drivers of gene expression differences between mature and developing life stages.

Transcriptomes also clustered based on their tissue of origin (**Figure 2C)**. In reproductive system transcriptomes we observed enrichment for several genes that have been used to create gonadal transgenic lines including *piwil1* [23], *ddx4* [24], and *dnd1* [25], among several other genes without previously reported expression data (**Figure 2D; Supplemental Figure S3**). Digestive system transcriptomes were enriched for *cldn15la*, which has been used to drive transgenic expression in the intestine [26], but most enriched genes in this tissue category lacked previously reported expression data (**Figure 2D**). Thus, a pan-archive atlas of zebrafish transcriptomes captures previously described expression patterns and extends quantitative contextual information to every gene in the genome.

We classified the millions of assembled sequences in zebrafish-associated metatranscriptomes as deriving from diverse taxa across the tree of life (**Figure 2E**). Bacteria accounted for the largest number of sequences with millions classified as *Aeromonas*, *Pseudomonas*, *Shewanella*, and other members of the class Gammaproteobacteria that frequently dominate zebrafish microbiota [13] (**Supplemental Figure S4**). Among eukaryotes we identified several previously documented intracellular and extracellular parasites of laboratory zebrafish including fungal pathogens in the phylum Microsporidia [14], water molds in the phylum Oomycota [27], and cancer-causing nematodes in the order Trichinellida [28, 29] (**Supplemental Figure S4**). Viral taxa spanned all Baltimore classes and included bacteriophage and viruses that naturally infect zebrafish [15, 30, 31] (**Figure 2F**). In addition to detecting previously reported associations, a pan-archive atlas of zebrafish metatranscriptomes reveals thousands of associations with known and novel taxa that potentially influence zebrafish gene expression patterns and physiology (**Supplemental Figures S1,S4**).

### AI modeling of archive-wide transcriptomes and analysis of gene representations

We leveraged the depth and scale of zebrafish RNA-seq data in the RNAquarium to produce gene representations using the transformer-based Geneformer [32] architecture. We pretrained foundation models with 74,275 bulk and pseudo-bulk transcriptomes (excluding Zebrahub data [33]) from the RNAquarium (hereafter RQ-GF). As controls, we pretrained equivalent models on 85,213 single-cell transcriptomes from the Zebrahub atlas of zebrafish development [33], the Genecorpus-30M human RNA-seq dataset [32], or random gaussian noise (**Figure 3A**).

**Figure 3.**
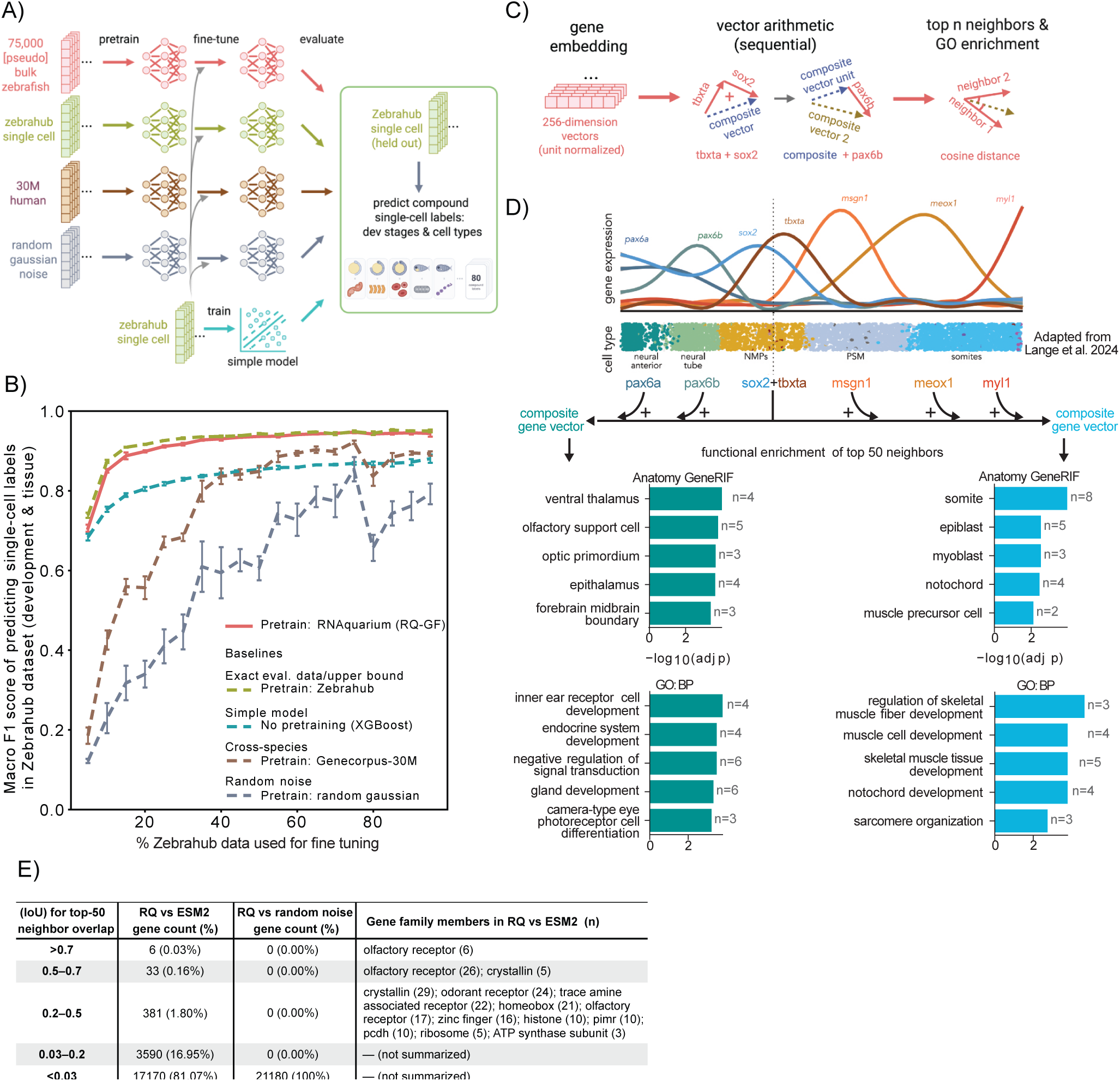
AI modeling of archive-wide transcriptomes and analysis of gene representations. (A) Schematic workflow of the single-cell label prediction benchmarking pipeline. Foundation models using the Geneformer architecture were pretrained on four distinct inputs: RNAquarium bulk/pseudo-bulk zebrafish transcriptomes (RQ-GF), Zebrahub single-cell transcriptomes (upper bound control), Genecorpus-30M human transcriptomes (cross-species control), or random Gaussian noise. Pretrained weights were then fine-tuned to predict compound tissue and developmental stage labels from the Zebrahub atlas, alongside a baseline XGBoost model without pretraining. (B) Fine-tuning performance across varying training data fractions. Model accuracy evaluated by Macro F1 score across 80 compound cell-type and developmental-stage label classes as a function of the fraction of data utilized during fine-tuning. Error bars indicate standard deviation across validation subsets. (C) Methodology for embedding vector arithmetic. Schematic showing L2-normalization of 256-dimensional gene embeddings followed by sequential vector addition to model the directional influence of transcription factor (TF) cascades on cell fate. (D) Sequential gene embedding arithmetic recapitulates lineage bifurcation from neuromesodermal progenitors. Top, normalized gene expression curves of key regulatory TFs tracking specific cell-type transitions during development (adapted from Lange et al., 2024). Middle, sequential addition of neural-promoting TFs (*pax6b*, *pax6a*) or mesodermal-promoting TFs (*msgn1*, *meox1*, *myl1*) to an initial NMP core vector (*sox2* + *tbxta*) to generate distinct composite cell-state vectors. Bottom, top 50 nearest-neighbor functional enrichment profiles derived from GO Biological Processes (GOBP) and Anatomy GeneRIF for both resulting composite lineages. (E) Comparison of ESM2 and RQ-GF nearest-neighbor gene sets in their embedding spaces. Quantified overlap of the top 50 nearest-neighbor genes between ESM2 and RQ-GF embedding spaces measured by Jaccard Index (Intersection over Union, IoU), benchmarked against a random noise control. Representative gene families displaying moderate-to-high neighborhood concordance are itemized with respective gene counts (n).

We next investigated the content of the representations produced by the RQ-GF model. First, we asked if the RQ-GF gene embeddings could be useful in a scRNA-seq setting. In a compound label prediction task where models must infer both the tissue type and developmental stage of single cells from the Zebrahub scRNA-seq atlas [33], RQ-GF achieved a 0.90 macro-F1 score across 80 classes using as little as 30% of the data for fine-tuning (see STAR Methods, **Figure 3B, Supplemental Figure S5**), approaching a theoretical upper bound obtained when Zebrahub data was used for both pretraining and fine-tuning. An XGBoost-based baseline model, which makes predictions by combining many simple decision trees, consistently underperformed RQ-GF by 0.10–0.15 macro-F1 across most fine-tuning fractions, whereas a noise-trained model baseline performed substantially worse. Notably, initializing from human-pretrained Geneformer weights surpassed the XGBoost baseline at high fine-tuning data fractions (**Figure 3B**). These results demonstrate that RQ-GF captured features at single-cell resolution despite being trained primarily with bulk RNA-seq data.

Gene embeddings from RQ-GF recovered a similar number of unique functional terms as a co-expression network built from the same transcriptomes (11,553 vs. 11,383; **Supplemental Figure S6**), indicating that RQ-GF captures pathway-level organization comparable to standard co-expression analysis. We next asked whether RQ-GF encoded regulatory structure beyond pairwise co-expression relationships (**Figure 3C,D**). To address this question, we examined the differentiation trajectories of neuro-mesodermal progenitors (NMPs), in which temporally coordinated transcription factor (TF) programs drive commitment toward neural or mesodermal fates (**Figure 3D, top**) [33]. To focus on information encoded by vector direction, we L2-normalized the gene embeddings and applied sequential vector arithmetic to model the progression of these regulatory programs (see STAR Methods; **Figure 3C**). Beginning with a composite NMP-state vector defined by *sox2* and *tbxta*, successive addition of the neural TFs *pax6b* and *pax6a* produced a vector whose nearest-neighbor genes were enriched for neurogenesis, visual-system development, and brain patterning (**Figure 3D, bottom left**). By contrast, addition of the mesodermal TFs *msgn1*, *meox1*, and *myl1* yielded a vector whose nearest-neighbor genes were enriched for notochord, skeletal-muscle, and somite development (**Figure 3D, bottom right**). Thus, simple arithmetic in the RQ-GF embedding space recapitulated the divergence of neural and mesodermal developmental programs, suggesting that RQ-GF captures regulatory organization extending beyond the symmetric relationships identified by co-expression analysis.

Finally, we assessed whether RQ-GF and ESM2 [34] encode complementary information: RQ-GF reflects gene expression in cellular/regulatory contexts, whereas ESM2 captures protein sequence features (domains, motifs, folds, and evolutionary relatedness). For each gene, we compared the overlap of nearest-neighbor sets in the two embedding spaces using the Jaccard index (**Figure 3E**). While the overlap was low for most genes as expected (98.02% with Jaccard *<* 0.2), a small subset showed moderate-to-high concordance (381 genes with 0.2–0.5; 39 genes with *>* 0.5). High-overlap genes were enriched for categories where sequence similarity and coordinated regulation coincide: large paralog families with niche expression (olfactory/odorant receptors, crystallins, TAARs), domain-defined transcription factors (homeobox, zinc finger), and tightly co-regulated complexes (histones, ribosomal proteins, ATP synthase subunits). In contrast, comparing RQ-GF to a noise-trained RQ-GF control yielded no genes with Jaccard *>* 0.03, indicating the signal is not an artifact. Thus, deep modeling of zebrafish transcriptomes and proteomes provides complementary representations of gene networks and functions.

### Discovery of diverse zebrafish viruses

We annotated approximately 180,000 transcripts assembled from zebrafish datasets as viral deriving from an estimated 11,000 taxa (**Figure 2; Supplemental Figure S1**). Clustering of the viral sequences reduced the total set of prospective taxa and resulted in 53 taxa that were related to viruses that infect fish, which we quantified across all zebrafish RNA-seq datasets (**Supplemental Data S1**). After filtering out taxa in the *Chuviridae* and *Adintoviridae* families as likely endogenous viral elements [35, 36] along with other insufficiently supported viruses (see STAR Methods), we identified 27 prospective taxa as strong candidate exogenous viruses of zebrafish (**Figure 4A**; **Table 1**). Aligning all assembled transcripts from each taxa to their best viral match revealed that 11 of the zebrafish viruses derived from known species while 16 had median amino acid percent identities of less than 70% and were provisionally named as novel species (**Table 1; Supplemental Figure S7**). Three of the known virus species derived from studies that had intentionally infected zebrafish [37–41], two were previously discovered in zebrafish [15, 31], and six had not been previously described in zebrafish. The novel viruses included prospective members of the *Caliciviridae*, *Hantaviridae*, *Nanhypoviridae*, *Orthomyxoviridae*, *Paramyxoviridae*, *Picornaviridae*, and *Poxviridae* families. Most zebrafish viruses were detected in more than one study and in total were detected in hundreds of BioProjects and thousands of accessions (**Table 1; Supplemental Data S1**), illuminating high viral prevalence across zebrafish studies.

**Figure 4.**
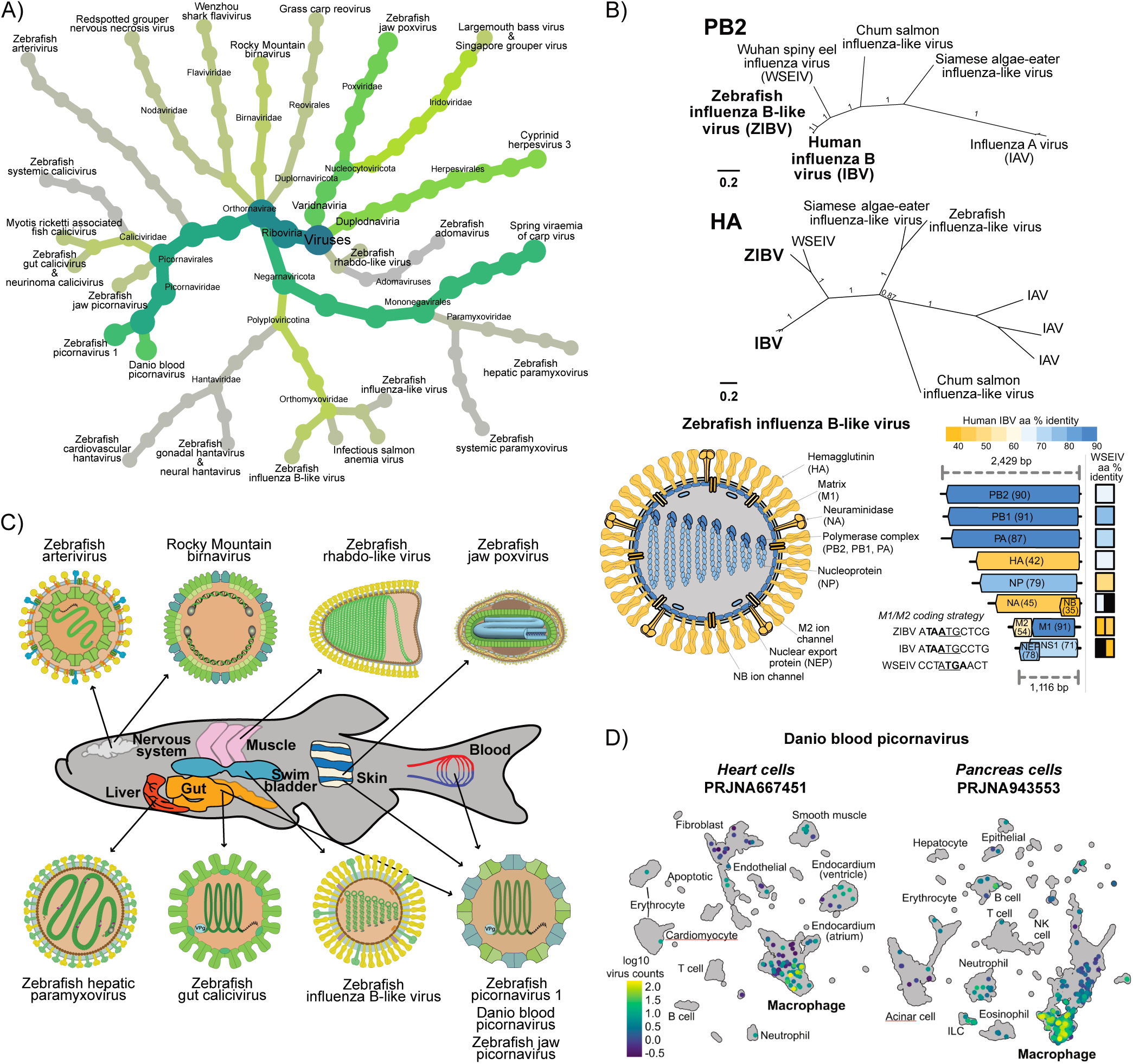
Diversity of evolutionary histories and tissue tropisms among zebrafish viruses. (A) Lineage tree visualization of the prospective zebrafish viruses detected and discovered in the RNAquarium. Both width of tree branches and shade are scaled by cumulative search result transcript length x coverage ‘abundance’ values across all transcripts in each clade. (B) Genome sequence features of influenza viruses discovered in zebrafish. Top, unrooted maximum-likelihood phylogenies of proteins encoded on segments 1 and 4 are plotted with support values, with the prospective Zebrafish influenza B-like virus (ZIBV) and human influenza B virus (IBV) in bold. Bottom left: schematic of the prospective Zebrafish influenza B-like virus (ZIBV) particle is colored based on the protein sequence similarity to human influenza B virus (IBV), with highly similar sequences in blue and divergent sequences in yellow. Bottom right: schematic of ZIBV’s eight genomic segments is similarly colored with percent amino acid identity in parentheses along with boxes denoting sequence similarity to Wuhan spiny eel influenza virus (WSEIV). Segment 7 of IBV and IBV-like viruses encodes two proteins (M1 and M2) separated by variable stop (bolded) and start (underlined) codons plotted as in [64]. (C) Visualization of prevalent zebrafish viruses by tissue tropism. (D) Detection of Danio blood picornavirus in single-cell RNA-seq datasets. Virus counts are plotted on UMAPs annotated with cell type labels.

**Table 1.**
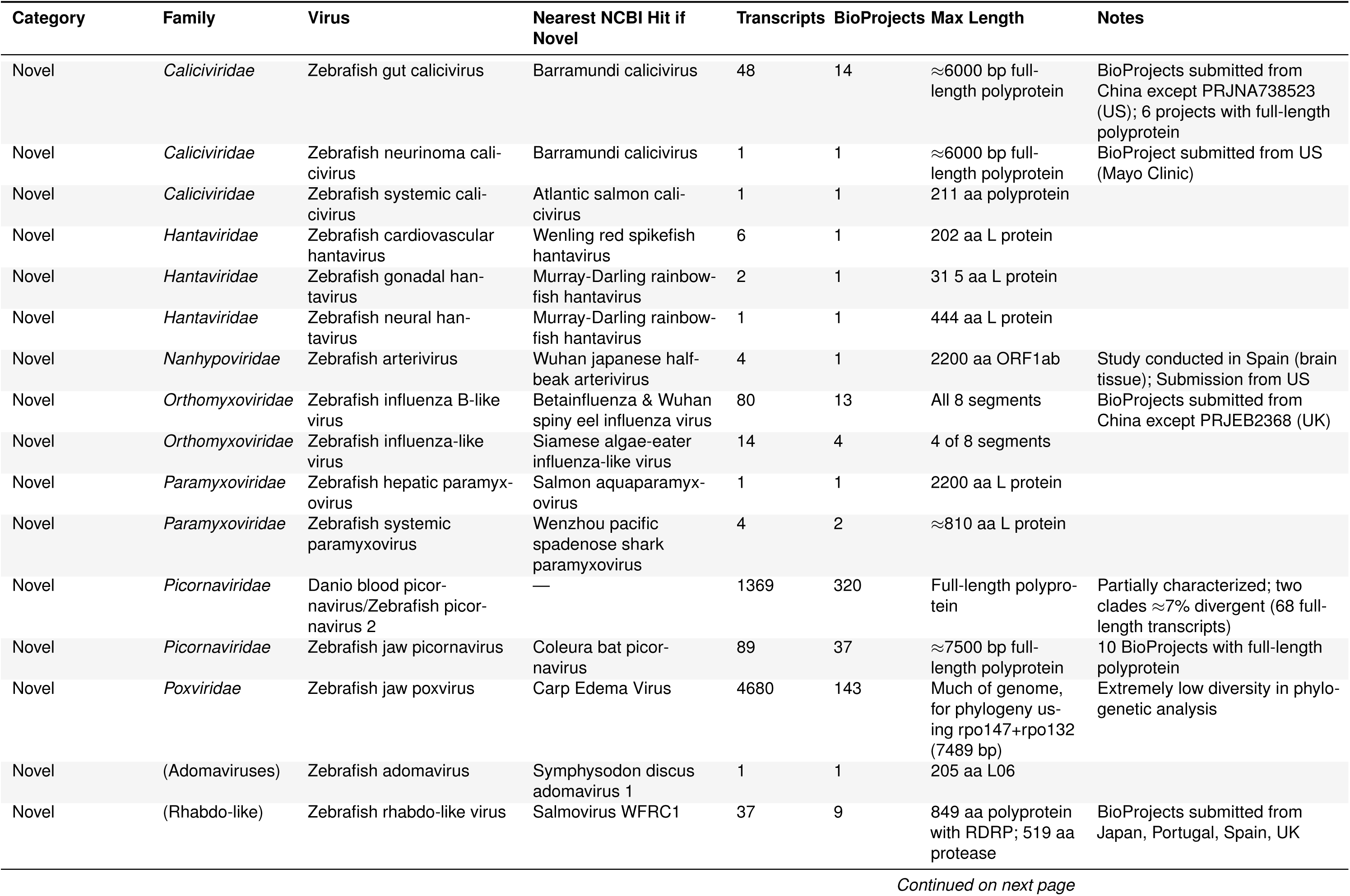
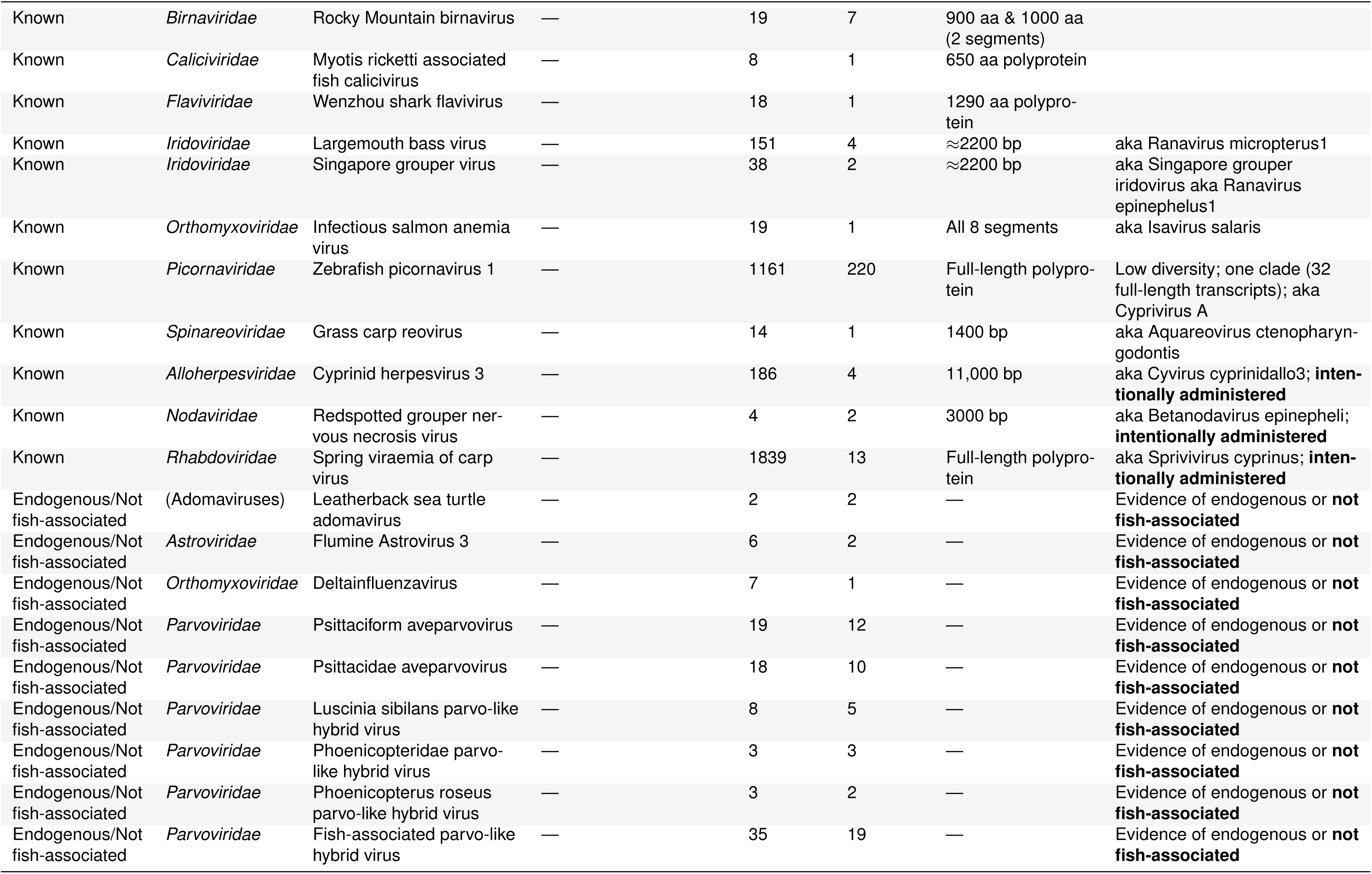
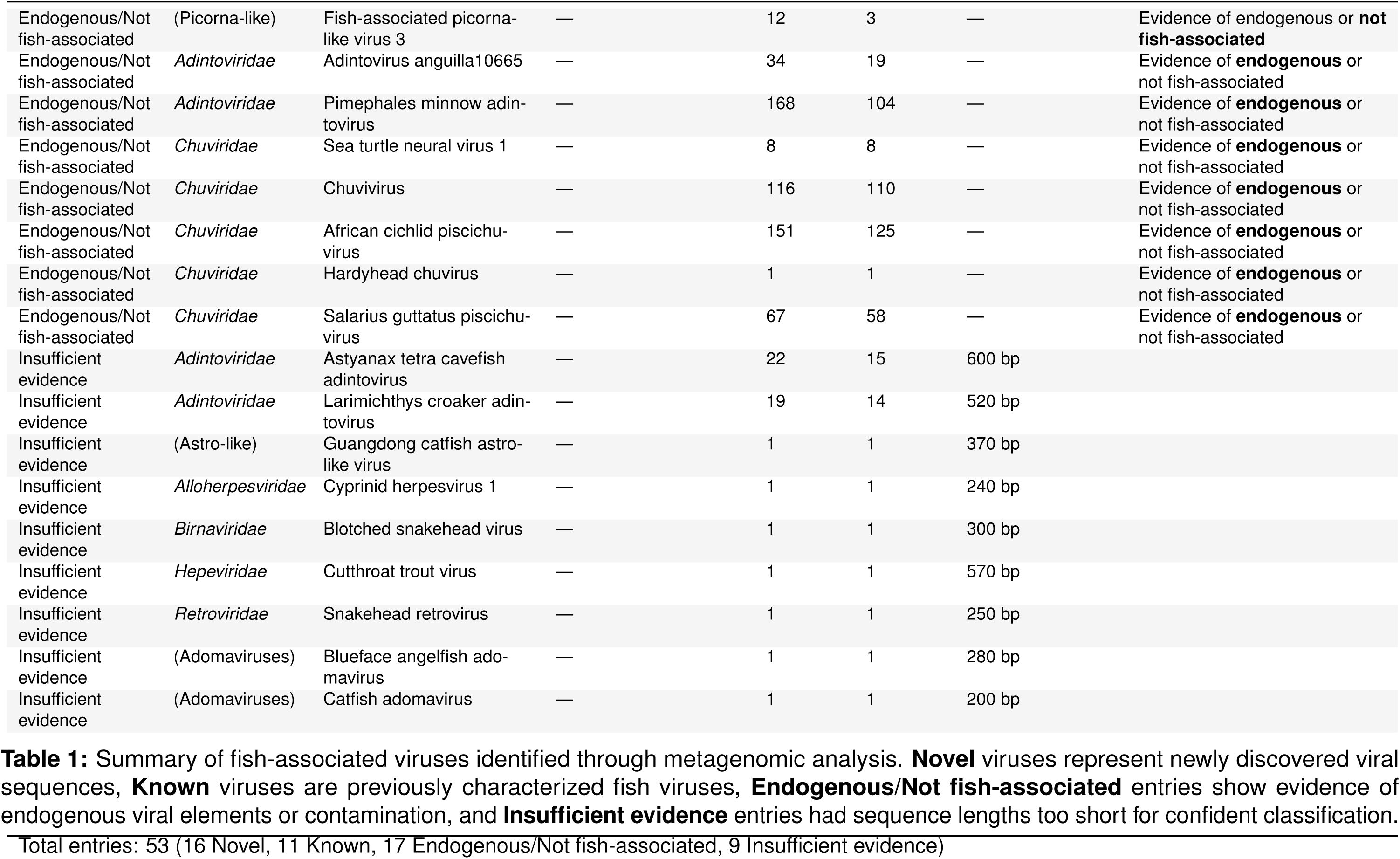
Summary of putative fish-associated viral transcripts.

| Category | Family | Virus | Nearest NCBI Hit if Novel | Transcripts | BioProjects | Max Length | Notes |
|---|---|---|---|---|---|---|---|
| Novel | *Caliciviridae* | Zebrafish gut calicivirus | Barramundi calicivirus | 48 | 14 | ≈6000 bp full-length polyprotein | BioProjects submitted from China except PRJNA738523 (US); 6 projects with full-length polyprotein |
| Novel | *Caliciviridae* | Zebrafish neurinoma cali-civirus | Barramundi calicivirus | 1 | 1 | ≈6000 bp full-length polyprotein | BioProject submitted from US (Mayo Clinic) |
| Novel | *Caliciviridae* | Zebrafish systemic cali-civirus | Atlantic salmon cali-civirus | 1 | 1 | 211 aa polyprotein |  |
| Novel | *Hantaviridae* | Zebrafish cardiovascular hantavirus | Wenling red spikefish hantavirus | 6 | 1 | 202 aa L protein |  |
| Novel | *Hantaviridae* | Zebrafish gonadal han-tavirus | Murray-Darling rainbow-fish hantavirus | 2 | 1 | 31 5 aa L protein |  |
| Novel | *Hantaviridae* | Zebrafish neural han-tavirus | Murray-Darling rainbow-fish hantavirus | 1 | 1 | 444 aa L protein |  |
| Novel | *Nanhypoviridae* | Zebrafish arterivirus | Wuhan japanese half-beak arterivirus | 4 | 1 | 2200 aa ORF1ab | Study conducted in Spain (brain tissue); Submission from US |
| Novel | *Orthomyxoviridae* | Zebrafish influenza B-like virus | Betainfluenza & Wuhan spiny eel influenza virus | 80 | 13 | All 8 segments | BioProjects submitted from China except PRJEB2368 (UK) |
| Novel | *Orthomyxoviridae* | Zebrafish influenza-like virus | Siamese algae-eater influenza-like virus | 14 | 4 | 4 of 8 segments |  |
| Novel | *Paramyxoviridae* | Zebrafish hepatic paramyx-ovirus | Salmon aquaparamyx-ovirus | 1 | 1 | 2200 aa L protein |  |
| Novel | *Paramyxoviridae* | Zebrafish systemic paramyxovirus | Wenzhou pacific spadenose shark paramyxovirus | 4 | 2 | ≈810 aa L protein |  |
| Novel | *Picornaviridae* | Danio blood picor-navirus/Zebrafish picor-navirus 2 | — | 1369 | 320 | Full-length polypro-tein | Partially characterized; two clades ≈7% divergent (68 full-length transcripts) |
| Novel | *Picornaviridae* | Zebrafish jaw picornavirus | Coleura bat picor-navirus | 89 | 37 | ≈7500 bp full-length polyprotein | 10 BioProjects with full-length polyprotein |
| Novel | *Poxviridae* | Zebrafish jaw poxvirus | Carp Edema Virus | 4680 | 143 | Much of genome, for phylogeny us-ing rpo147+rpo132 (7489 bp) | Extremely low diversity in phylo-genetic analysis |
| Novel | (Adomaviruses) | Zebrafish adomavirus | Symphysodon discus adomavirus 1 | 1 | 1 | 205 aa L06 |  |
| Novel | (Rhabdo-like) | Zebrafish rhabdo-like virus | Salmovirus WFRC1 | 37 | 9 | 849 aa polyprotein with RDRP; 519 aa protease | BioProjects submitted from Japan, Portugal, Spain, UK |
| Known | *Birnaviridae* | Rocky Mountain birnavirus | — | 19 | 7 | 900 aa & 1000 aa (2 segments) |  |
| Known | *Caliciviridae* | Myotis ricketti associated fish calicivirus | — | 8 | 1 | 650 aa polyprotein |  |
| Known | *Flaviviridae* | Wenzhou shark flavivirus | — | 18 | 1 | 1290 aa polypro-tein |  |
| Known | *Iridoviridae* | Largemouth bass virus | — | 151 | 4 | ≈2200 bp | aka Ranavirus micropterus1 |
| Known | *Iridoviridae* | Singapore grouper virus | — | 38 | 2 | ≈2200 bp | aka Singapore grouper iridovirus aka Ranavirus epinephelus1 |
| Known | *Orthomyxoviridae* | Infectious salmon anemia virus | — | 19 | 1 | All 8 segments | aka Isavirus salaris |
| Known | *Picornaviridae* | Zebrafish picornavirus 1 | — | 1161 | 220 | Full-length polypro-tein | Low diversity; one clade (32 full-length transcripts); aka Cyprivirus A |
| Known | *Spinareoviridae* | Grass carp reovirus | — | 14 | 1 | 1400 bp | aka Aquareovirus ctenopharyn-godontis |
| Known | *Alloherpesviridae* | Cyprinid herpesvirus 3 | — | 186 | 4 | 11,000 bp | aka Cyvirus cyprinidallo3; **inten-tionally administered** |
| Known | *Nodaviridae* | Redspotted grouper ner-vous necrosis virus | — | 4 | 2 | 3000 bp | aka Betanodavirus epinepheli; **intentionally administered** |
| Known | *Rhabdoviridae* | Spring viraemia of carp virus | — | 1839 | 13 | Full-length polypro-tein | aka Sprivivirus cyprinus; **inten-tionally administered** |
| Endogenous/Not fish-associated | (Adomaviruses) | Leatherback sea turtle adomavirus | — | 2 | 2 | — | Evidence of endogenous or **not fish-associated** |
| Endogenous/Not fish-associated | *Astroviridae* | Flumine Astrovirus 3 | — | 6 | 2 | — | Evidence of endogenous or **not fish-associated** |
| Endogenous/Not fish-associated | *Orthomyxoviridae* | Deltainfluenzavirus | — | 7 | 1 | — | Evidence of endogenous or **not fish-associated** |
| Endogenous/Not fish-associated | *Parvoviridae* | Psittaciform aveparvovirus | — | 19 | 12 | — | Evidence of endogenous or **not fish-associated** |
| Endogenous/Not fish-associated | *Parvoviridae* | Psittacidae aveparvovirus | — | 18 | 10 | — | Evidence of endogenous or **not fish-associated** |
| Endogenous/Not fish-associated | *Parvoviridae* | Luscinia sibilans parvo-like hybrid virus | — | 8 | 5 | — | Evidence of endogenous or **not fish-associated** |
| Endogenous/Not fish-associated | *Parvoviridae* | Phoenicopteridae parvo-like hybrid virus | — | 3 | 3 | — | Evidence of endogenous or **not fish-associated** |
| Endogenous/Not fish-associated | *Parvoviridae* | Phoenicopterus roseus parvo-like hybrid virus | — | 3 | 2 | — | Evidence of endogenous or **not fish-associated** |
| Endogenous/Not fish-associated | *Parvoviridae* | Fish-associated parvo-like hybrid virus | — | 35 | 19 | — | Evidence of endogenous or **not fish-associated** |
| Endogenous/Not fish-associated | (Picorna-like) | Fish-associated picorna-like virus 3 | — | 12 | 3 | — | Evidence of endogenous or **not fish-associated** |
| Endogenous/Not fish-associated | *Adintoviridae* | Adintovirus anguilla10665 | — | 34 | 19 | — | Evidence of **endogenous** or not fish-associated |
| Endogenous/Not fish-associated | *Adintoviridae* | Pimephales minnow adin-tovirus | — | 168 | 104 | — | Evidence of **endogenous** or not fish-associated |
| Endogenous/Not fish-associated | *Chuviridae* | Sea turtle neural virus 1 | — | 8 | 8 | — | Evidence of **endogenous** or not fish-associated |
| Endogenous/Not fish-associated | *Chuviridae* | Chuvivirus | — | 116 | 110 | — | Evidence of **endogenous** or not fish-associated |
| Endogenous/Not fish-associated | *Chuviridae* | African cichlid piscichu-virus | — | 151 | 125 | — | Evidence of **endogenous** or not fish-associated |
| Endogenous/Not fish-associated | *Chuviridae* | Hardyhead chuvirus | — | 1 | 1 | — | Evidence of **endogenous** or not fish-associated |
| Endogenous/Not fish-associated | *Chuviridae* | Salarius guttatus piscichu-virus | — | 67 | 58 | — | Evidence of **endogenous** or not fish-associated |
| Insufficient evidence | *Adintoviridae* | Astyanax tetra cavefish adintovirus | — | 22 | 15 | 600 bp |  |
| Insufficient evidence | *Adintoviridae* | Larimichthys croaker adin-tovirus | — | 19 | 14 | 520 bp |  |
| Insufficient evidence | (Astro-like) | Guangdong catfish astro-like virus | — | 1 | 1 | 370 bp |  |
| Insufficient evidence | *Alloherpesviridae* | Cyprinid herpesvirus 1 | — | 1 | 1 | 240 bp |  |
| Insufficient evidence | *Birnaviridae* | Blotched snakehead virus | — | 1 | 1 | 300 bp |  |
| Insufficient evidence | *Hepeviridae* | Cutthroat trout virus | — | 1 | 1 | 570 bp |  |
| Insufficient evidence | *Retroviridae* | Snakehead retrovirus | — | 1 | 1 | 250 bp |  |
| Insufficient evidence | (Adomaviruses) | Blueface angelfish ado-mavirus | — | 1 | 1 | 280 bp |  |
| Insufficient evidence | (Adomaviruses) | Catfish adomavirus | — | 1 | 1 | 200 bp |  |

Total entries: 53 (16 Novel, 11 Known, 17 Endogenous/Not fish-associated, 9 Insufficient evidence)

Phylogenies revealed diverse evolutionary histories among zebrafish virus genomes (**Supplemental Figure S8**). Strikingly, one of the two newly identified *Orthomyxoviridae* family viruses showed discordance in segment trees where some segments grouped closely with human influenza B virus (IBV) while others grouped with influenza viruses of fish (**Figure 4B, Supplemental Figure S8**). We provisionally named this virus Zebrafish influenza B-like virus (ZIBV). Similar to IBV, the ZIBV genome is made up of eight single-stranded negative-sense RNA segments. Segments encoding the ZIBV viral polymerase and proteins that function within cells were highly similar to IBV proteins, with three proteins exceeding 90% amino acid identity. In contrast, segments encoding proteins that facilitate entry and transmission between cells showed a different pattern. Phylogenies of the viral surface glycoproteins hemagglutinin (HA) and neuraminidase (NA) grouped ZIBV and Wuhan spiny eel influenza virus (WSEIV) [42] as sister taxa (**Figure 4B**). We also observed distinct sequence similarities for the two proteins encoded on genomic segment 7, with high conservation between ZIBV and IBV structural protein M1 but high divergence between ZIBV and IBV viral envelope-spanning ion-channel protein M2. These patterns indicate a complex evolutionary history for ZIBV and highlight an influenza-like virus from zebrafish as the closest known relative of a significant human pathogen.

We identified a variety of tissue and cell type tropisms among zebrafish viruses (**Figure 4C,D; Supplemental Figure S9**). ZIBV was detected at high levels in the swim bladder, which is evolutionarily related to human lungs [43, 44] and can be experimentally infected by human influenza A virus [45]. Three of the most prevalent viruses were picornaviruses including Zebrafish picornavirus 1 (ZfPV), which was most abundant in the digestive system as previously reported [46, 47]. Sequences from another picornavirus closely matched GenBank accession OK619588, a partial viral genome labeled as Zebrafish picornavirus 2. A *Picornaviridae* phylogeny placed this virus in a large clade of fish-infecting viruses that are very distantly related to the other zebrafish picornaviruses (**Supplemental Figure S8**). This virus was detected in many tissues and was most abundant in cancer-related datasets (**Figure 4C**). We detected the highest viral reads in samples from a leukemia study [48] (**Supplemental Data S1**), indicating an association with blood, and hereafter we refer to this virus as Danio blood picornavirus (DBPV). Additionally, we identified scRNA-seq datasets with high DBPV counts that were processed as pseudo-bulk by the RNAquarium pipeline, which we re-processed to characterize viral tropism at single-cell resolution. Among all cells that were profiled in studies of the heart [49] and pancreas [50], we detected DBPV predominantly in macrophages (**Figure 4D**). Additionally, macrophages were the predominant cell type containing DBPV transcripts in two studies that profiled immune cells from the brain [51] and kidney [52] (**Supplemental Figure S10**). These results suggest that the broad prevalence of DBPV across tissues is driven by infection of a cell type that is ubiquitously distributed throughout the body.

### Linking host transcriptomes to infection biology

Next, we leveraged our paired transcriptome–virus detection dataset to characterize systematic host transcriptional changes associated with viral presence. We focused on the 18 fish-associated viruses detected in at least 10 transcriptomes. For each virus, we quantified per-gene expression shifts using normalized expression values (**Figure 5A**). We then compared expression shifts of 87 immune-related genes spanning five functional categories with the corresponding genome-wide shifts (**Supplemental Figure S11**). We observed contrasting response patterns (**Figure 5B**). Nine viruses (**Figure 5B**, columns 1–9) were associated with a broad innate immune program comprising a shared interferon-associated module (*ifih1, irf3, irf7, isg15, mavs, mxa, pkz, rsad2, stat2, stat4*) together with immune modulators (e.g., *socs3a*), and additionally showed robust induction of complement components (*c3a.1, c3a.2, c3a.3, c3a.6, c5, c9*) and complement factor b (*cfb*). Five viruses (**Figure 5B**, columns 11–15) induced the same core interferon-associated module but lacked the complement signature, consistent with a more restricted intracellular antiviral defense response. Of note, Grass carp reovirus (**Figure 5B**, column 10) elicited the most extensive transcriptional response, encompassing the shared interferon module together with activation of additional innate sensing and inflammatory pathways, including cGAS–STING/TBK1 signaling (*sting1, tbk1*), Toll-like and NOD-like receptors (*nlrc3, nlrp3, nod2, tlr2, tlr3*), downstream regulators (*irf4a, irf5, nfkbiaa*), pro-inflammatory cytokine/chemokine genes (*cxcl8a, il1b*) and the chemokine receptor *ccr2* (**Supplemental Figure S11**).

**Figure 5.**
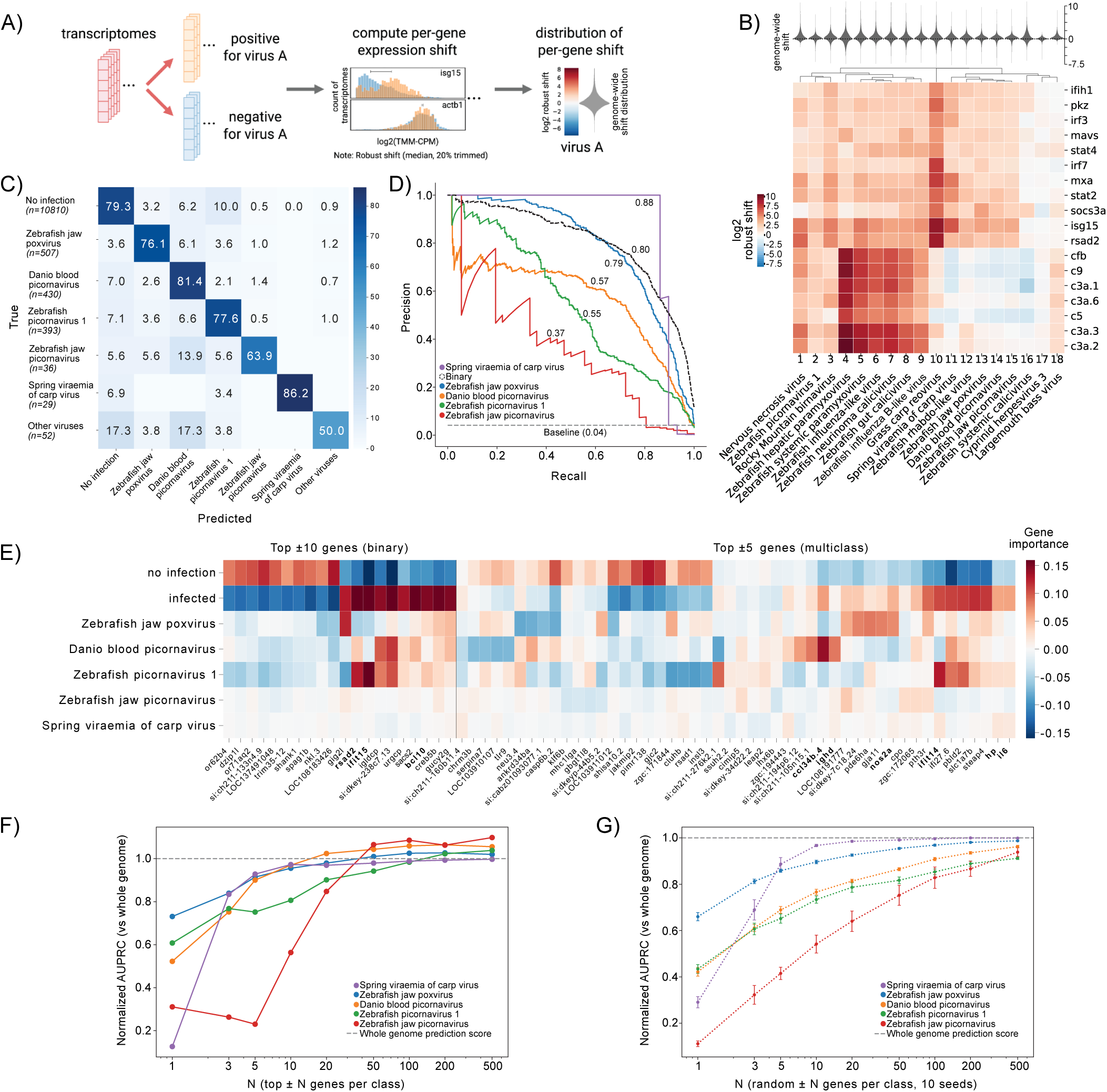
Transcriptomic signatures of virus infections. (A) Computational workflow for quantifying per-gene expression shifts between virus-positive and virus-negative transcriptomes. A robust expression shift was computed for each gene as the difference between the median normalized log-expression [log_2_(TMM-CPM + prior count)] of the virus-positive and virus-negative sets, equivalent to the log_2_ fold change of median expression. A genome-wide distribution of expression shifts per virus serves as the transcriptome-wide background. (B) Heatmap of the log_2_ robust shift for a subset of immune genes (rows, see **Supplemental Figure S11B** for the full panel), grouped by functional category, across 18 viruses (columns), each detected in *≥* 10 transcriptomes. Rows and columns are hierarchically clustered. Violin plots above each column show the genome-wide shift distribution for each virus, providing the transcriptome-wide baseline against which the immune-gene shifts can be read. (C) Confusion matrix showing the row-normalized performance of a Support Vector Machine (SVM) classifier for predicting the dominant virus given a transcriptome. The “others” category is an aggregation of eight distinct virus taxa with low sample representation (*n <* 100 per class). Row totals in the matrix that do not sum to 100% reflect minor scoring cases that were removed from the off-diagonal without being reclassified as primary hits. A comprehensive confusion matrix including all model categories and misclassification instances is provided in **Supplemental Figure S11D,E**. (D) Precision-Recall Curve (PRC) of the five most prevalent viral classes. The PRC for a binary “infected vs. uninfected” classifier is indicated by the black dotted line. The gray dashed line represents the baseline performance based on the maximum frequency class. (E) Gene importance heatmap derived from linear SVM coefficients for the top 10 positive and negative gene features contributing to the binary classification as well as top five positive and negative genes contributing to the multiclass classification of the five most prevalent viruses. (F) Model performance across varying numbers of genes with the highest importance values. Classification AUPRC is plotted as a function of combining the number of highest ranking genes (N = 1 to 500) from linear SVM per class. All scores are normalized against the AUPRC achieved using the entire genome (indicated by the horizontal dashed line at 1.0). (G) Performance of randomly selected gene sets of varying size.

To control for atlas composition, we recomputed each virus’s expression shifts within matched developmental-stage and tissue strata (direct standardization; see STAR Methods), testing whether the complement signature of the broad-innate viruses persists once stage and tissue are held constant. After matching, a complement signal remained detectable (95% bootstrap CI excluding zero) in eight of nine broad-innate viruses, as did the interferon module (positive control: genuinely virus-induced genes expected to persist after matching). The complement shift, however, was attenuated in magnitude, indicating that a fraction of the apparent complement elevation is attributable to sample composition (**Supplemental Figure S12**). For Zebrafish hepatic paramyxovirus, whose positive samples derive entirely from adult liver, the signature was abolished (**Supplemental Figure S12**), consistent with a compositional rather than an infection-associated origin. The finding was robust to finer stratification and to a stage/tissue-adjusted linear model; one broad-innate virus (Zebrafish influenza-like virus) was excluded owing to insufficient stratum coverage. These results indicate that the complement association with the broad-innate viruses is genuine albeit partly compositional: its magnitude is inflated by, and for at least one virus fully attributable to, the developmental-stage and tissue distribution of the virus-positive samples.

To determine whether these distinct host transcriptional signatures are robust enough to predict the existence of virus transcript as well as the viral identity, we developed a Support Vector Machine (SVM) classifier with a Radial Basis Function (RBF) kernel utilizing L2 regularization (see STAR Methods). We designed two distinct classification tasks based on host gene expression profiles: a binary classifier to distinguish between virus-associated and non-virus-associated zebrafish transcriptomes, and a multiclass classifier to predict the specific virus association. For transcriptomes containing multiple viruses, the multiclass label was assigned based on the most abundant viral transcript present. The assignment of virus class can be found in the UMAP plot of **Supplemental Figure S11** with the abundance of each group in **Supplemental Table S2**. Models were evaluated using a 20% hold-out test set. The binary classifier achieved over 85% accuracy in distinguishing infected from uninfected samples (**Supplemental Figure S11**). For the multiclass task, the confusion matrix (**Figure 5C; Supplemental Figure S11**) demonstrates the model’s capacity to correctly assign specific viral classes, with the overall accuracy exceeding 70% for the major classes. A substantial portion of misclassifications occurred in samples with co-infections, where the predicted secondary virus was present but not designated as the ‘major’ class based on transcript counts. Furthermore, the Precision-Recall (PR) curves for the five most abundant viruses also demonstrate strong classification performance, with each viral class tracking well above the baseline (**Figure 5D**).

To interpret the biological features driving these predictions, we extracted the gene weights from the trained SVM models (**Figure 5E**). We visualized the top 10 most positive and negative gene weights for the binary classifier, alongside the top five genes for each of the five most abundant viral classes (the multiclass categories). In this context, positive weights indicate that higher gene expression drives the prediction towards a specific class, whereas negative weights push the prediction away. For the binary classifier (**Figure 5E, left**), the features most strongly driving the ‘infected’ prediction were dominated by canonical antiviral and immune-signaling genes. Notably, the classifier heavily weighted known interferon-stimulated genes, including *ifit15* and *rsad2*, alongside the NF-*κ*B signaling regulator *bcl10*, to positively define the infected state. This confirms that the binary SVM successfully anchors its predictions on the shared, core vertebrate antiviral defense program.

Next we compared gene expression levels between datasets that the model classified as infected to datasets classified as uninfected (**Supplemental Figure S11**). This analysis demonstrated that the differences between infected and uninfected transcriptomes are dominated by the core antiviral signature driving the binary classification. We observed a robust, universal upregulation of canonical interferon-stimulated genes, such as *isg15*, *rsad2*, and *stat1b*, across all viral classes. This alignment confirms that the high classification accuracy of the binary model is rooted in a consistent antiviral host response. Because this dominant signature is shared ubiquitously across infections, it is uninformative for distinguishing between infections by different virus species. Consequently, the multiclass classification task must look beyond this universal immune baseline, forcing the model to isolate distinct gene expression patterns tailored to individual viruses.

While binary classification was dominated by high-magnitude gene-weight profiles, multiclass classification captured a more nuanced landscape of lower-magnitude, virus-specific feature peaks (**Figure 5E, right**), likely reflecting distinct modes of virus-host interaction. To biologically interpret these signatures, we focus on the top-weighted features that possess well-characterized functional annotations. For example, the classifier for Danio blood picornavirus is strongly informed by a coordinated systemic immune profile; this includes the robust induction of the chemokine *ccl34b.4* alongside the B-cell marker *ighd* (immunoglobulin heavy constant delta), consistent with a blood-associated nature of infection. The classification of Zebrafish jaw poxvirus relies on a combination of localized immune defense and epithelial remodeling, featuring high weights for antiviral gene *nos2a* (inducible nitric oxide synthase) alongside the gap junction marker *gja1*, which likely reflects the host cellular response to poxvirus-induced epidermal lesions. In contrast, the classification for Zebrafish picornavirus 1 relies on a distinct set of systemic inflammation and innate immunity responses, showing strong positive weights for *il6* and *hp* (haptoglobin), and the interferon stimulated gene *ifit14*. Zebrafish jaw picornavirus and Spring viraemia of carp virus exhibited comparatively weaker weight magnitudes across their feature sets. This attenuation is a direct consequence of the L2 regularization applied during training, though for opposing reasons: Spring viraemia of carp virus is highly separable, achieving the highest Area Under the Precision-Recall Curve (AUPRC, **Figure 5D**) and thereby requiring smaller weights to achieve accurate classification, whereas Zebrafish jaw picornavirus presents a highly difficult classification task (with lowest AUPRC, **Figure 5D**), preventing the model from assigning large, confident weights to individual genes.

Finally, we sought to determine whether the top SVM-weighted genes encode uniquely informative signatures of virus association, or whether similar predictive information is distributed across the broader transcriptomic landscape. By retraining the model using the gene set which combines the top N most positively and negatively weighted genes per class, we tracked the resulting AUPRC scores normalized against the performance of the whole-transcriptome model (**Figure 5F**). Notably, selecting just the top 50 to 100 genes per class achieved classification performance comparable or even higher than the models utilizing the entire transcriptome (detailed precision-recall curves for each class at varying values of N are provided in **Supplemental Figure S11**). To rigorously test the specificity of these markers, we compared them against models trained on equivalent numbers of randomly selected genes. Crucially, these random sets were extracted from the whole genome after excluding the 50 most positive and negative genes for each class to ensure no overlap with the primary predictive features. While these “constrained” random models yielded markedly inferior and more variable performance compared to the SVM-weighted sets, they still achieved relatively high AUPRC scores as the number of features increased (**Figure 5G**). This phenomenon suggests that while specific “marker” genes provide the most focused predictive signal, biological information regarding the virus association is distributed across the general transcriptomic landscape via correlated gene networks and interconnected signaling pathways. This “wisdom of crowds” effect indicates that viral infection triggers broad, systemic shifts in the host transcriptome rather than isolated changes in a few discrete genes. Similar phenomena regarding the high predictive potential of distributed gene patterns and the “wisdom of crowds” in transcriptomic data have been reported previously under different biological contexts [53, 54].

### RNAquarium portal

To facilitate access and exploration of the reprocessed zebrafish transcriptome collection and its associated microbial and viral content, we built the RNAquarium portal at portal.rnaquarium.org (**Figure 6)**. Users can explore zebrafish transcriptomes and gene expression and/or microbial association patterns using coordinated linked UMAP views. Taxon- and metadata-based interfaces support browser-based investigation of microbial diversity, including fish-associated viral transcripts, as well as sample- and contig-level metadata. An integrated AI chatbot allows querying our entire data resource using natural language. Researchers can ask questions that require code execution to complete analyses that extend beyond the portal’s pre-programmed views.

**Figure 6.**
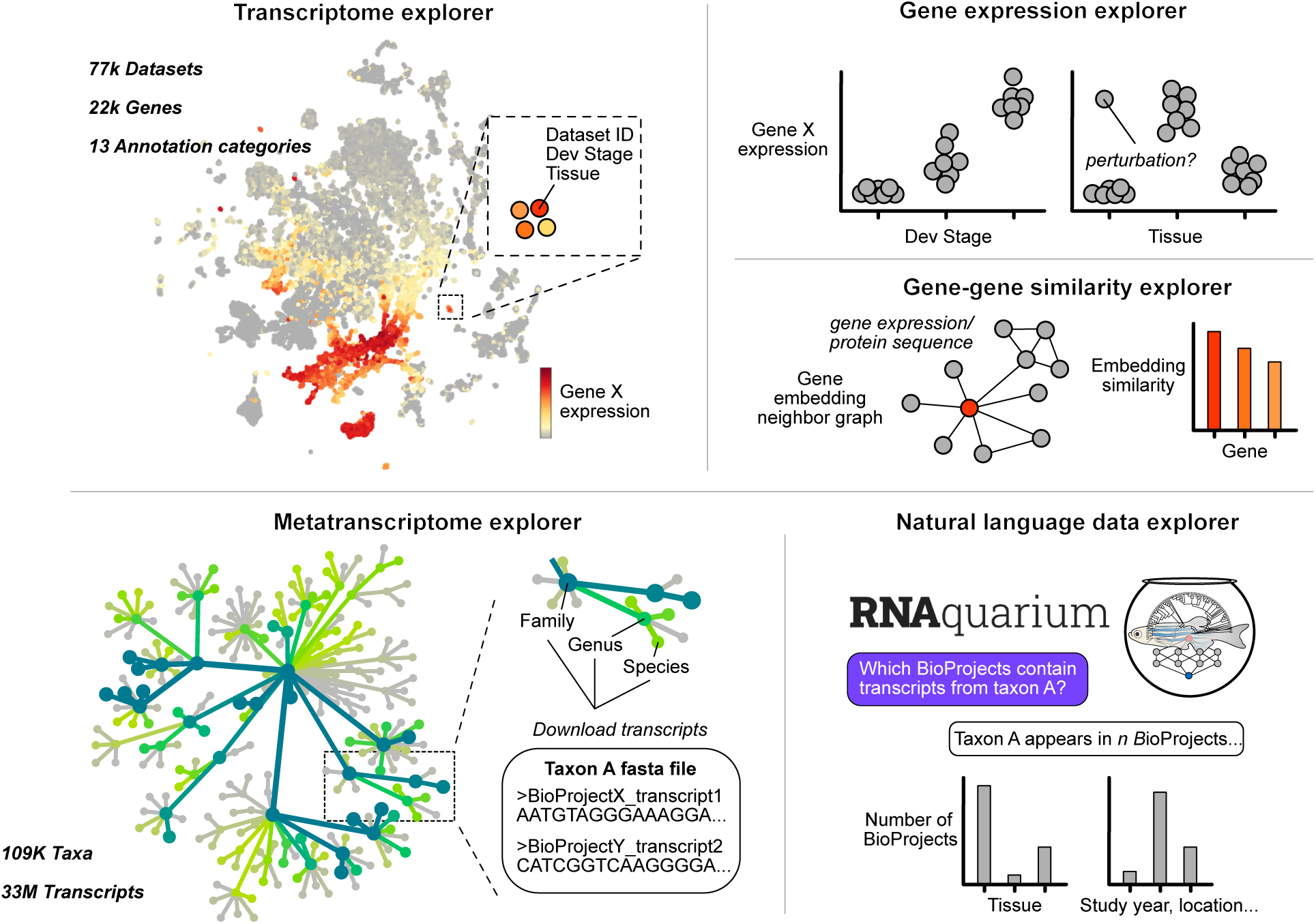
The RNAquarium website. Overview of resources and tools for exploring the RNAquarium atlases of zebrafish transcriptomes and metatranscriptomes at https://portal.rnaquarium.org/.

## DISCUSSION

This study quantified zebrafish gene expression and annotated non-zebrafish sequences across 77,188 RNA-seq runs to generate a multi-scale atlas of transcriptomes and biological communities associated with a widely used vertebrate model organism.

The RNAquarium zebrafish atlas presented here complements several important gene-related resources. ZFIN provides invaluable curation of gene annotations based on published zebrafish research using approaches to infer wild-type expression patterns in standard environmental conditions [19] for 14,472 genes at the time of this study. Our atlas provides quantitative contextual annotations for 22,252 genes across approximately 75,000 observations and illuminates diverse associated taxa as potential environmental contributors to gene expression even in standard zebrafish laboratory conditions. We annotated most datasets with ZFIN ontologies for life stage and anatomy [18] to harmonize gene queries across resources. In addition to revealing gene expression patterns across ages and tissues, we generated gene representations that captured gene-gene relationships and cellular-scale ontologies demonstrating potential value to single-cell applications. The RNAquarium zebrafish atlas complements a growing collection of whole-body zebrafish scRNA-seq atlases [33, 55–61] by leveraging deeper sampling of transcriptomes and an extremely diverse set of experimental conditions that were intentionally and incidentally profiled across all life stages. Combining these atlases could yield cell-type estimates through deconvolution [62] of the billions of cells that were collectively sampled using bulk RNA-seq in the thousands of zebrafish studies summarized here. We anticipate that the RNAquarium zebrafish atlas will facilitate marker gene discovery and the generation of new tools for visualizing and perturbing cell types and states across scales in living animals.

The RQ-GF foundation model demonstrates that compendium-scale bulk RNA-seq, when sufficiently diverse, can serve as a substrate for learning gene representations that generalize to single-cell. We note that most scRNA-seq accessions in the RNAquarium were pseudo-bulked prior to pretraining, although some sequencing runs derived from individual cells that provided the model with limited exposure to transcriptomes at single-cell resolution. Exhaustive inclusion of all RNA-seq datasets in the archive at single-cell resolution would further enhance the multi-scale representations we constructed with primarily bulk sequencing data. Beyond classification of cells, we found that RQ-GF generated gene embeddings that could be combined through vector arithmetic to assemble cell differentiation trajectories, indicating that the model captured an approximate, compositional grammar of regulatory logic. Building on the model with refinements that account for nonlinearity, dosage, and timing would strengthen the ability to rank transcription factor combinations for directed differentiation or reprogramming of cell types and states in silico.

Fish, humans, and other vertebrates are infected by related viruses [42, 63] and our atlas of zebrafish-associated taxa reveals newly described representatives from diverse viral families in a widely used model vertebrate. We detected prospective zebrafish viruses in approximately 10% of all RNA-seq accessions, with the most prevalent viruses detected in thousands of accessions spanning the earliest to most recently submitted zebrafish datasets from laboratories around the world. Our atlas provides a unique hypothesis generation tool for exploring archival data and a resource for establishing surveillance strategies to detect, manage, and study viruses in zebrafish colonies. We distinguished zebrafish viruses from viruses infecting other zebrafish-associated organisms or contamination in the atlas based on evidence from phylogenetics, tissue tropism, and antiviral signatures in zebrafish transcriptomes. Contaminants were often classified based on having greater than 90% sequence similarity to a known virus from another organism, with one notable exception being a newly described zebrafish virus that was highly similar to human IBV. There is currently no clear evidence for an established non-human IBV reservoir although reverse zoonosis from humans to other mammals is suspected [64]. The closest known relatives to IBV were recently described in metatranscriptomic studies of amphibians and fish [42, 65, 66], which fall as sister groups in phylogenies of viral polymerase protein PB1 and share between 70–78% amino acid identity with PB1 from IBV. To our knowledge, the 91% amino acid sequence identity between PB1 from a zebrafish virus and IBV is higher than any other fish and mammalian virus sequence comparison reported to date. Possible evolutionary scenarios for the contrasting phylogenetic groupings of influenza B-like proteins include divergent and convergent processes, a host switching event involving reassortment of segments between human and fish viruses, and combinations of evolutionary histories. In addition to revealing unprecedented sequence similarities between the viruses of fish and humans, we anticipate that the atlas of diverse zebrafish viruses described here will provide new opportunities to measure and model shared features of infection biology and uncover novel regulatory and structural modules that can be repurposed as molecular tools for gene expression and cellular engineering.

In RNA-seq studies of zebrafish, laboratory conditions are generally reported only for abiotic variables including temperature and light cycle. Our atlas of zebrafish-associated taxa paired with zebrafish transcriptomes illuminates biotic components of laboratory environments and identifies viral taxa as hidden drivers of zebrafish gene expression patterns. The approaches we took to measure gene expression shifts associated with viral presence and classify infected transcriptomes could be extended to identify significant interactions between zebrafish and other taxonomic groups represented in the atlas, including bacteria, fungi and other eukaryotes, as well as dark matter sequences that lack detectable sequence similarity to taxa in the reference databases we used.

We created the RNAquarium pipeline to process large numbers of RNA-seq datasets for coupling gene expression quantification and taxonomic classification of non-reference sequences. Our solutions for metadata-independent pre-processing, optimization in distributed computing, and extension of research-subject centered transcriptome quantification to pan-taxonomic profiling, illustrate how data processing workflows can be made more robust, scalable, and generalizable to illuminate the complexity of biological systems captured in RNA-seq data. Our study of all zebrafish RNA-seq accessions stored in the SRA as of July 2025 enriches our understanding of one of the most used vertebrate model systems. Furthermore, the pipeline engineering choices and outputs presented here will be valuable for training and evaluating future AI systems designed to automate the processing and analysis of RNA-seq datasets from all research subjects represented in public repositories.

## Limitations of the study

We unified the global research community’s zebrafish transcriptomic data through the lens of a single reference genome, which is a limitation given the genetic diversity that exists among laboratory zebrafish [67–70]. Emerging zebrafish pangenome resources [12] will provide more inclusive reference mapping sets, and alternative approaches that assemble and annotate all sampled transcripts in the input data could enable exhaustive representations of transcriptomes. We note that a wide range of documented biases inherent to RNA-seq data generation [71, 72] likely influence the transcriptomic and metatranscriptomic atlases presented here, one example being that bacteria will be underrepresented in datasets generated with enrichment of polyadenylated transcripts [73]. We aggregated transcriptomes from bulk RNA-seq and pseudo-bulked scRNA-seq datasets. Our single-cell analysis of a small fraction of the scRNA-seq datasets indicates that a systematic survey would enhance a multi-scale representation of zebrafish gene expression and associated taxa, which could be accomplished through further improvements to the Seq-Detective tool we developed for this study. Other engineering and processing challenges to consider in future large-scale studies include further optimization of workflow management structures that minimize the significant bottlenecks of disk-intensive processes, refinement of dynamic resource scaling strategies, and more comprehensive parsing and processing of technical and biological sequences that overcomes imperfect or absent metadata in public sequencing repositories. Beyond zebrafish, this study establishes a framework for leveraging community-generated RNA-seq data to jointly investigate transcriptomes and metatranscriptomes across the full diversity of life represented in public repositories.

## RESOURCE AVAILABILITY

### Lead contact

Requests for further information and resources should be directed to and will be fulfilled by the lead contact, Keir Balla.

### Materials availability

This study did not generate new materials.

### Data and code availability

- All generated data is available on the RNAquarium portal at portal.rnaquarium.org, publicly available as of the date of publication.
- All original code is available on GitHub at github.com/czbiohub-sf/RNAquarium and github.com/czbiohub-sf/seq-tech-detective.
- Code to generate tables and figures is at github.com/czbiohub-sf/rnaquarium-figures.
- Sequences of 15 novel zebrafish viruses (all but zebrafish jaw poxvirus) have been submitted to GenBank as TPA sequences XXXXXXXX - XXXXXXXX.
- Any additional information required to reanalyze the data reported in this paper is available from the lead contact upon request.

## ACKNOWLEDGMENTS

We thank the zebrafish research community for generating and sharing data used in this study, and the International Nucleotide Sequence Database Collaboration for making it publicly accessible. We thank Alejandro Granados, Deepika Sundarraman, Fitzgerald Small, and Anthony Cu for contributions to early versions of analyses, and Sandy Schmid and Joe Derisi for helpful discussion and feedback on the manuscript.

This work was funded by Biohub.

## AUTHOR CONTRIBUTIONS

Conceptualization, D.P. and K.B.; methodology and investigation, all authors; writing-—original draft, Y.A., E.W., L.L., H.H., Y.S., D.P., and K.B.; writing-—review & editing, all authors.

## SUPPLEMENTAL INFORMATION INDEX

Figures S1-S16, tables S1-S4 and their legends in a PDF.

Data S1. Virus counts for 53 putative fish-associated viruses across all SRA accessions, related to Supplemental Figure S1 and Figure 2.

Data S2. Set of 150 SRA accessions used for benchmarking Seq-Detective, related to STAR Methods.

Data S3. All 77,329 transcriptomic zebrafish SRA accessions found, searched on 22 June 2025, related to STAR Methods.

## Declaration of generative AI and AI-assisted technologies in the manuscript preparation process

During the preparation of this work the authors used Claude(Anthropic) and ChatGPT(OpenAI) in order to perform literature search, draft portions of the methods, and edit the text. Coding agents (Claude Code, Codex, and Cursor) were used to develop the web portal, to prototype analyses and create pipeline documentation for GitHub. After using these tools, the authors reviewed and edited the content as needed and take full responsibility for the content of the published article.

## STAR METHODS

## 1 Overview of RNAquarium pipeline

The RNAquarium pipeline begins by specifying a target species and its reference genome. Very broadly the pipeline can be divided into two workflows: A Transcriptomic + Filtering portion followed by a Metatranscriptomic portion (**Figure 1A; Supplemental Figure S1**). Outputs of Transcriptomic + Filtering are a matrix of host species gene counts, host species sequence data in CRAM format and a large set of unmapped sequencing reads. The Metatranscriptome portion of the pipeline outputs summaries of the taxonomic findings of all non-host transcripts and optionally extracts transcripts of curated taxa of interest, for example viral transcripts, and performs additional clustering and quasi-mapping of all unmapped sequencing reads to a curated set of viral transcripts to create a matrix of curated taxon counts. The entire Transcriptomic + Filtering portion of RNAquarium is wrapped into a Nextflow pipeline [74], and much of the Metatranscriptome portion is similarly wrapped into a second Nextflow pipeline to better accommodate compendium-scale analysis. The final steps of the Metatranscriptome portion including all curated taxon-specific steps are divided into Slurm scripts for running on a local High Performance Computing (HPC) cluster [75].

### 1.1 RNAquarium pipeline – Seq-Detective & Transcriptomic + Filtering

The pipeline is implemented as a Nextflow DSL2 workflow and processes accessions concurrently (set at 50 by default and staggered for rate limits). For each SRA accession, reads are retrieved in full with the SRA Toolkit (prefetch + fasterq-dump --split-3; on download failure, fastq-dump is used as a fallback); a 200,000-read subsample is then drawn with seqtk sample and passed to Seq-Detective (described below) before any quality filtering, with only the B-labeled mate file(s) advancing. Quality filtering is then applied with fastp (Phred quality threshold 17, up to 15% low-quality bases tolerated per read, minimum length 2 bp, polyX trimming, adapter trimming against a custom adapter list). Runs for which Seq-Detective filtering rejected all mate files as indistinguishably technical/low-quality/non-transcriptomic are recorded as dropouts and excluded from further processing. Local FASTQ inputs bypass the download and Seq-Detective steps and enter the quality-filtering stage directly after a simple read-length barcode filter.

After quality filtering, reads are routed in parallel through two branches. The reference mapping branch aligns each run to GRCz12tu (NCBI RefSeq GCF_049306965.1) with STAR 1 (--outFilterMultimapNmax 20, --outFilterMismatchNoverLmax 0.04, splice-junction overhang 1 8/1, intron range 20–1,000,000 bp); per-gene read counts are then computed with featureCounts against the GRCz12tu annotation, and an optional CRAM alignment file is written for storage. The host-filtering branch applies a sequential aligner gauntlet to remove host-mapping reads: first, kallisto in bulk mode against a contaminant index combining the zebrafish RefSeq transcript sequences and common plasmid sequences (AddGene plasmid repository) removes reads matching known molecular biology contaminants; the surviving reads are then passed through HISAT2, STAR, and bowtie2, each indexed against GRCz12tu with ERCC92 spike-in sequences appended; optical and PCR duplicates are then collapsed with czid-dedup (exact matching at full read length); and a final GSNAP alignment filter pass. For paired-end runs throughout the gauntlet, discordant pair alignments (pairs where one mate maps and one does not) are optionally retained as non-mapping candidates, an option we enabled in this study for increased sensitivity at the expense of falsely retaining more host species reads from problematic libraries. Reads surviving all six stages are the unmapped candidates forwarded to the downstream metatranscriptomic assembly and BLAST taxonomic identification workflow.

The first step of the pipeline downloads and pre-processes each sequencing run accession to filter out technical sequences, which we now describe in detail. As previously alluded to, näıve mass processing of SRA transcriptomic datasets will encounter submissions marked as paired end data where the actual library layout is not actually biological cDNA reads, such as barcoded scRNA-seq (T-B or B-T), that silently produce near-zero alignment rates in one mate (**Supplemental Figure S13**), which previous zebrafish reprocessing projects either ignored [4] or noticed but did not diagnose [71]. We first considered whether a simple read length heuristic could solve this problem. For example, in 10x single-cell the informative barcode technical read length is 26 bp (28 bp in v3) and the biological read length usually exceeds 50 bp. However, we observed that many 10x single-cell RNA-seq submissions sequenced both the barcode and cDNA reads at the same length, often choosing a 2 x 75 bp or 2 x 150 bp layout, and a simple length heuristic fails in this case (**Supplemental Figure S14**). Furthermore, while 10x is by far the most popular single-cell sequencing technology in this dataset by read volume, it is not the only type of library that poses an issue, and other barcoded single-cell methods require other length thresholds. A more subtle case is the DeTCT library layout, in which rather than fully T or B, one mate has technical sequence (consisting of 12 random bases, an 8 bp sample tag, and 14 bp poly-T) [76] interleaved with biological sequence in the same read.

We developed Seq-Detective to act as an early filter for bad mates as well as other non-transcriptomic submissions such as species label swaps and immunoprecipitation-based protocols by quickly mapping a subset of downloaded reads to the presumed host, and classifying each file as either true biological (B) reads from that host genome or technical/non-mapping reads. Seq-Detective’s core classification mode processes each FASTQ file in an accession independently. For each file, a sample of up to 200,000 downloaded reads is first passed through fastp (quality threshold Q17, minimum length 2 bp, polyX trimming enabled) to warm the OS file cache and collect pre-alignment QC statistics, then aligned in single-end mode to a HISAT2 [77] index of the host genome (-k 1, no temporary splice sites). Aligned reads are quantified against the host gene annotation with featureCounts [78], yielding per-file estimates of mapping rate, no-feature rate (reads aligning to the genome but outside annotated features), mapping quality-filtered rate, gene count sparsity, and positive-strand ratio. These metrics are serialized to JSON and consumed by the judge module (judge.py -b), which applies a two-stage decision tree to assign each file a B (biological) or T (technical) label. The first stage handles unambiguous cases by read length alone: any file whose reads exceed 500 bp is flagged T (outside the range of the short-read aligner), and for paired submissions any mate whose reads are *≤* 29 bp while the other mate’s reads are *≥* 49 bp is immediately labeled T (barcode-length) with its partner labeled B. For pairs that survive the length gate, the second stage computes a signed percent-difference in mapping rate between the two mates (mapping_diff = (r1 - r2) / max(r1, r2)): if both mates are below 9% mapping rate the run is rejected as T/T; if mapping_diff exceeds *±*0.25, the lower-mapping mate’s no-feature rate is compared against a linear threshold (nofeature_rate *> ±*0.6*×*mapping_diff+0.45) to determine whether it is unambiguously technical or merely weaker but still transcriptomic; remaining pairs whose minimum no-feature rate falls below the envelope *|−*2.8*×*mapping_diff+1.0| are accepted as B/B by a biological fallback assumption, and all remaining pairs assign T to whichever mate has the lower mapping rate. Single-end files use a simpler rule: mapping rate *≥* 1.2% yields B, below that yields T. Specific mapping metric thresholds were tuned by decision tree against 60k human-curated expected decisions set for *Danio rerio* reference GRCz12tu [79] and may need adjustment for other species and/or reference genomes. Runs where all files are classified T are flagged as non-transcriptomic or low-quality and excluded from further processing. For runs with mixed B/T classifications, only the B-labeled files are carried forward. Seq-Detective is run prior to quality filtering and host alignment and operates solely on raw downloaded reads, without reference to submitter-provided metadata.

We validated Seq-Detective’s per-file read-type calls against a manually curated benchmark of 150 accessions spanning bulk RNA-seq, 10x Chromium, Smart-seq, CEL-seq, inDrops, and nine additional minority protocols, selecting for diversity in submission format (single FASTQ, paired FASTQ, BAM, CRAM, and mixed archives; **Supplemental Figure S15; Supplemental Data S2**). Seq-Detective achieved 95.3% exact agreement with curated labels overall, with perfect agreement on 10x (n=40) and near-perfect agreement on bulk (95.7%, n=47); the remaining disagreements were concentrated in unusual submission formats (Smart-seq archived as interleaved BAM+CRAM) and DeTCT, whose reads partially interleave biological and barcode sequence within a single file in a way that resists binary B/T classification. The coherence of Seq-Detective outcomes across submission formats is shown in **Supplemental Figure S2**. We see in the UMAP that DeTCT’s partially-technical format does not fully integrate, as expected with Seq-Detective whole-mate classification resolution (**Supplemental Figure S16**).

In the Transcriptomic + Filtering Workflow, reads are first rapidly depleted of host sequences by Kallisto pseudo-alignment against the zebrafish transcriptome, greatly reducing the post-trimming reads. The remaining reads pass through a cascade of four aligners with complementary strengths (HISAT2, STAR, Bowtie2, and GSNAP), each operating only on the unmapped output of the preceding step, ensuring that host-derived reads are not erroneously carried into metagenomic assembly due to the blind spots of any single aligner. Following de-duplication, remaining reads are retained as putative non-host sequence, while mapped reads are quantified against zebrafish gene annotations to yield host expression profiles.

### 1.2 RNAquarium pipeline – Metatranscriptome

Once the Transcriptomic + Filtering portion of the RNAquarium pipeline is complete, the second Nextflow pipeline implements the Metatranscriptome portion. To agnostically search all possible microbial and non-microbial sources of non-host sequence in the RNA-seq datasets, we chose a very conservative search strategy. This portion of the pipeline first *de novo* assembles the sequencing reads unmapped to the host species, after grouping into SRA BioProject, and then conducts both preliminary and primary taxonomic searches on the resulting expressed contigs, which we term transcripts. The primary inputs to the Metatranscriptome portion of the pipeline are the set of sequencing reads unmapped to the host species split by SRA accession, while secondary inputs are a mapping of SRA accessions to BioProjects and in-house versions of NCBI BLAST databases [80]. The initial steps of this portion are the merging of accessions to the SRA BioProject level followed by *de novo* assembly using rnaSPAdes [81], creating sets of expressed contigs, which we term transcripts. The transcripts are shuffled into equal-sized chunks and then passed through a preliminary BLAST step (using the compressed NT database, [82]), searching only for transcripts matching human or *Danio* sequences in the NCBI NT database (with e-value threshold of 1 x 10^-20^) to further filter to non-host transcripts. Remaining transcripts next undergo a thorough search (with e-value thresholds of 0.05) of both the NCBI protein NR database (clustered; March 2025, [83]) using the Diamond algorithm [84] (using Diamond parameters --ultra-sensitive & --top 3) and the NCBI nucleotide NT database (Core nt; May 2025 [85]) using the BLAST algorithm to identify the nearest taxonomic match for each transcript if found in NCBI databases.

The results of the transcript identification steps are raw database summary (flat) files from the Diamond and BLAST searches, which concludes the Nextflow steps. All subsequent RNAquarium pipeline steps are run as individual Slurm scripts for running on a local HPC cluster. The pipeline next separately analyzes the protein and nucleotide results. This includes extracting full taxonomic info for each transcript from the raw taxid field, calculating the LCA (Last Common Ancestor) for each transcript by removing any taxonomic hits less than 95 percent of the maximum bitscore found, and then finding the LCA of the remaining top taxonomic results. Protein and nucleotide results are then merged, with protein database results only being used for transcripts when the maximum protein bitscore B_NR_ is equal or greater than the nucleotide bitscore B_NT_ (null expectation is B_NR_ = 1/3 B_NT_). The analyzed taxonomic summaries are next visualized with both heatmaps and alluvial plots and the results are split into broad taxonomic groups, followed by a step merging transcript sequences into the summary files. A final general processing step calculates accounting metrics across the Metatranscriptomic steps to make a schematic of the pipeline numbers, and additionally creates a set of ‘dark matter’ transcripts which were not matched to any entries in either protein or nucleotide databases.

All subsequent RNAquarium pipeline steps are taxon-specific. With our interest in viruses infecting zebrafish, these scripts are focused on the viral transcripts but can be modified to focus on other taxonomic groups (e.g. Archaea, Bacteria, Fungi). These steps first split the viral transcripts into phage and non-phage categories, and then create virus-specific alluvial plots and heatmaps which include summaries of the most common virus LCAs found. Subsequent virus-specific steps include additional visualizations of non-phage LCAs using a Metacoder ‘heat tree’ [86], and identifying viral transcript clusters and aligning viral clusters using mmseqs2 easy-cluster and Minimap2 respectively [87, 88]. The scripts finally create summary files with the cluster information, heatmap visualizations of the most common clusters, and collections of clusters for follow-up bioinformatic analyses.

An important final step in the RNAquarium pipeline is to create virus counts for all SRA accessions. Virus counts are calculated using the Salmon quasi-mapping algorithm [89] [90], with inputs for quasi-mapping both the unmapped reads from the Transcriptomic + Filtering steps and a curated set of virus transcripts. The output of the quasi-mapping is a 73,955 x 180,189 matrix of virus counts, with rows representing SRA run accessions and columns representing virus transcripts.

### 1.3 Curation of zebrafish RNA-seq metadata

We additionally gathered and curated metadata for all SRA run accessions deposited as of May 2025 (77,329) from two NCBI sources: SRA and the Gene Expression Omnibus (GEO). First, scripts were created that extract metadata from four levels of organization in the SRA database: Run accession, Sample, Experiment and Study (prioritized in that order). After extracting at each level, the metadata is merged at the run level, keeping the level explicitly named in each column. An intermediate output step is a metadata file with every run (rows) with corresponding metadata fields (columns). Notable fields in SRA run metadata include submission center names and Pubmed IDs when provided by submitters.

We next use the raw metadata from these initial columns to create curated calls for select characteristics. We first sample all possible date fields from the metadata to create an earliest date field. We next use scripts to create curated calls of developmental stage, tissue type, and sequencing technology-related information. These scripts separately search for keywords from the SRA metadata columns, using the four levels of organization to prioritize the curation call. For example, a run-specific tissue description takes precedence over any other level. Similarly, keywords found in study-level metadata would only be used when the other levels do not contain parsable information and even then, are used to populate a secondary “coarse” curation column. Finally, we include mapping metrics and read filtering decisions by Seq-Detective Core as used in the RNAquarium pipeline. The final metadata summary file has 77,329 rows (SRA accessions) and 90 columns of metadata (**Supplemental Table S3; Supplemental Data S3**). This SRA accession total includes a small number of runs not processed by the first step of Seq-Detective (77,188 sequencing runs).

### 1.4 Compilation of the zebrafish RNA-seq expression compendium

The host-genome count matrix produced by the RNAquarium Transcriptomic + Filtering pipeline, a sparse matrix of *≈*75,000 SRA run accessions by GRCz12tu gene features, was loaded into an AnnData object and joined with the curated SRA metadata table (Section 1.3) on SRA run accession. Viral transcript counts from the Salmon quasi-mapping step (Section 1.2) were appended as additional sample-level columns. Gene-level annotations (ZFIN identifier, NCBI GeneID, miRBase accession) were extracted from the GRCz12tu gene annotation and added to the feature metadata. Gene symbols were resolved via the ZFIN (Zebrafish Information Network) gene alias table. Gene biotype, genomic length, and Ensembl stable IDs were appended from an Ensembl BioMart export keyed on ZFIN symbol. Quality control filtering removed any sample or gene with fewer than 10 total counts, followed by a sparsity filter excluding samples and genes for which more than 90% of values were zero. Retained counts were normalized using Trimmed Mean of M-values (TMM; rnanorm) and subsequently log transformed for downstream analysis.

### 1.5 Additional RNAquarium pipeline processing to refine non-host results

For our analyses, we used additional processing steps across much of the RNAquarium pipeline to optimize and refine our results after examining preliminary results. Initial results showed that virus hits such as “Carnation yellows virus” and “Porcine bastrovirus” found in the NCBI NT database are not viral but mislabeled adapter and primer sequence [91]. We included additional primer and adapter sequence into a Host Filtering step to remove such sequence from subsequent analyses. Recent results also indicate more similar mislabeled sequence in the databases, notably “Crocidura shantungensis ribovirus 3” and “Brassica yellows virus.” To more thoroughly filter such mislabeled adapter and primer sequence, we have added an additional post-hoc BBDuk filter to our results, masking all such adapters and primers as Ns and then removing any transcripts with *<* 150 bp of sequence after masking.

We also ran iterations of some processing steps to maximize the number of identified transcripts. For example, we initially used NCBI databases and taxonomic names from spring 2025, but then further ran our taxonomic processing steps through a more recent taxonomic name database from summer 2025. This update was necessary in part because NCBI changed the viral top-level taxonomic rank from ‘clade’ to the unique field ‘realm’. Similarly, because Diamond occasionally fails to add the taxid field to hits (*≈*10% of hits), we added a similar processing step that finds these missing taxid values using the raw targetid column. Finally, for calculating the virus counts, we found that small numbers of transcripts classified as Sprivivirus (Spring viraemia of carp virus) contained sequence similar enough to zebrafish genomic sequence that these led to artificially inflated counts. To correct this, we replaced all the Sprivivirus transcripts in the Salmon pseudo-mapping step with two full length Sprivivirus genomes.

### 1.6 RNAquarium pipeline repository

Both portions of the RNAquarium pipeline, including Nextflow and post-Nextflow steps are available on GitHub at github.com/czbiohub-sf/RNAquarium. Code for Seq-Detective as well as scripts to generate the RNAquarium metadata and subsequently create curated columns are available on GitHub at github.com/czbiohub-sf/seq-tech-detective. A full set of inputs and outputs across the steps of the RNAquarium pipeline, the set of software packages, and databases required to run the pipeline are all listed in GitHub documentation.

## 2 AI modeling of archive-wide transcriptomes and analysis of gene representations

### 2.1 Tokenization of transcriptomic data for foundation model training

To prepare transcriptomic data for Geneformer pretraining, we adapted the Geneformer tokenization pipeline to operate on the zebrafish expression compendium produced by RNAquarium. The tokenization procedure converts each transcriptome into a rank-ordered sequence of gene tokens, following the rank value encoding scheme described in Theodoris et al. [32].

Briefly, we computed corpus-specific gene normalization factors using the t-digest streaming algorithm (crick library) to estimate the non-zero median expression of each gene across all transcriptomes in the RNAquarium corpus. For each transcriptome, raw counts were normalized by total counts per sample and scaled by a factor of 10,000 (counts per ten thousand), then divided by the gene-specific non-zero median expression value. This median-scaling procedure deprioritizes ubiquitously highly-expressed housekeeping genes and elevates genes whose expression is more informative of biological state [32]. Genes with zero expression in a given transcriptome were excluded, and the remaining genes were ranked in descending order of their normalized expression values. The resulting rank-ordered gene token sequences were truncated to a maximum length of 2,048 tokens. No CLS or EOS special tokens were appended, consistent with the Geneformer 30M model series convention. For the Zebrahub single-cell atlas [33], we applied identical tokenization using the same corpus-derived gene normalization factors and token dictionary as the RNAquarium data. Cell metadata including developmental stage, zebrafish anatomy ontology class, time point, and fish identity were carried forward as dataset attributes for downstream fine-tuning.

To establish a noise-trained baseline, we generated a random noise version of the RNAquarium count matrix by replacing all gene expression values with random integers drawn uniformly from [0, 100), thereby destroying all biological signal while preserving the matrix dimensions and gene annotations. This randomized matrix was then processed through the same tokenization pipeline (normalization, median-scaling, ranking, and truncation) using the same token dictionary and normalization factors as the original RNAquarium data.

### 2.2 Geneformer pretraining

We pretrained Geneformer models from scratch on transcriptomic data using a 6-layer BERT architecture (BertForMaskedLM) [32]. The model configuration comprised 6 transformer encoder layers, 4 attention heads, a hidden size of 256 dimensions, an intermediate feed-forward size of 512, and ReLU activation. Dropout rates for both attention probabilities and hidden states were set to 0.02. The maximum positional embedding size was 2,048, matching the tokenization truncation length. The vocabulary size was determined by the number of detected protein-coding and miRNA genes in the corpus plus two special tokens (pad and mask), yielding 22,176 tokens for the RNAquarium dictionary.

Pretraining was performed using a masked language modeling objective, where 15% of gene tokens in each transcriptome were randomly masked and the model was trained to predict the identity of the masked genes from the context of the remaining unmasked genes. Training was conducted for 30 epochs with a batch size of 12, a maximum learning rate of 1 *×* 10*^−^*^3^ with a linear learning rate schedule, 10,000 warm-up steps, AdamW optimizer with a weight decay of 0.001, and random seed 42. Training was distributed across 4 GPUs on a single node using DeepSpeed.

Four separate models were pretrained: (1) RQ-GF: pretrained on the 74,275 RNAquarium bulk and pseudo-bulk transcriptomes; (2) Random gaussian noise: pretrained on the random noise version of the RNAquarium data described above; (3) Zebrahub-pretrained: pretrained on 85,213 Zebrahub single-cell transcriptomes; (4) Cross-species transfer: the human Genecorpus-30M pretrained Geneformer model (GF-6L-30M-i2048) [32], used directly for fine-tuning without additional pretraining on zebrafish data. All *de novo* pretrained models (1–3) used identical architecture and training hyperparameters.

### 2.3 Fine-tuning for cell state classification

To prepare the Zebrahub dataset for fine-tuning, we constructed compound labels by concatenating developmental stage and zebrafish anatomy ontology class for each cell (e.g., “larval-2dpf—central nervous system”). To mitigate class imbalance in this compound label space, we removed classes with fewer than 100 cells and downsampled classes with more than 3,000 cells to 3,000, yielding a balanced dataset of 85,213 single-cell transcriptomes spanning 80 compound classes.

Fine-tuning was performed using the Geneformer Classifier module in cell classification mode. Six transformer layers were frozen during fine-tuning, and training hyperparameters were optimized separately for the RNAquarium-pretrained and Zebrahub-pretrained models using Bayesian hyperparameter search (5 trials per configuration). The search space included number of training epochs (up to 5), learning rate (*≈* 4 *×* 10*^−^*^4^ to 1 *×* 10*^−^*^3^), learning rate schedule (linear, cosine, or polynomial), warmup steps (100–2,000), weight decay (0.02–0.30), and random seed. The batch size was fixed at 12. Optimized hyperparameters were selected based on the macro-F1 score on a held-out validation set.

To assess the effect of pretraining data volume on fine-tuning performance, we varied the fraction of Zebrahub data used for fine-tuning from 5% to 95% in 5% increments, holding out the remainder for evaluation. For each fine-tuning fraction, hyperparameters were re-optimized. Models evaluated included: RQ-GF (pretrained on RNAquarium), random gaussian noise baseline, Zebrahub-pretrained, and a cross-species transfer model initialized from the human Genecorpus-30M pretrained weights (GF-6L-30M-i2048) [32]. Performance was measured using macro-F1 score on the held-out test set.

As a simple model baseline, we trained XGBoost classifiers on PCA-reduced representations of the same Zebrahub data. Principal component analysis was applied with 20, 50, and 100 components, and XGBoost hyperparameters were optimized using Ray Tune with Optuna search over 100 trials (ASHA scheduler, max depth 3–15, learning rate 0.001–0.5, subsample 0.5–1.0, colsample bytree 0.5–1.0, L1/L2 regularization, and 50–500 boosting rounds). Six replicates were run per configuration with random train/test split at the same test-size fractions. Performance was evaluated using macro-F1 score.

### 2.4 Gene embedding extraction

Embeddings were extracted from the second-to-last transformer layer (layer index 1), which has been shown to capture rich contextual representations in transformer models. The extraction procedure passes each tokenized transcriptome through the model and averages the hidden state vectors for each gene token across all transcriptomes in which that gene appears, producing a single 256-dimensional embedding vector per gene. A forward batch size of 200 was used.

### 2.5 Gene network construction and functional enrichment analysis

To assess whether RQ-GF gene embeddings capture biologically meaningful gene-gene relationships, we constructed gene networks from embedding similarities and compared them to co-expression-based networks. For the embedding-based network, pairwise cosine similarity was computed between all gene embedding vectors, yielding a gene x gene similarity matrix. For the co-expression baseline, TMM (Trimmed Mean of M-values) normalization was applied to the RNAquarium count matrix, and expression values were transformed to log_2_(TMM-CPM + prior count). Pairwise gene-gene correlations were computed using a robust correlation with light winsorization (quantile 0.001) to reduce the influence of outliers.

Leiden community detection was applied to both the embedding-based and co-expression-based gene networks across a range of resolution parameters (0.1-100) to identify gene modules at multiple scales. For each set of clusters at each resolution, Gene Ontology (GO) enrichment analysis was performed using g:Profiler. The number of unique GO terms recovered was tallied across resolutions and compared between the RQ-GF embedding network, the co-expression network, and the noise-trained baseline to evaluate functional recall.

### 2.6 Gene similarity ranking and query-based analysis

For individual genes of interest (e.g., ifnphi1, stat1b, usp18, rsad2), we ranked all other genes by cosine similarity in the RQ-GF embedding space. In parallel, we ranked genes by co-expression correlation (robust winsorized, Pearson, and Spearman) and merged the resulting ranked lists into a combined similarity table for comparative analysis. Thresholded gene neighborhood graphs for selected query genes were exported for network visualization in Cytoscape, retaining the top 50 neighbors above specified similarity thresholds for both embedding-based and co-expression-based rankings.

### 2.7 Sequential vector arithmetic on transcription factor embeddings

To test whether RQ-GF embeddings encode directional information about transcription factor (TF) cascades, we performed sequential vector arithmetic on L2-normalized gene embedding vectors. Starting from a composite vector representing neuromesodermal progenitors (NMPs), formed by averaging and L2-normalizing the embeddings of sox2 and tbxta, we sequentially added TFs associated with two divergent developmental fates. Neural track: sequential addition of pax6b, then pax6a. Mesodermal track: sequential addition of msgn1, then meox1, then myl1.

At each step, the new TF embedding was averaged with the existing composite vector and the result was L2-normalized. The nearest-neighbor genes of each intermediate and final composite vector were retrieved by cosine similarity, and GO enrichment analysis was performed on the top 50 neighbors to assess whether the composite vectors progressively shifted toward functional annotations consistent with the respective developmental lineages.

### 2.8 Comparison with ESM2 protein sequence embeddings

To assess whether RQ-GF captures complementary information to protein sequence-based representations, we compared nearest-neighbor gene sets in the RQ-GF embedding space with those derived from ESM2 [35], a 15-billion-parameter protein language model. For each gene, the top 50 nearest neighbors were identified in each embedding space by cosine similarity. Overlap between the two neighbor sets was quantified using the Jaccard index. This analysis was also performed between RQ-GF and the noise-trained control to establish a baseline expectation for overlap.

## 3 Characterizing zebrafish sequences found in RNAquarium

### 3.1 Identification of putative zebrafish viruses

The RNAquarium pipeline outputs both ‘flat’ files and plots summarizing the non-host transcripts found, notably heatmaps showing broad taxonomic category numbers as well as Metacoder heat trees [86] showing the taxonomic diversity of the same broad categories grouped at the family level. With our focus on viral sequences, the pipeline creates additional virus-specific outputs, notably viral transcript lists that include cluster information.

We examined the viral transcript lists to find the subset of viruses found in the RNAquarium pipeline that are associated with fish. Notably, a large fraction (*>* 60%) of the viral transcripts are classified as bacteriophage, and even the non-phage transcripts require curation. To create an initial set of possible fish-associated viruses, we manually inspected the clustered virus LCAs and pulled a subset of promising LCA viral names, resulting in a set of *≈*10,000 transcripts from 53 unique cluster LCA names (**Table 1**). We next created visualizations to examine overall nucleotide or protein similarity to known viruses (**Figure 4B**). For each of these unique clusters, we manually inspected the transcripts of each, binning the clusters into four categories based on a set of criteria. If there are no transcripts that generate open reading frames (ORFs) of at least 200 aa, we classified the cluster as “insufficient evidence”. For the remainder we initially divided the clusters into “Novel” or “Known” if the average transcript identity was greater than 90% in nucleotide space. We also examined the putative fish-associated transcripts (**Figure 4B; Supplemental Figure S7**) to rule out evidence for endogenous viruses looking specifically for any possible hits to host genomes. We determined that eight of the putative clusters belonged to a fourth category “evidence of endogenous or not fish-associated”. This category also includes clusters that we initially marked for phylogenetic analysis, but upon examination were found to not have any nearest hits to viruses known to infect fish.

### 3.2 Phylogenetic analysis of putative zebrafish viruses

For all potentially novel viruses, we selected either full length polyproteins if available, or RDRP fragments, and ran blastx searches to find the nearest known viral proteins in NCBI databases (nearest 20 hits or all hits *>* 75% aa identity). We then created protein alignments for these searches using the MAFFT-DASH algorithm [92], followed by phylogenetic analyses using raxmlng [93], carried out at the family level. Alignments and phylogenies were made using entire polyproteins (*Caliciviridae*) or portions of polyproteins that included the RDRP or polymerase (*Hantaviridae*, *Nanhypoviridae*, *Paramyxoviridae*, *Poxviridae*, Rhabdo-like; **Table 1**). For examining the novel and known recovered *Picornaviridae* transcripts in a family-level analysis, we created an alignment and phylogeny from the 3C3D portion of the polyprotein, and for examining the placement of the influenza-like viruses we created protein alignments and phylogenies for the eight influenza viral segments separately (**Figure 4C; Supplemental Figure S8**).

### 3.3 Single-cell RNA-seq analysis of Danio blood picornavirus

Runs containing Danio blood picornavirus counts were downloaded with technical reads from BioProjects PRJNA667451 (SRR12774372, SRR12774391), PRJNA943553 (SRR23824316-SRR23824323), PRJNA811542 (SRR18214218), and PRJNA1011833 (SRR25868017). Reads were mapped to the zebrafish transcriptome (GRCz12tu annotations) appended with the virus genome using cellranger-7.0.1. Data were analyzed with Monocle 3 filtering for cells with less than 40% mitochondrial read content, more than 500 UMIs and 100 genes, and fewer than 50,000 UMIs. Batch correction was applied to datasets containing more than one run. Cell type labels were annotated using marker gene sets for each cluster based on the published study or cross reference with other studies.

## 4 Machine learning to link host transcriptomes to infection biology

For each gene *g* and virus *v*, the expression shift statistic was defined as the difference in median normalized log-expression between virus-positive and virus-negative samples:

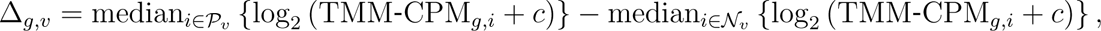

where *P_v_* and *N_v_* denote the virus-positive and virus-negative sample sets for virus *v*, respectively, and *c* denotes the prior count. Because expression values were analyzed on the log_2_ scale, Δ*_g,v_* provides a robust estimate of the log_2_-fold change in median gene expression between virus-positive and virus-negative samples. Positive values indicate higher expression in virus-positive samples, whereas negative values indicate lower expression. The median was used rather than the mean to reduce sensitivity to the heavy-tailed distributions and outliers commonly observed in RNA-seq data.

To evaluate the capacity of host transcriptomic signatures to distinguish between infection states across the 62,000 samples, we designed two distinct classification tasks based on host gene expression profiles: a binary classifier (infected versus uninfected) and a multiclass classifier to predict the specific infecting virus. Samples with zero viral counts were labeled as ‘no infection’. For the multiclass task, infected samples were assigned to specific viral categories; in instances of co-infection, the label was strictly determined by the virus with the highest relative abundance.

The dataset was partitioned into an 80% training set and a 20% hold-out test set. We implemented an SVM with a Radial Basis Function (RBF) kernel. To account for inherent data imbalance across the viral classes, we applied balanced class weights during training and incorporated L2 regularization. Model performance on the hold-out test set was evaluated using confusion matrices, Precision-Recall Curves (PRC), and the Area Under the Precision-Recall Curve (AUPRC).

While the RBF-kernel SVM provided optimal classification accuracy, we subsequently employed a linear SVM with L2 regularization to characterize the underlying gene features driving these predictions. Although linear models may not capture the complex non-linear interactions accessible to kernel-based SVMs, they offer significantly greater interpretability. Given that the classification performance of the linear SVM closely approximated that of the RBF-kernel model, we concluded that the primary biological drivers were effectively captured through linear coefficients (**Supplemental Figure S11**). Feature weights were extracted from the class-specific decision hyperplanes. Positive coefficients identify genes whose up-regulated expression serves as a predictor for a specific class, while negative coefficients denote genes whose expression levels inversely correlate with that class assignment (**Figure 5E**).

To validate the biological relevance and information content of the features selected by the linear model, we conducted a targeted feature ablation study using the RBF-kernel SVM. Specifically, the RBF model was retrained on restricted subsets containing only the top N most positively and negatively weighted genes per class, as determined by the linear SVM ranking. The resulting AUPRC scores were recorded and normalized against the AUPRC of the baseline model utilizing the entire transcriptome (**Figure 5F)**. To establish a random baseline, this process was repeated using models trained on equivalent numbers of randomly selected genes across multiple random seeds to capture performance variance (**Figure 5G**).

## 5 RNAquarium portal

The RNAquarium portal (portal.rnaquarium.org, **Figure 6**) provides a single web entry point to the reprocessed collection of Zebrafish transcriptomes and their associated microbes. The three main sections of the landing page (**Figure 6**) highlight how users can query any of the 43,091 catalogued zebrafish genes, search across the microbial and viral taxa detected in the *≈*75,000 SRA accessions reprocessed through our pipeline, or pose free-form questions to an integrated AI chatbot that is backed by MCP servers over the live metadata and visualization APIs. From the main buttons on the landing page users can click through to more detailed tools for deeper exploration. For example, the UMAP viewer (umap.rnaquarium.org) lets researchers query where in zebrafish transcriptomic space a gene is expressed, which developmental stages and tissues populate each region, and whether virus-positive samples form coherent clusters or are dispersed across the UMAP (**Figure 6, top left**). Searching for a gene or clicking through the options under Gene Analysis will give detailed information on every catalogued zebrafish gene, including (on a single page) a UMAP of all samples colored by that gene’s expression alongside the same UMAP colored by developmental stage and by tissue type, violin plots of expression by developmental stage and by tissue, and two gene-embedding summaries: a cross-model similarity chart and an interactive nearest-neighbor graph (**Figure 6, top right**). Other portal tools include the Gene Expression viewer (gene-xp.rnaquarium.org) and the Gene Embeddings viewer (embeddings.rnaquarium.org). The Gene Expression viewer is for multi-gene comparisons, including tissue- and stage-resolved violin plots, clustered heatmaps, pairwise gene scatter plots, and virus-conditioned robust-shift plots contrasting virus-positive with virus-negative samples, while the Gene Embeddings viewer is for nearest-neighbor search across three embedding spaces (a Geneformer model trained on RNAquarium, a Zebrahub-fine-tuned Geneformer, and the ESM2 protein language model), with results rendered as ranked similarity tables and interactive network graphs. Microbial and viral diversity is explored through the taxon viewer (**Figure 6, bottom left**), which couples an NCBI taxonomy tree pruned to organisms with detected transcripts to per-node transcript-count bar charts, and can be restricted to the fish-associated viral transcripts recovered from the corpus. A dedicated Data page provides access to all datasets generated by the project, and is accompanied by a Metadata portal section (metadata.rnaquarium.org/) that lets users query sample- and contig-level metadata interactively in the browser, without needing to download the full archive.

The integrated AI chatbot is designed to lower the barrier to this resource further by making both the data and the analyses built on top of it accessible through natural language. Researchers can ask questions such as “how does tp53 expression change across zebrafish developmental stages?”, “show me a scatter plot comparing tp53 and isg15 expression”, or “which zebrafish genes are most impacted by viral infections?”, and the agent answers by routing the request through the same metadata and visualization APIs that back the portal, running code in a sandbox environment, and returning figures, tables, and data-grounded explanations on the fly, so that quick plots, exploratory analyses, and data-driven insights become accessible to researchers who lack the programming expertise or compute infrastructure to run them themselves. Together, these entry points let users move seamlessly between gene, sample, and taxon views of the same underlying corpus, supporting hypothesis generation, candidate-gene prioritization, and cross-study comparison without requiring local compute or command-line expertise.

Details of the portal architecture and backend components are listed in **Supplemental Table S4**.

