## Supplemental Figures S1 to S16 Tables S1 to S4 for "RNAquarium: an archive-scale atlas of zebrafish gene expression coupled with pan-taxonomic profiling reveals diverse viral drivers of transcriptomic states"

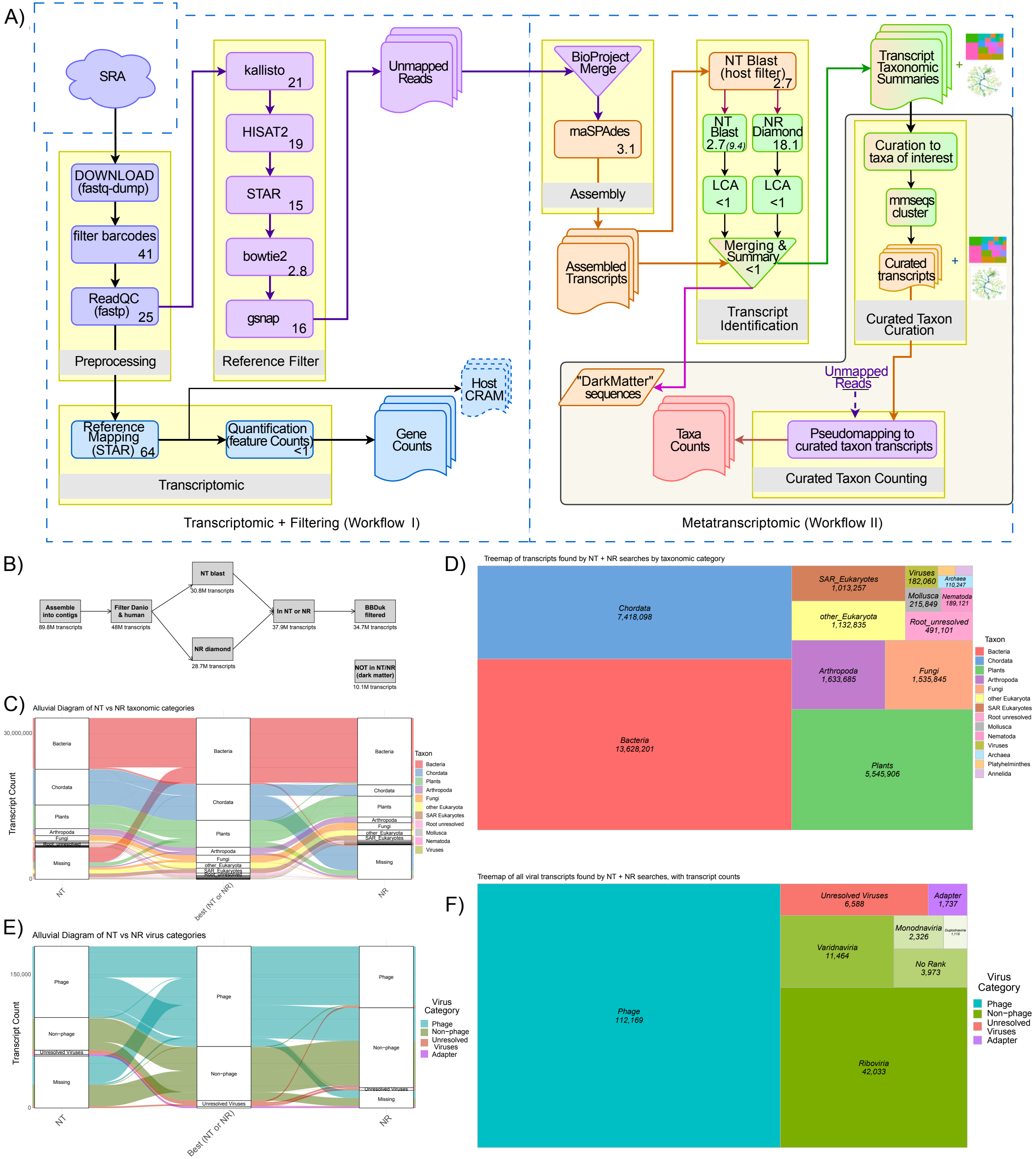


**Supplemental Figure S1. Details of RNAquarium pipeline, related to Figure 1.** (A) Detailed schematic of RNAquarium pipeline. The boxed region in the Metatranscriptomic portion of the workflow are steps run outside of Nextflow using SLURM. Values in boxes indicate the processing time (in thousands of CPU-hours) of each step in the Nextflow pipelines for our analysis of ≈75,000 zebrafish RNA-seq SRA runs, with the value in italics indicating the initial time of the full NT Blast processing before optimization. (B) Non-host transcript numbers found using the RNAquarium pipeline on ≈75,000 zebrafish RNA-seq datasets. (C) Alluvial plot showing numbers of non-host transcripts found via NT or NR searches. (D) Treemap of non-host transcripts. (E) Alluvial plot showing numbers of viral transcripts found via NT or NR searches. (F) Treemap of viral transcripts.

**Supplemental Figure S2. Seq-Detective filtering decisions by submitted format, related to STAR Methods.** SRA B/BB decision inferred from fastq-dump outputs without --include-technical. B→T and BB→TT: all mates failed basic mapping thresholds (e.g. blanks/quality issues/orphaned technical mate/too long for short read processing).

| all (n = 77,188) | | | | | | | |
|---|---|---|---|---|---|---|---|
| | SRA = B (single-end) | | SRA = BB (paired-end) | | | | |
| Submitter format | B → B | B → T | BB → BB | BB → TB | BB → BT | BB → TT | n |
| fastq ×2 | 4% | 0% | 81% | 8% | 1% | 4% | 33309 |
| fastq | 95% | 4% | 0% | | | 0% | 18722 |
| cram | 0% | | 32% | 63% | 2% | 1% | 16892 |
| bam | 61% | 0% | 29% | 8% | 0% | 0% | 2695 |
| fastq ×3 | 92% | 0% | 2% | 2% | 2% | | 2439 |
| unknown/NA | 54% | 3% | 39% | 0% | 0% | 1% | 1329 |
| fastq ×4 | 54% | 0% | 28% | 16% | | | 696 |
| bam+cram | | | 89% | | | 10% | 568 |
| 10X bam | 97% | | | 2% | | | 179 |
| fastq ×8 | 13% | | 29% | 56% | | | 65 |
| SOLiD | 100% | | | | | | 65 |
| fastq ×6 | 8% | | 91% | | | | 60 |
| fastq ×16 | 51% | | | 9% | 25% | 13% | 52 |
| bam ×2 | 88% | | 11% | | | | 34 |
| srf | | | 100% | | | | 28 |
| Illumina native | 58% | 33% | 8% | | | | 12 |
| nanopore | 18% | 81% | | | | | 11 |
| fastq ×12 | 14% | | 28% | | 57% | | 7 |
| fastq ×24 | | | | 33% | 66% | | 3 |
| fastq ×48 | | | | | 100% | | 3 |
| fastq ×14 | 100% | | | | | | 2 |
| fastq ×9 | 50% | | | 50% | | | 2 |
| pacbio | | | 100% | | | | 2 |
| bam ×3 | | | 100% | | | | 2 |
| fastq ×32 | | | | | 100% | | 2 |
| fastq ×10 | 100% | | | | | | 2 |
| fastq ×15 | 100% | | | | | | 1 |
| fastq ×7 | 100% | | | | | | 1 |
| fastq ×42 | | | 100% | | | | 1 |
| fastq ×13 | 100% | | | | | | 1 |
| bam+fastq | | | 100% | | | | 1 |
| fastq ×18 | | | 100% | | | | 1 |
| fastq ×38 | | | 100% | | | | 1 |

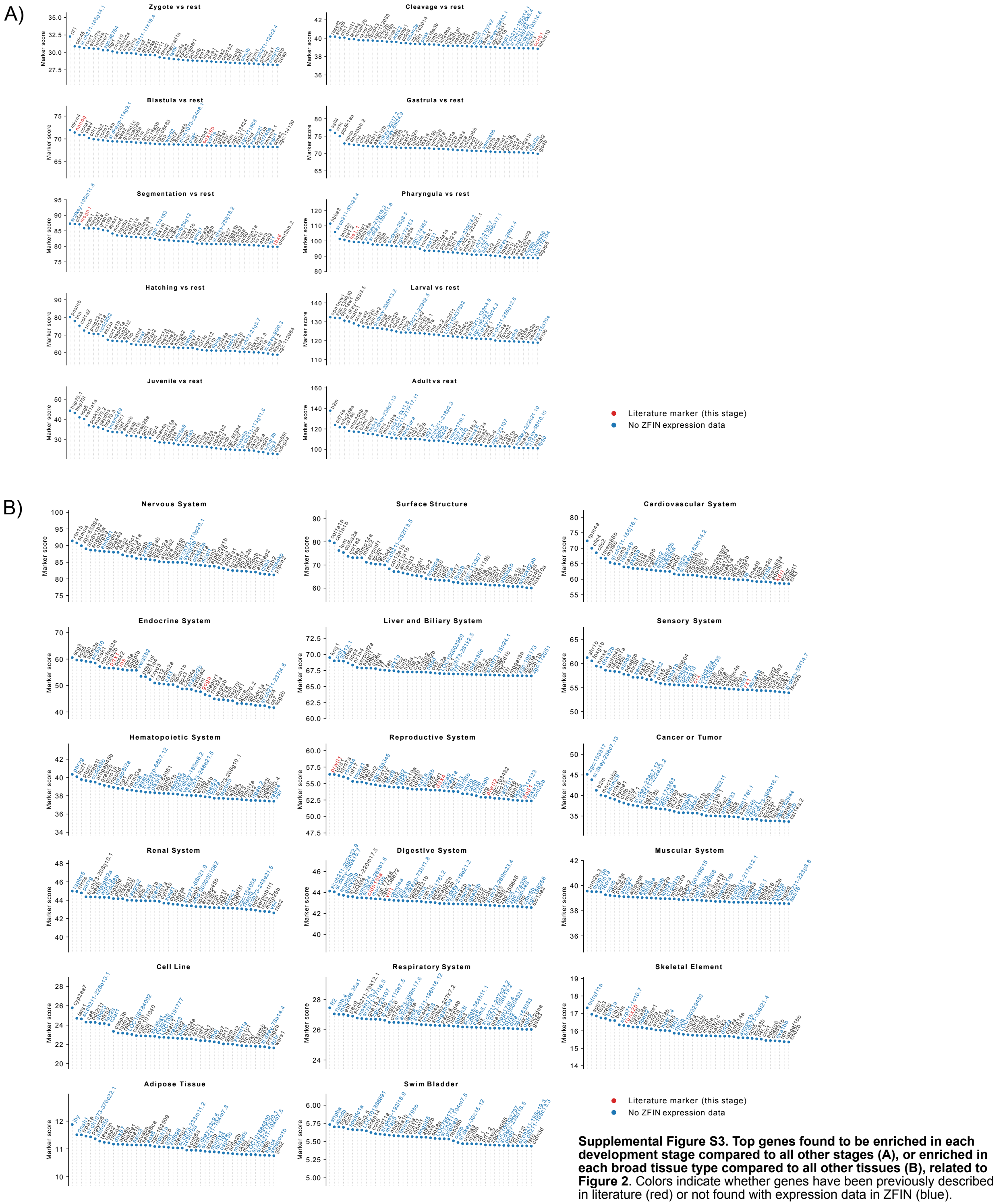


**Supplemental Figure S3. Top genes found to be enriched in each development stage compared to all other stages (A), or enriched in each broad tissue type compared to all other tissues (B), related to Figure 2**. Colors indicate whether genes have been previously described in literature (red) or not found with expression data in ZFIN (blue).

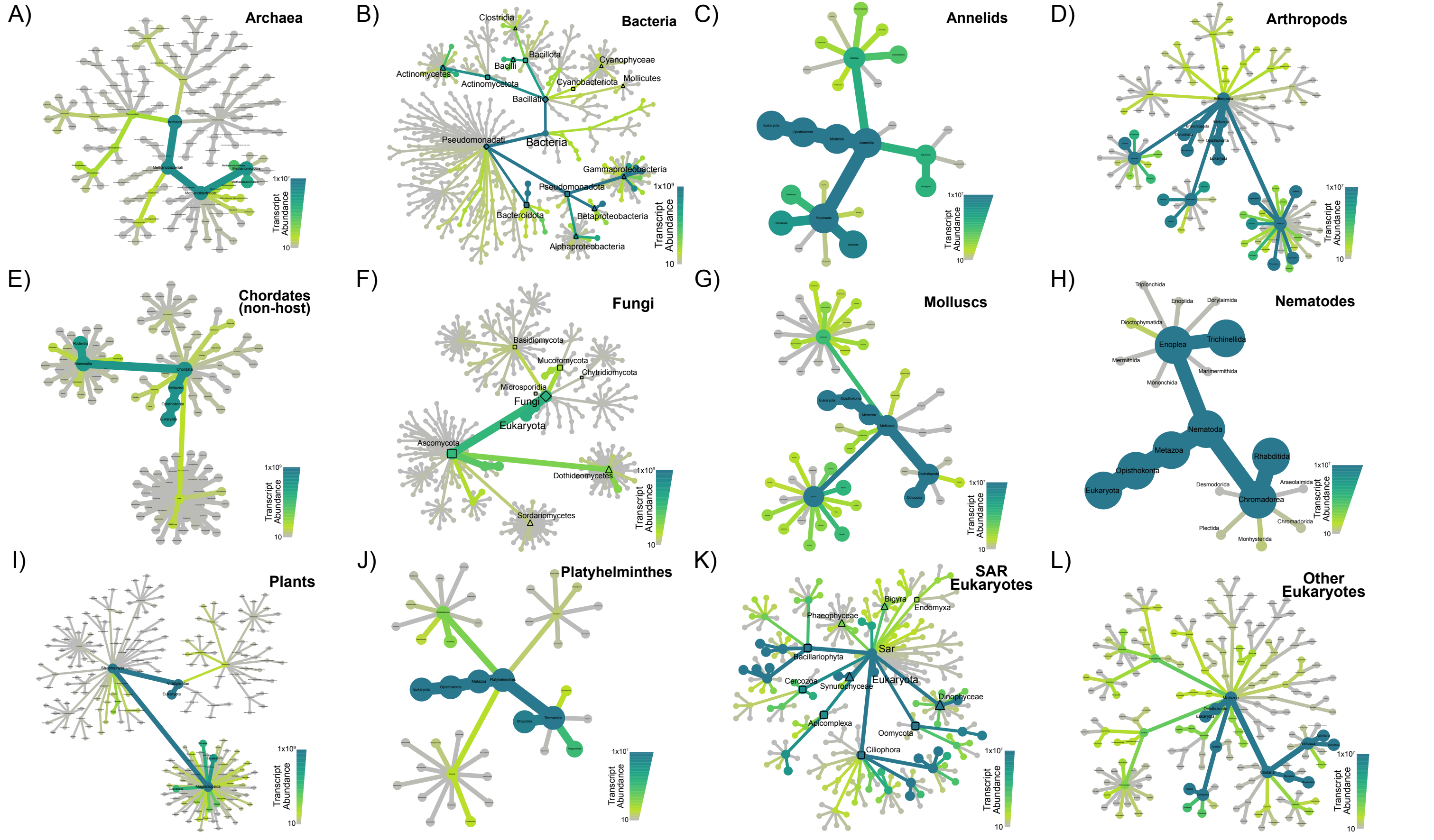


**Supplemental Figure S4. Metacoder heat trees of non-host transcripts showing phylogenetic diversity (to orders) of RNAquarium results, related to Figure 2**. Heat trees of (A) Archaea, (B) Bacteria, (C) Annelids, (D) Arthropods, (E) Chordates (non-host), (F) Fungi, (G) Molluscs, (H) Nematodes, (I) Plants, (J) Platyhelminthes, (K) SAR Eukaryotes, and (L) other Eukaryotes. Shapes in heat trees indicate taxonomic level: diamonds = kingdom; squares = phylum; open circles = orders, black circles = families. Both width of tree branches and shade are scaled by cumulative search result transcript length x coverage 'abundance' values in each clade.

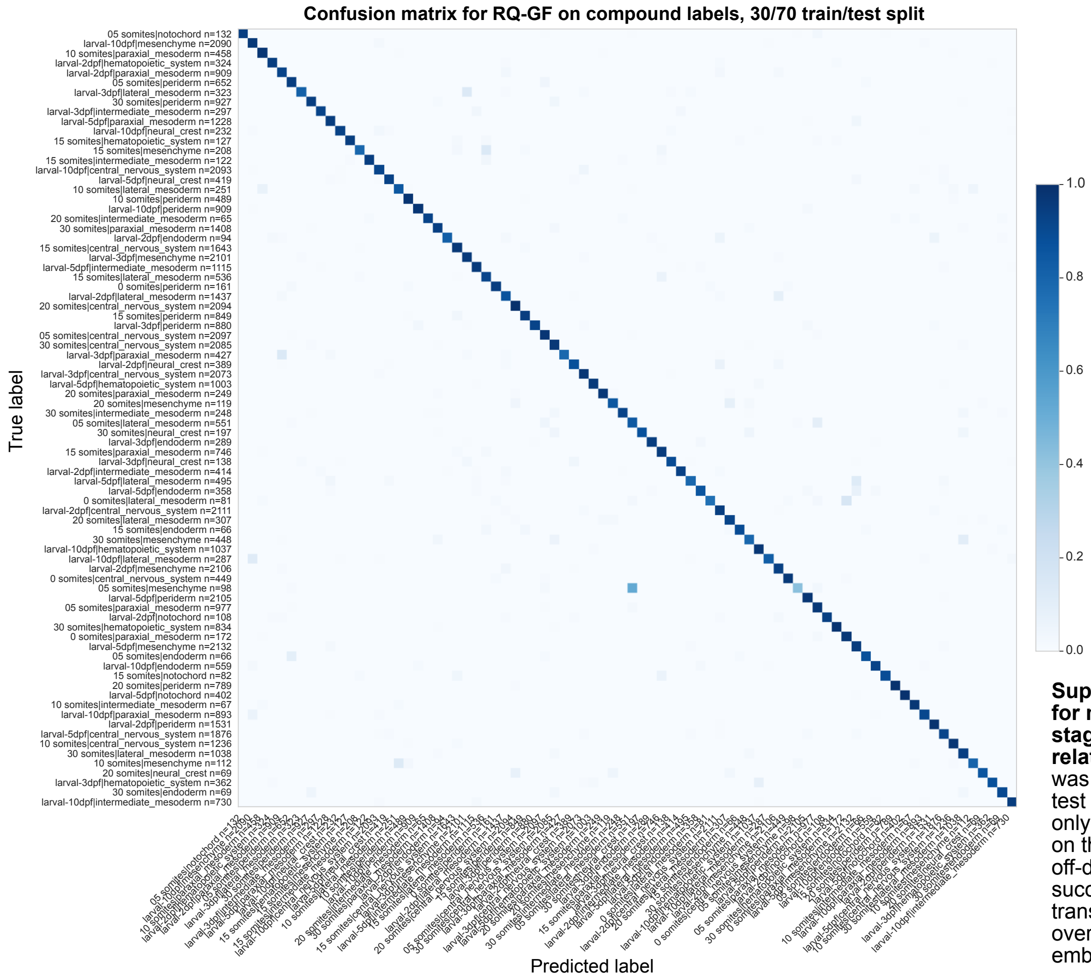


**Supplemental Figure S5. Confusion matrix for multiclass compound developmental stage and tissue label prediction by RQ-GF, related to Figure 3**. Classification performance was evaluated using a challenging 30/70 train/test split, where the model was fine-tuned on only 30% of the Zebrahub dataset and tested on the remaining 70% holdout set. The minimal off-diagonal density confirms that RQ-GF successfully distinguishes subtle, transcriptomically similar cell states across overlapping developmental windows and embryonic lineages.

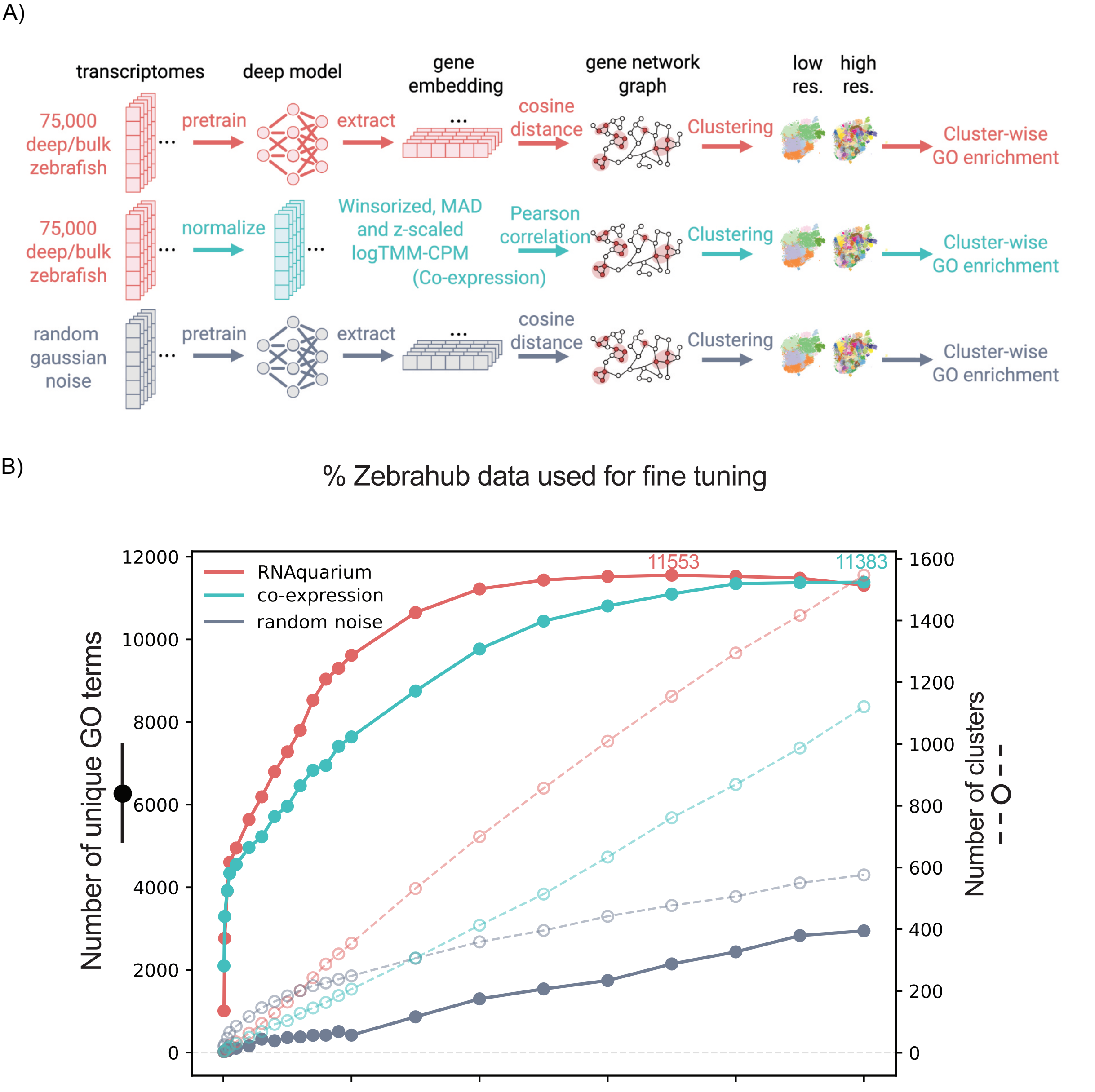


**Supplemental Figure S6. AI modeling of archive-wide transcriptomes and analysis of gene representations, related to Figure 3**. (A) Workflow for gene network construction and biological pathway recovery. Gene networks were generated via pairwise cosine distances of extracted RQ-GF gene embeddings (top), standard log-transformed Pearson co-expression profiling from identical input transcriptomes (middle), or random Gaussian noise embeddings (bottom), followed by multi-resolution Leiden clustering and cluster-wise Gene Ontology (GO) enrichment. (B) Functional recall of unique biological terms across clustering resolutions. The number of unique GO terms successfully recovered (solid lines, left y-axis) and the total number of clusters generated (dashed lines, right y-axis) plotted against Leiden clustering resolutions. Peak unique GO terms recovered for RQ-GF and co-expression are annotated at the top right.

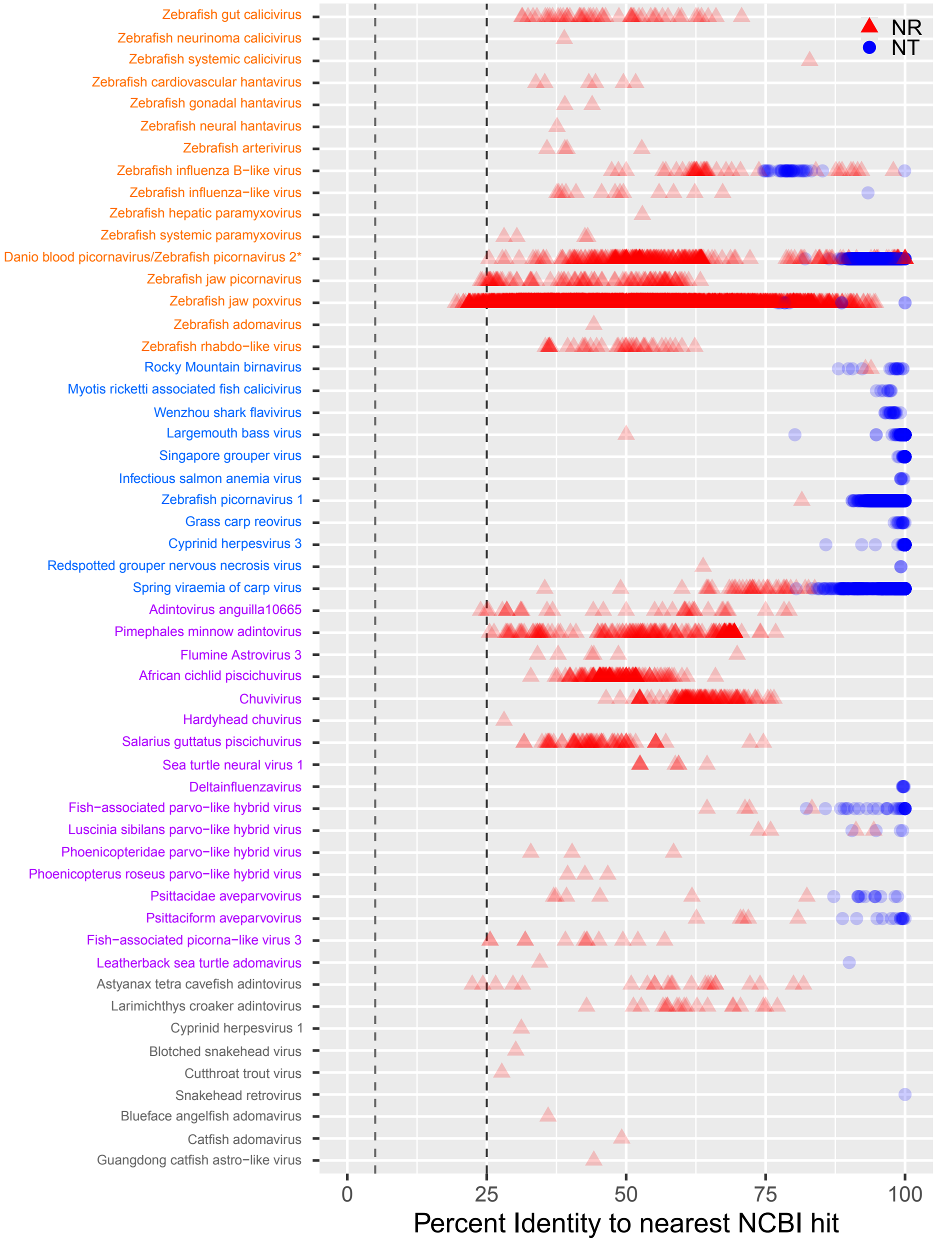


**Supplemental Figure S7. Nucleotide (NT; circles) and protein (NR; triangles) identity scores for 53 putative fish-associated viruses, related to Figure 4 and Table 1**. Colors of virus names correspond to curation category: orange - novel; blue - known; purple - endogenous or not fish-associated; gray - insufficient evidence. The asterisk refers to the Danio blood picornavirus, which has previously been included in the NCBI database but poorly characterized. The final three viruses in the known category were intentionally administered. Dashed lines indicate null expectations of NT identity (25%) and NR identity (5%).

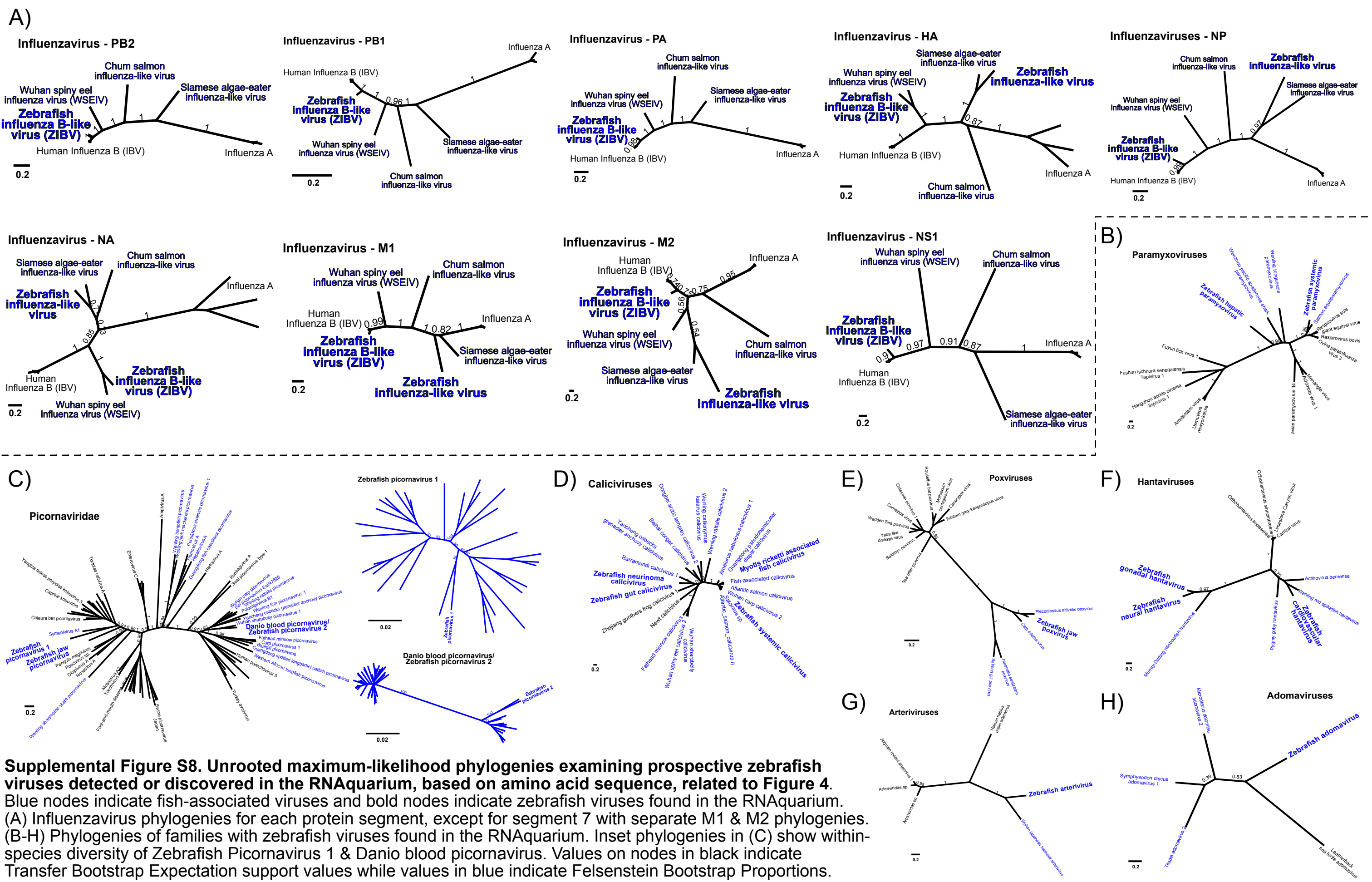


**Supplemental Figure S8. Unrooted maximum-likelihood phylogenies examining prospective zebrafish viruses detected or discovered in the RNAquarium, based on amino acid sequence, related to Figure 4.** Blue nodes indicate fish-associated viruses and bold nodes indicate zebrafish viruses found in the RNAquarium. (A) Influenzavirus phylogenies for each protein segment, except for segment 7 with separate M1 & M2 phylogenies. (B-H) Phylogenies of families with zebrafish viruses found in the RNAquarium. Inset phylogenies in (C) show within-species diversity of Zebrafish Picornavirus 1 & Danio blood picornavirus. Values on nodes in black indicate Transfer Bootstrap Expectation support values while values in blue indicate Felsenstein Bootstrap Proportions.

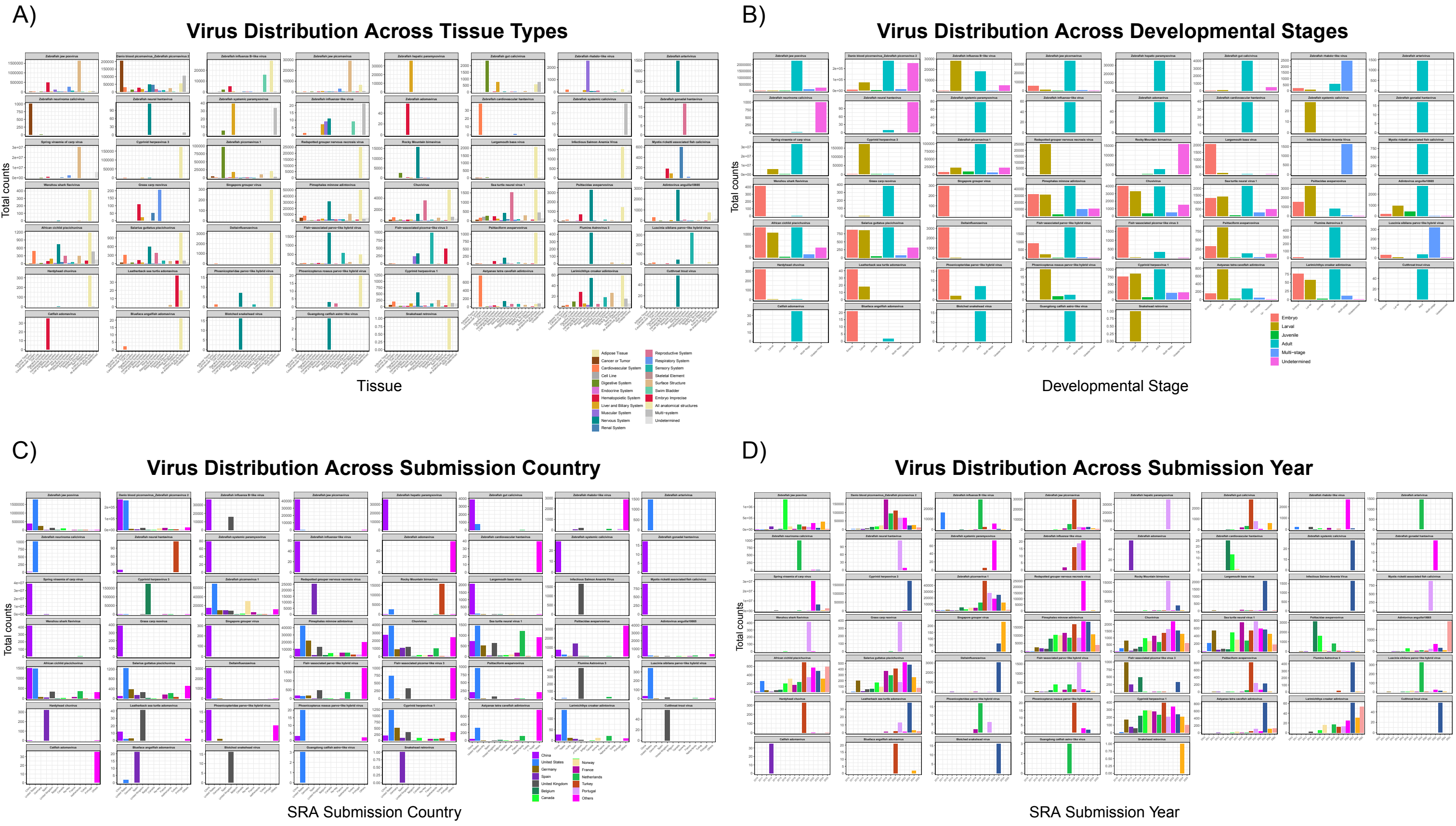


**Supplemental Figure S9. Counts of 53 putative fish-associated viruses across RNA-seq runs by attribute, related to Figure 4.** (A) Counts by curated tissue category. (B) Counts by developmental stage. (C) Counts by country of submission (top 12 + all others). (D) Counts by year of submission.

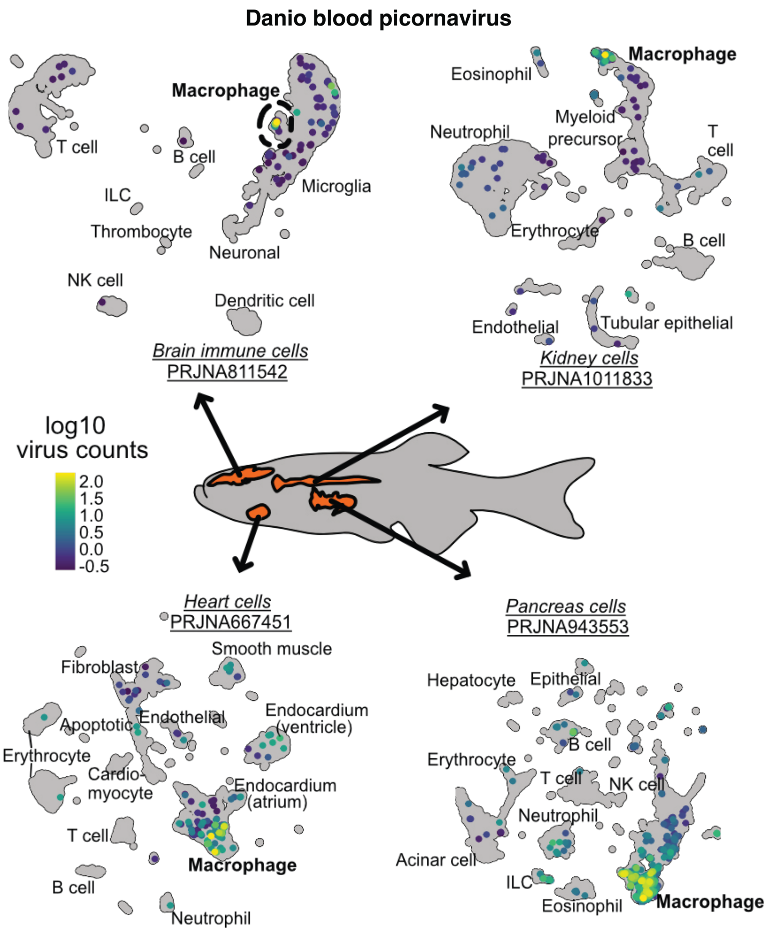


**Supplemental Figure S10. Danio blood picornavirus in single-cell RNA-seq datasets, related to Figure 4.**

Detection of Danio blood picornavirus (DBPV) in four single-cell RNA-seq datasets, each from a different tissue. Virus counts are plotted on UMAPs annotated with cell type labels. In all four studies, macrophages were the predominant cell type containing DBPV transcripts.

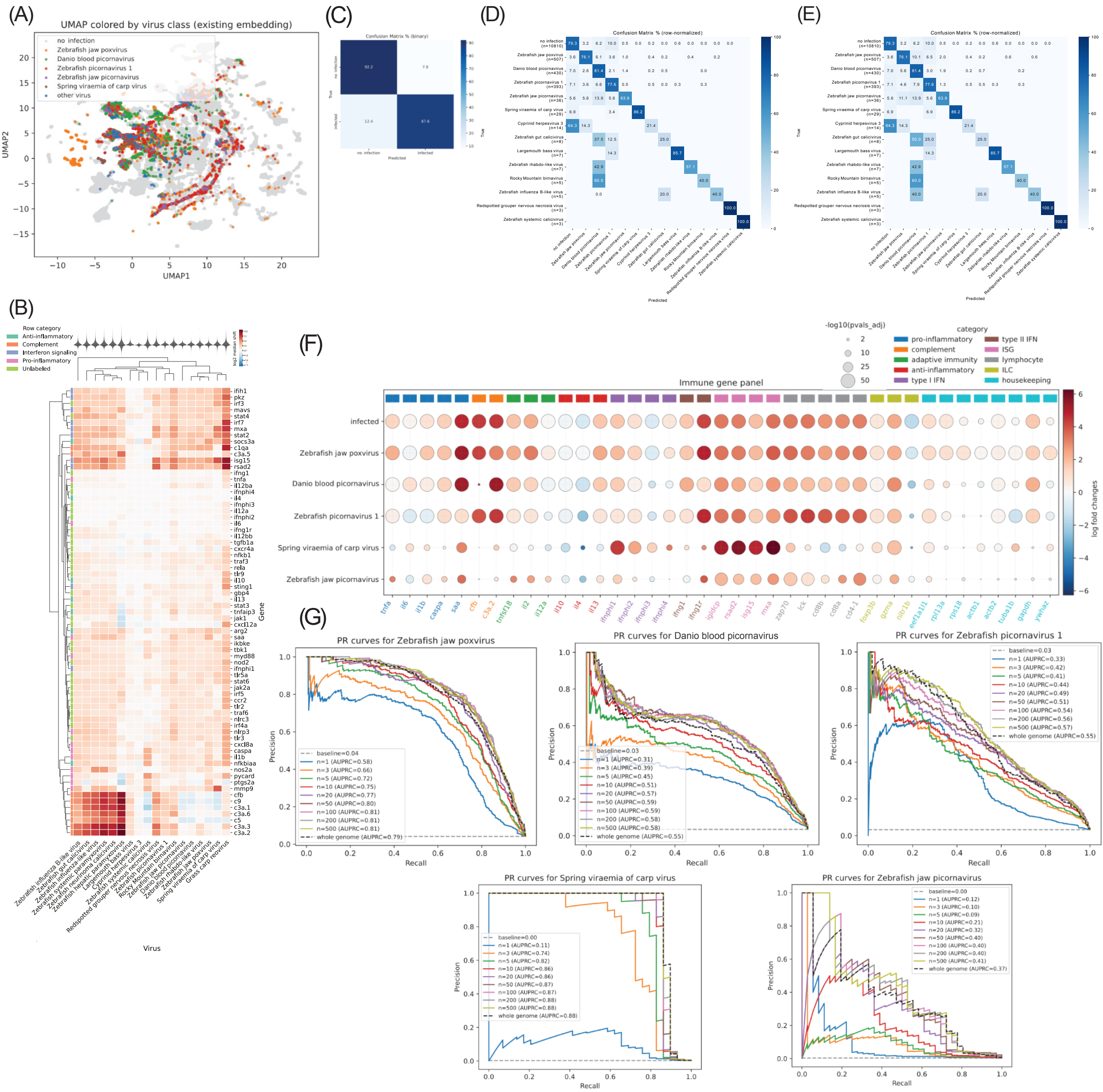


**Supplemental Figure S11**. **Linking host transcriptomes to infection biology, related to Figure 5.** (A) UMAP visualization of viral association states. Zebrafish transcriptomes are projected onto a pre-existing UMAP embedding, with individual samples colored by the dominant viral transcript identified. (B) Heatmap of the log2 median shift for a curated panel of immune marker genes (rows), grouped by functional category, across 18 fish-associated viruses (columns), each detected in ≥10 transcriptomes. Rows and columns are hierarchically clustered. The mini-violin above each column shows that virus's genome-wide shift distribution, providing the transcriptome-wide baseline against which the immune-gene shifts can be read. Clustering reveals both pan-viral host responses, such as interferon-stimulated genes like *isg15*, and virus-specific transcriptional signatures. (C) Binary classification confusion matrix showing performance of the SVM model in distinguishing between "No infection" and "Infected" states. (D) Confusion matrix showing the row-normalized classification performance for predicting the dominant virus transcript. High diagonal values indicate strong predictive accuracy across most viral classes. To focus on the host response specific to the dominant infection, off-diagonal instances where the predicted virus was present as a non-dominant transcript were excluded from the visualization. Consequently, row totals in the matrix may not sum to 100%, as these ambiguous co-infection cases were removed from the off-diagonal without being reclassified as primary hits. (E) High-confidence multiclass "strict" confusion matrix, showing a refined version of the multiclass prediction matrix in (D) but using more restrictive criteria for classification. (F) Differential expression of host markers in true-positive samples. Dot plot showing the expression of key immune and inflammatory genes across viral classes. To ensure transcriptomic signals reflect active infection, this analysis is restricted to "true positive" samples (those with detectable viral transcripts). The color intensity represents the magnitude of the log2fold change compared to non-infected controls, while dot size indicates the percentage of samples within that class expressing the gene. (G) Class-specific Precision-Recall (PR) curves. Detailed performance metrics for the top five viral classes (Zebrafish jaw poxvirus, Danio blood picornavirus, Zebrafish picornavirus 1, Spring viraemia of carp virus, and Zebrafish jaw picornavirus). Curves illustrate the model's performance as the selection of the number of top positive/negative genes N increases from 1 to 500 from each class, with the dashed black line representing the whole-genome baseline.

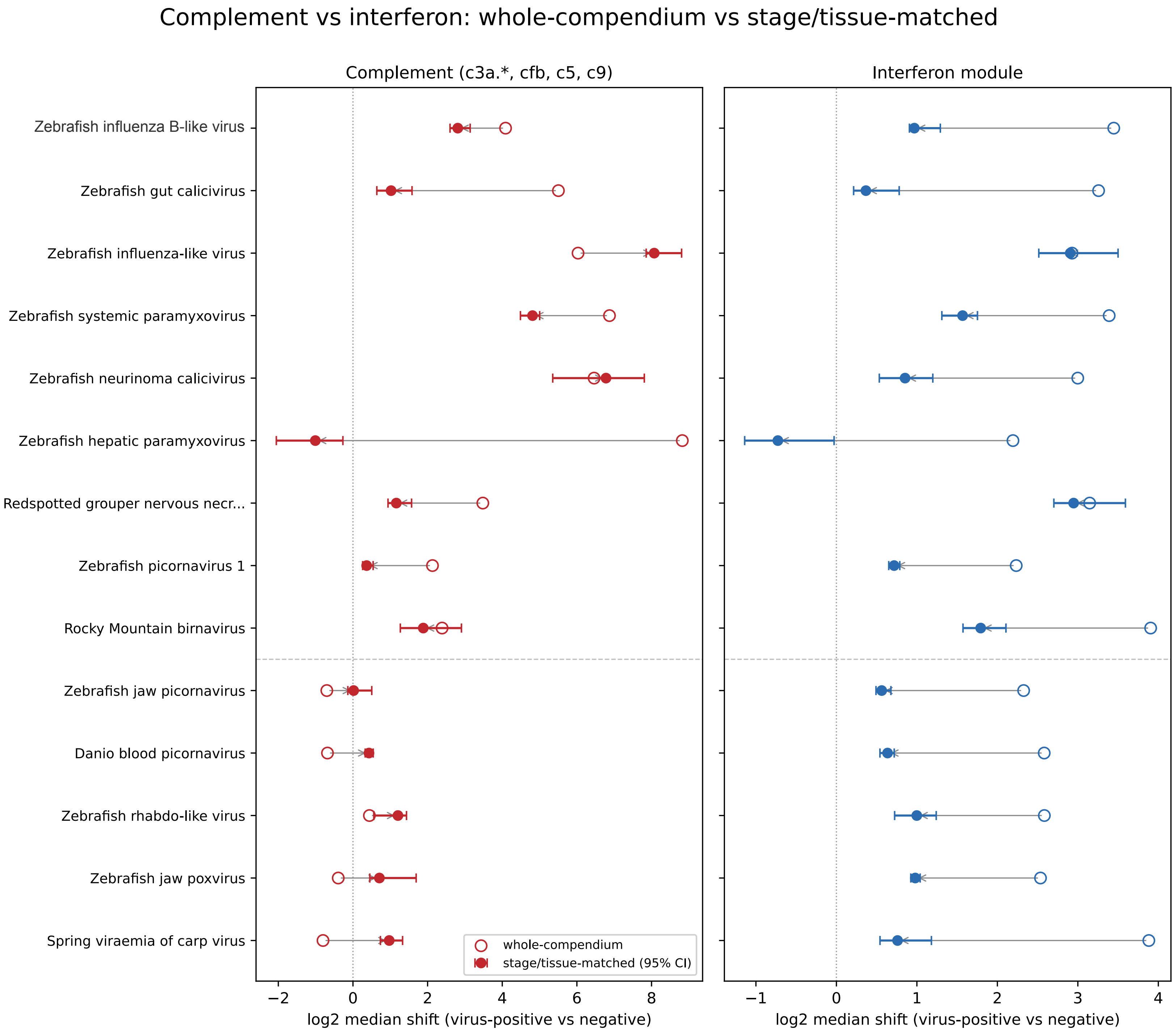


**Supplemental Figure S12. Viral expression shift validation summary, related to Figure 5.**
To control for atlas composition, we recomputed viral expression shifts (log2 median expression shift of virus-positive vs. negative runs) within matched developmental-stage and tissue strata (closed circles) relative to shifts across the whole compendium (open circles), for complement genes (left panel) and for interferon module genes (right panel). Shifts were calculated for 14 fish-associated viruses described in the results and **Figure 5B and Supplemental Figure S11** — the first nine are viruses associated with a broad innate immune program (above dashed line), and the last five are associated only with the interferon module (below dashed line).

**Supplemental Figure S13. Seq-Detective supplemental plots, related to STAR Methods.** (A) Seq-Detective per-mate judgment (biological "B" vs. technical "T") vs. mate1/mate2 mapping rate, for SRA runs with two read files, both already tagged READ_TYPE=B by SRA (the population this pipeline downloads) as of 2025-04. Unsurprisingly, Seq-Detective's single-technical-mate calls "T B" and "B T" are correlated with cases where one mate file exhibits low mappability relative to the other, i.e., cases where SRA's own biological/technical tagging failed to catch a technical mate. Cases where neither file can be reliably mapped after fastp trimming ("T T") are also rejected by Seq-Detective. (B) Seq-Detective per-mate filtering decisions by gene sparsity. Left: single-end runs, mapping rate (log scale) vs gene sparsity. Right: paired-end runs (runs with two read files designated READ_TYPE=B by SRA), mate1/mate2 mapping ratio (log scale) vs minimum gene sparsity. In the paired-end panel, "T T" shows extreme sparsity (a pattern consistent with some alignments to gene features by chance or debris in an "empty" well); these mates have been rejected by Seq-Detective. Paired-end runs with mate mapping ratio near 1 exhibit gene sparsity anywhere from almost 0.6 (40% of all annotated features have at least 1 count) to 1.0 (no annotated genes detected), consistent with large scale paired-end bulk assays and highly-specific single-cell Smart-Seq2 protocols being represented in this category. In the single-end panel, "T" filtering outcomes runs also cluster at low mapping rate and high sparsity. (C) Gene sparsity vs mapping rate/ratio, annotated by metadata-derived technology labels. In the paired-end panel, two separate clusters of DETCT runs are visible, one with a partially technical mate 1 and higher minimum gene detection sparsity, and another cluster with a partially technical mate 2 and lower minimum gene detection sparsity. A loose clustering of 10x-labeled runs is seen around 0.8 sparsity in the paired-end panel, while a tight cluster of 10x-labeled runs at 0.98-0.99 sparsity appears only in the single-end panel, at mapping rates near 0 and rejected as T by Seq-Detective, an outcome consistent with submissions where the cell barcode or index read was mistakenly uploaded as a separate run accession. While for the most part, paired-end Smart-Seq runs correspond to a mate ratio near 1, we also encountered modified Smart-Seq protocols with partially barcoded reads in the dataset, which can be seen here in the lower-sparsity clusters off the midline according to the read modified.

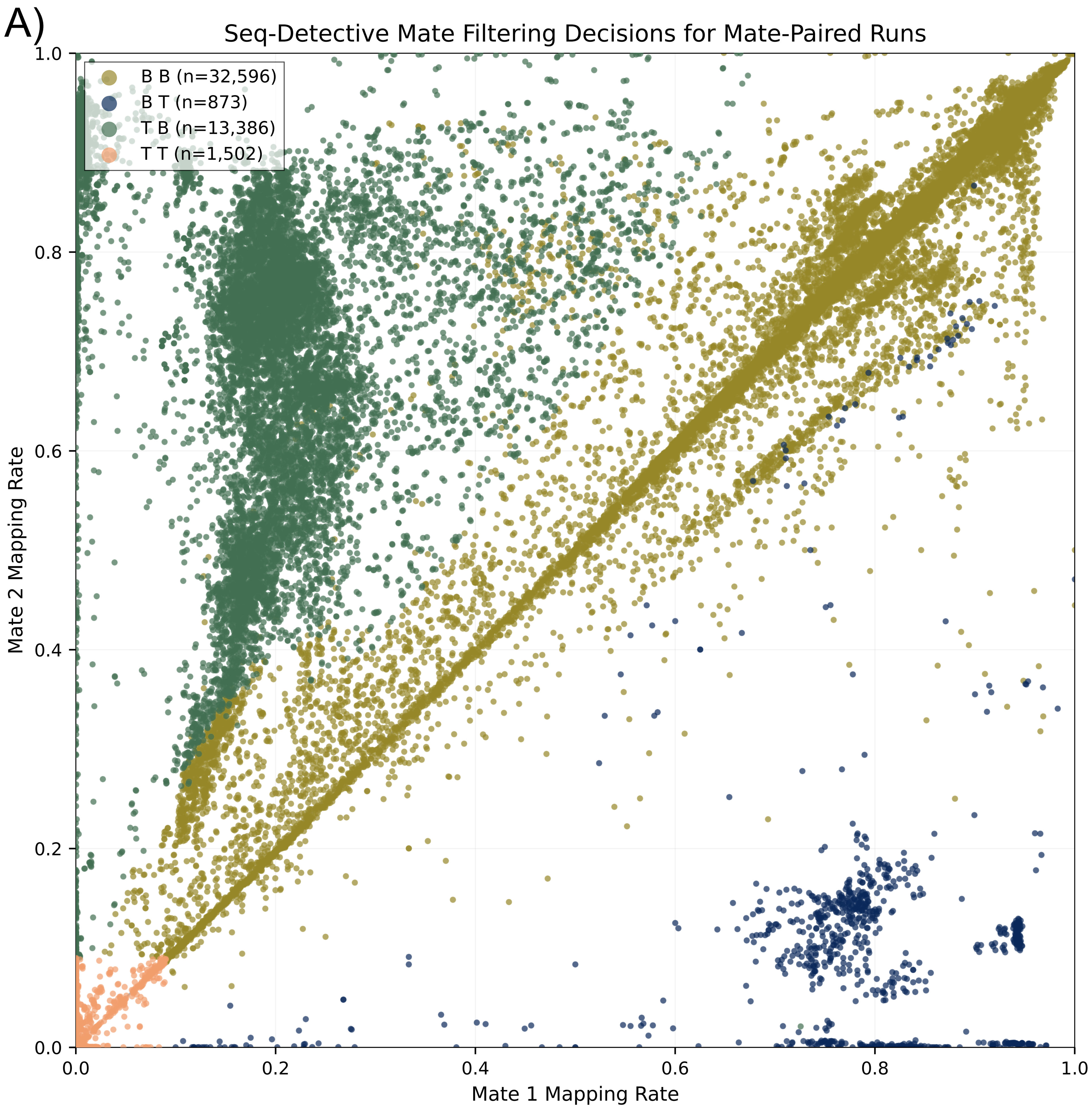


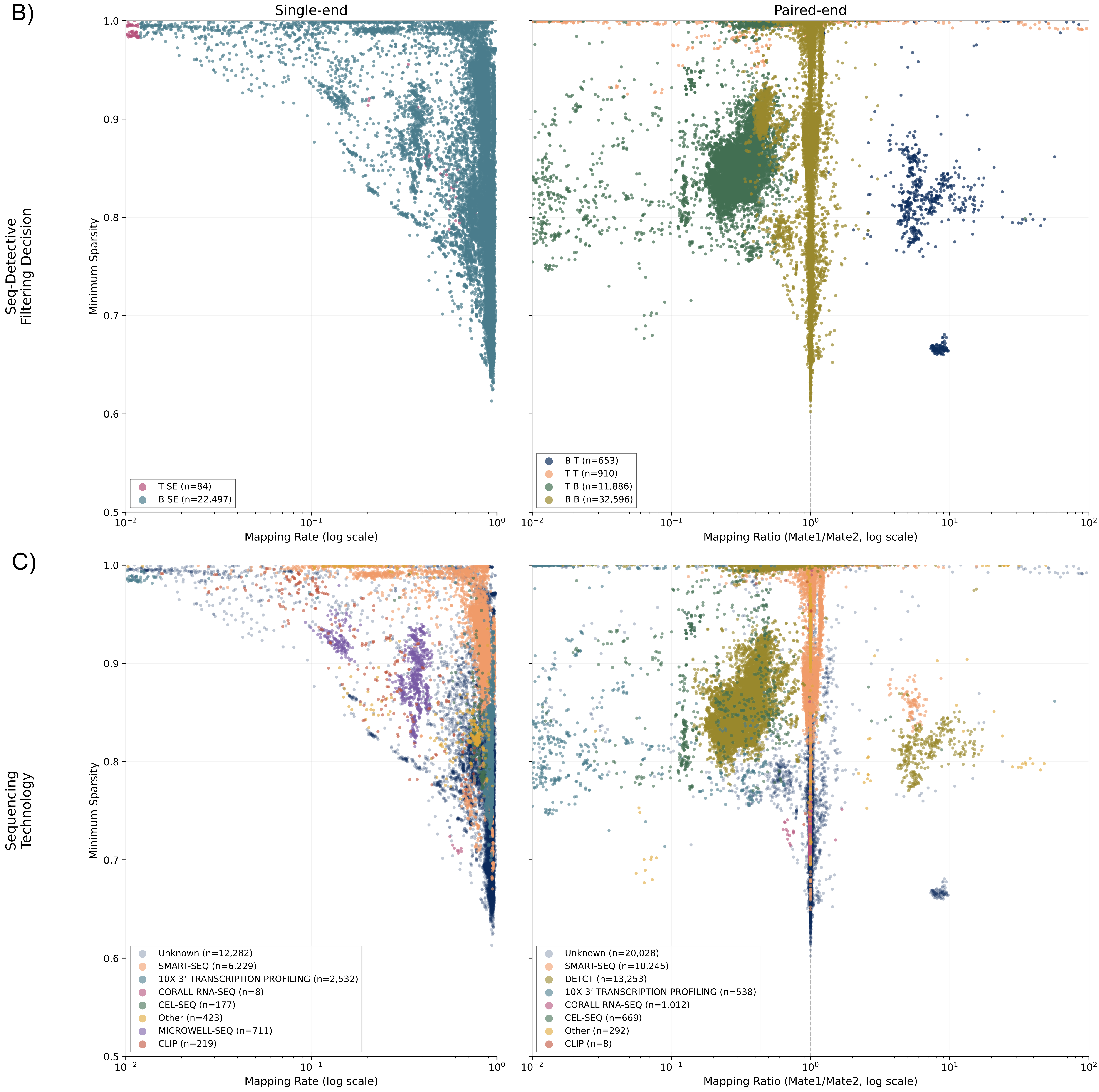

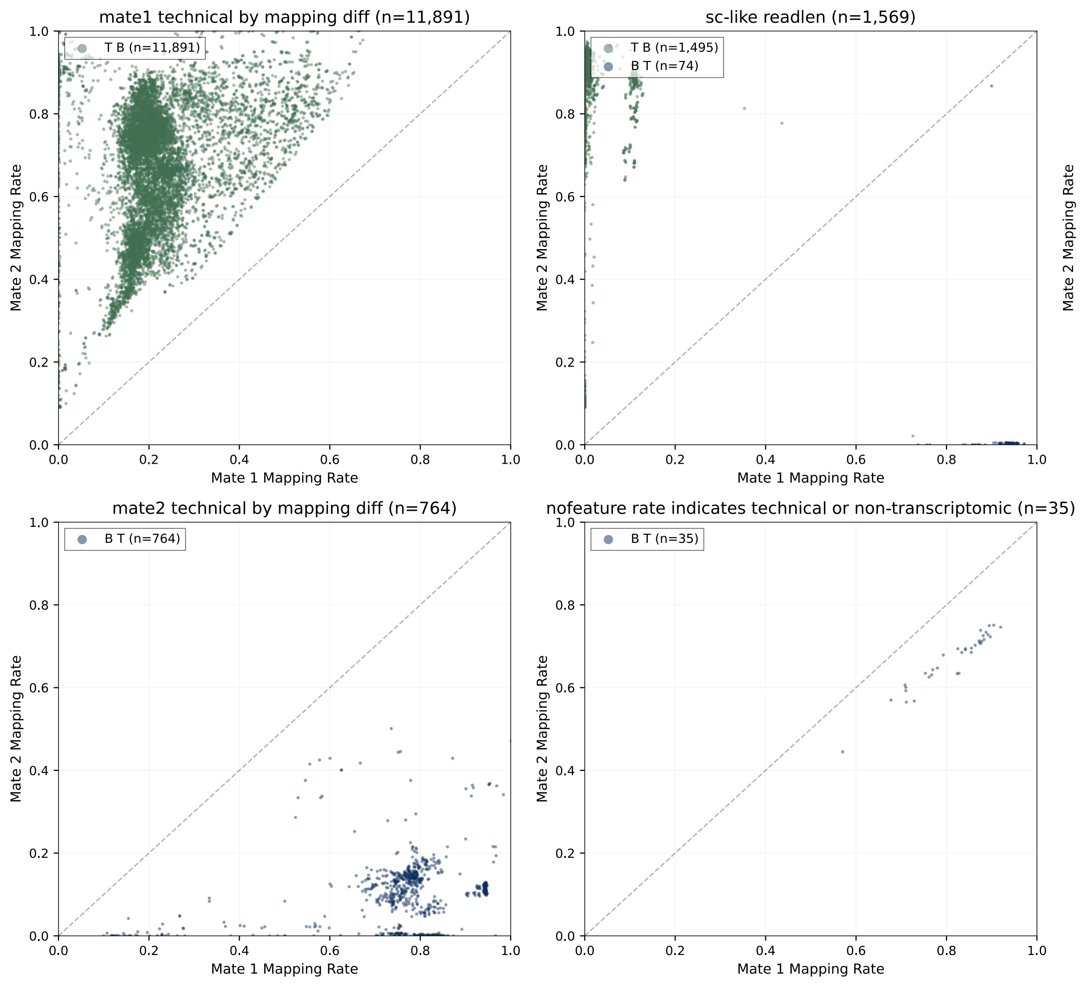


**Supplemental Figure S14. Seq-Detective paired decision "reason" groups, related to STAR Methods.**
Mate 1 vs. Mate 2 mapping rate for runs where SRA's metadata designates both mate files "biological" (B) (as of the 2025-04 processing date in this analysis), but Seq-Detective's re-derived classification assigns "technical" (T) to at least one mate. Runs where Seq-Detective agrees with SRA's B/B labeling are excluded. Panels separate these disagreements by the "reason" field Seq-Detective reports, the branch of its decision tree that drove the T call for each mate (see **Supplemental Figure S13** legend T/B classification scheme).

**Supplemental Figure S15. Seq-Detective validation summary, related to STAR Methods.**
Accessions were selected for unique combinations of study + submission format + metadata-inferred technology (see **Supplemental Data S2** describing the initial 156 accession set). Accessions were further annotated based on available metadata, sequencing read layout, and available study information to determine actual sequencing technology and possible issues. The final validation set of 150 accessions were curated by taking plausible, available, RNA-seq datasets without known blanks and with a human-inferred sequencing method, then adding curator expected filtering decision.

| Technology | n | Agreement % | Remark |
|---|---|---|---|
| bulk | 47 | 95.7 | 2x 36bp piRNA on Illumina Genome Analyzer II judged low mapping |
| 10x | 40 | 100.0 | |
| smartseq | 13 | 92.3 | |
| other | 50 | 92.0 | 1x 454 and 1x microwell-seq show quality issues on inspection<br>1x DeTCT "T B" judged "B B" from partial-T high mapping rate |
| **Total** | **150** | **95.3** | |

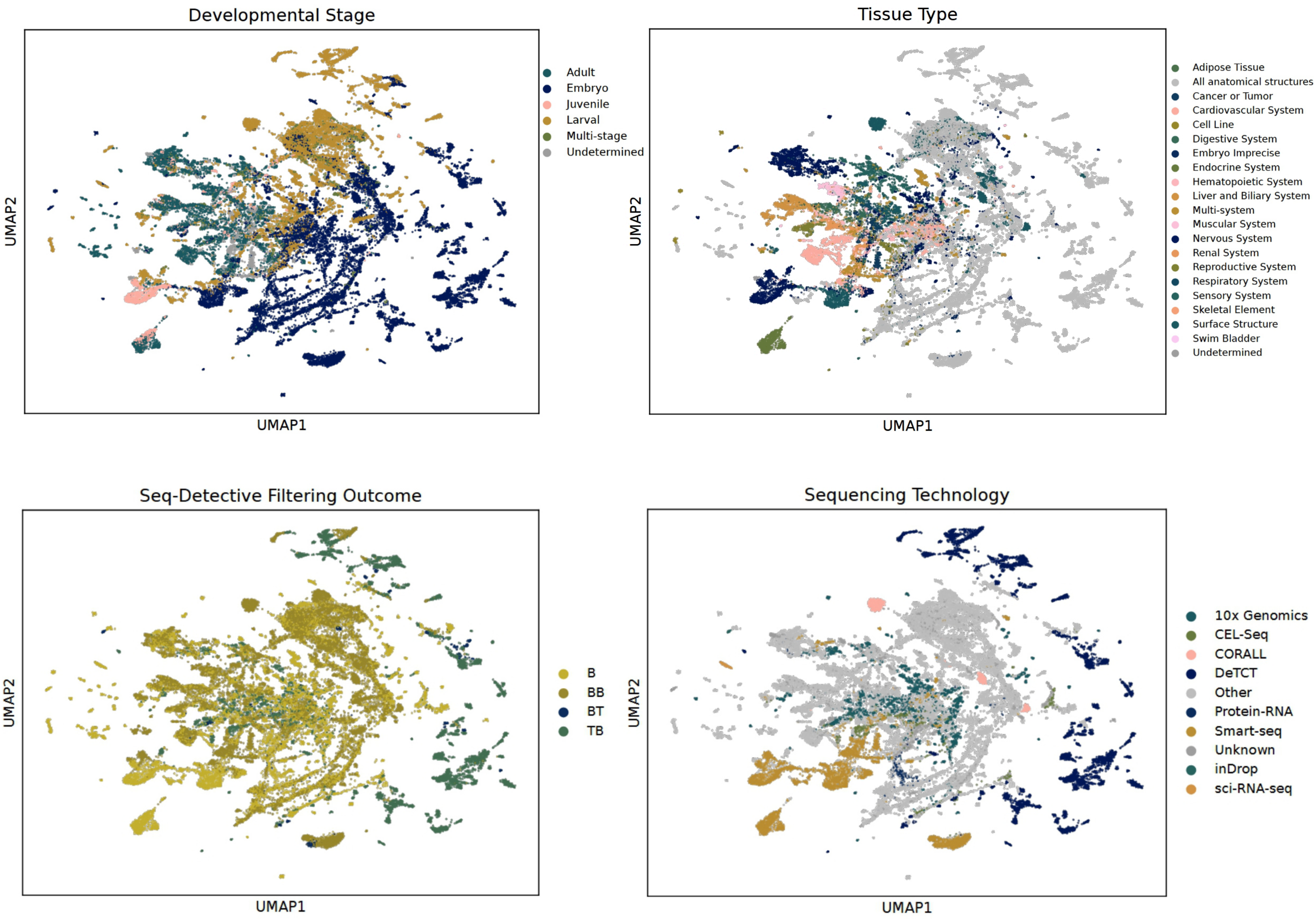


**Supplemental Figure S16. UMAP colored by Seq-Detective filtering outcome and metadata-derived technology (same embedding as Figure 2), related to Figure 2 and STAR Methods.** (Above left, right) developmental stage and tissue-type UMAPs from **Figure 2A and 2C**, shown for reference. (Below left) the same UMAP colored by Seq-Detective filtering outcome. (Below right) the same UMAP colored by metadata-derived sequencing technology group. DeTCT forms a distinct, separated cluster, consistent with its reads retaining partial technical sequence even after Seq-Detective's whole-mate classification resolution.

**Supplemental Table S1. Comparison of software tools and portals for metagenomic analyses of RNA-seq data, related to Introduction.**

| | Citation | Scale | Scope | Approach | Key output | Interactive data portal | Methods | | | | | |
|---|---|---|---|---|---|---|---|---|---|---|---|---|
| | | | | | | | Host transcriptomics | Taxonomy method | Non-host target | Assembly | Quantification | Compute |
| **CZ ID** | Simmonds et al. 2024; 10.1101/2024.02.29.579666v1 | Per-sample (user-uploaded) | User samples, all pathogens | Host subtraction + NCBI alignment | Pathogen reports + visualizations | Yes - web-based sample reports with interactive visualizations | - | NCBI NT/NR alignment | All pathogens | Yes (SPAdes) | RPM (reads per million) | Cloud (free hosted platform) |
| **Hecatomb** | Roach et al. 2024; 10.1093/gigascience/giae020 | Per-study (user-supplied) | User samples, virus-focused | Assembly + tiered database search | Annotated viral contigs | - | - | MMseqs2 + BLAST (tiered) | All viruses | Yes (MEGAHIT) | Read counts | Local (Snakemake + Conda) |
| **kallisto translated search** | Luebbert et al. 2025; 10.1038/s41587-025-02614-y | Demonstrated on 3.4M runs | All SRA, virus-focused | Pseudoalignment (translated k-mers) | Virus detection calls + cell-level counts | - | Yes | Pseudoalignment to PalmDB (aa) | RNA viruses (RdRP-containing) | No (read-level only) | Per-cell or per-run pseudocounts | Local or cloud |
| **Logan** | Chikhi et al. 2025; 10.1101/2024.07.30.605881 | 27M accessions (all SRA) | All SRA, general purpose | De novo assembly of entire SRA | Assembled contigs (public S3) | Logan-Search (k-mer lookup) | - | K-mer indexing | All sequences | Yes (global unitigs/contigs) | Not per-run | Cloud (AWS) |
| **Serratus** | Edgar et al. 2022; 10.1038/s41586-021-04332-2 | 5.7M samples (all SRA) | All SRA, virus-focused | Read-level alignment to RdRP | Alignment summaries + palmprints | Serratus.io explorer (palmprint search, geographic/host range) | - | DIAMOND to RdRP/palmDB | RNA viruses (RdRP-containing) | No (read-level only) | Alignment depth (per-run) | Cloud (AWS, ~$0.01/sample) |
| **RNAquarium** | this study | 75k runs (one host species) | Single host species, full pipeline | Sequential host-filtering + assembly + BLAST/Diamond | Gene counts + non-host contigs + curated viral contigs + virus counts matrix | Yes - web portal with gene search, UMAP viewer, gene-centric expression plots, phylogenetic viewer, virus explorer, and AI chatbot for natural-language data queries | Yes | BLASTn (nt) + Diamond (nr) + Taxonomizr | All viruses + all non-host taxa | Yes (SPAdes, per-bioproject) | Salmon (per-run x per-contig) | HPC (Nextflow + SLURM) |

**Supplemental Table S2. Distribution of samples across infection classes, related to Supplemental Figure S11 and Figure 5.**

| Infection Class | Samples | Fraction |
|---|---|---|
| No infection | 54050 | 0.88196 |
| Zebrafish jaw poxvirus | 2537 | 0.04140 |
| Danio blood picornavirus/Zebrafish picornavirus 2 | 2150 | 0.03508 |
| Zebrafish picornavirus 1 | 1964 | 0.03205 |
| Zebrafish jaw picornavirus | 182 | 0.00297 |
| Spring viraemia of carp virus | 147 | 0.00240 |
| Cyprinid herpesvirus 3 | 70 | 0.00114 |
| Zebrafish gut calicivirus | 39 | 0.00064 |
| Zebrafish rhabdo-like virus | 35 | 0.00057 |
| Largemouth bass virus | 34 | 0.00055 |
| Zebrafish influenza B-like virus | 25 | 0.00041 |
| Rocky Mountain birnavirus | 24 | 0.00039 |
| Redspotted grouper nervous necrosis virus | 14 | 0.00023 |
| Zebrafish systemic calicivirus | 13 | 0.00021 |

**Supplemental Table S3. RNAquarium SRA metadata column descriptions, related to STAR Methods.**

| Column Name | Level | Source | Description |
|---|---|---|---|
| run.accession | Run | SRA | Unique SRA accession for the sequencing run (e.g. SRR...) |
| experiment.accession | Experiment | SRA | SRA experiment accession (e.g. SRX...) |
| sample.accession | Sample | SRA | SRA sample accession (e.g. SRS...) |
| study.accession | Study | SRA | SRA study accession (e.g. SRP...) |
| bioproject | Study | SRA | NCBI BioProject accession (e.g. PRJNA...) |
| study.title | Study | SRA | Title of the SRA study |
| study.alias | Study | SRA | Submitter-assigned alias for the study |
| study.type | Study | SRA | Study type classification |
| study.abstract | Study | SRA | Abstract describing the study |
| study.attributes | Study | SRA | Key-value attributes associated with the study |
| study.PMIDs | Study | SRA | PubMed IDs linked to the study |
| sample.description | Sample | SRA | Free-text description of the sample |
| sample.title | Sample | SRA | Title of the SRA sample |
| sample.alias | Sample | SRA | Submitter-assigned alias for the sample |
| sample.centername | Sample | SRA | Submitter/center name parsed from SRA sample XML |
| sample.attributes | Sample | SRA | Key-value attributes associated with the sample (often contains tissue, age, treatment, etc.) |
| GEOsample.title | GEO Sample | GEO | Title of the GEO sample |
| GEOsample.dataprocessing | GEO Sample | GEO | Data processing steps described in GEO submission |
| GEOsample.source | GEO Sample | GEO | Source material described in GEO submission |
| GEOsample.treatmentprotocol | GEO Sample | GEO | Treatment protocol from GEO submission |
| GEOsample.extractprotocol | GEO Sample | GEO | RNA/DNA extraction protocol from GEO submission |
| GEOsample.growthprotocol | GEO Sample | GEO | Growth/rearing protocol from GEO submission |
| GEOsample.characteristics | GEO Sample | GEO | Sample characteristics from GEO (key-value pairs, e.g. genotype, age, tissue) |
| GEOsample.accession | GEO Sample | GEO | GEO sample accession (e.g. GSM...) |
| experiment.title | Experiment | SRA | Title of the SRA experiment |
| experiment.alias | Experiment | SRA | Submitter-assigned alias for the experiment |
| experiment.library_name | Experiment | SRA | Library name as submitted |
| experiment.design_description | Experiment | SRA | Free-text description of the experimental design |
| experiment.library_construction_protocol | Experiment | SRA | Protocol used for library construction |
| experiment.attributes | Experiment | SRA | Key-value attributes associated with the experiment |
| experiment.library_strategy | Experiment | SRA | Library strategy (e.g. RNA-Seq, miRNA-Seq) |
| experiment.library_source | Experiment | SRA | Library source (e.g. TRANSCRIPTOMIC) |
| experiment.library_selection | Experiment | SRA | Library selection method (e.g. cDNA, size fractionation) |
| experiment.library_layout | Experiment | SRA | Library layout as reported by submitter (SINGLE or PAIRED) |
| experiment.platform | Experiment | SRA | Sequencing platform (e.g. ILLUMINA) |
| experiment.instrument_model | Experiment | SRA | Instrument model (e.g. Illumina HiSeq 2500) |
| experiment.spot_descriptor | Experiment | SRA | Technical description of read/spot structure |
| experiment.study_ref | Experiment | SRA | Study accession referenced by the experiment |
| run.title | Run | SRA | Title field for the sequencing run |
| run.attributes | Run | SRA | Key-value attributes associated with the run |
| run.filename | Run | SRA | Filename(s) associated with the run |
| run.semantic_name | Run | SRA | Space-separated list of file format tokens describing the run's files (e.g. "fastq fastq", "10x Genomics bam file fasta", "pacbio_native pacbio_native") |
| run.total_bases | Run | SRA | Total number of bases sequenced in the run |
| run.total_spots | Run | SRA | Total number of spots (reads) in the run |
| run.alias | Run | SRA | Submitter-assigned alias for the run |
| run.read_lengths | Run | SRA | Read length(s) for the run |
| run.base_counts | Run | SRA | Per-base counts for the run |
| run.r1_length | Run | SRA | Length of read 1 |
| run.r2_length | Run | SRA | Length of read 2 |
| run.r3_length | Run | SRA | Length of read 3 (if applicable) |
| run.r4_length | Run | SRA | Length of read 4 (if applicable) |
| run.Acount | Run | SRA | Count of A bases in the run |
| run.Ccount | Run | SRA | Count of C bases in the run |
| run.Gcount | Run | SRA | Count of G bases in the run |
| run.Tcount | Run | SRA | Count of T bases in the run |
| run.Ncount | Run | SRA | Count of N (ambiguous) bases in the run |
| run.experiment | Run | SRA | Experiment accession linked to this run |
| run.pool_member | Run | SRA | Pool member information if run is part of a pooled library |
| submission.accession | Submission | SRA | SRA submission accession (e.g. SRA...) |
| submission.srasource | Submission | SRA | SRA source/submitter information parsed from submission XML |
| submission.bioprojectsource | Submission | SRA | Submitter/center name parsed from BioProject XML |
| seqdetective.n_mates | Run | Seq-Detective | Number of FASTQ files evaluated by Seq-Detective: 1 for single-end, 2 for paired-end. Discrepancies with layout='paired' and n_mates=1 (or vice versa) indicate metadata errors or file availability issues. |
| seqdetective.mapping_rate.mate1 | Run | Seq-Detective | HISAT2 alignment mapping rate for mate 1 (or the sole file for single-end runs) on a 200k-read subsample aligned to the zebrafish reference genome. Proportion [0–1]. Null when only mate 2 was evaluated or the run was not processed. |
| seqdetective.mapping_rate.mate2 | Run | Seq-Detective | HISAT2 mapping rate for mate 2 on subsampled reads. Null for single-end runs. Large mate1/mate2 discrepancy may indicate a barcode-chemistry paired-end run or adapter contamination. |
| seqdetective.nofeature_rate.mate1 | Run | Seq-Detective | Fraction of mate 1 reads aligning to the genome but not assigned to an annotated gene feature (featureCounts 'no_feature'), on the subsampled reads. High values may indicate antisense preparation, ncRNA enrichment, or annotation gaps. |
| seqdetective.nofeature_rate.mate2 | Run | Seq-Detective | Fraction of mate 2 reads with no assigned gene feature. Null for single-end runs. |
| seqdetective.sparsity.mate1 | Run | Seq-Detective | Fraction of annotated genes with zero counts for mate 1 in the subsampled data. High sparsity is expected for droplet single-cell data; unusually high sparsity in bulk data may indicate low quality or index hopping. |
| seqdetective.sparsity.mate2 | Run | Seq-Detective | Gene count sparsity for mate 2. Null for single-end runs. |
| seqdetective.pos_strand_rate.mate1 | Run | Seq-Detective | Fraction of mate 1 reads mapping to the positive strand. ~0.5 indicates unstranded; ~1.0 or ~0.0 indicates stranded. Stored as string to accommodate 'NA' entries where strand information could not be computed. |
| seqdetective.pos_strand_rate.mate2 | Run | Seq-Detective | Positive-strand fraction for mate 2. For stranded paired-end libraries, mate 2 is typically antisense, so this value will be near 0.0 where mate 1 is near 1.0. Null for single-end runs. |
| seqdetective.readlen.mate1 | Run | Seq-Detective | Observed read length (bp) for mate 1 from the subsampled FASTQ. Reflects actual read length in subsample; may differ from the nominal length in SRA metadata. |
| seqdetective.readlen.mate2 | Run | Seq-Detective | Observed read length (bp) for mate 2. Null for single-end runs. For 10x Chromium and similar droplet protocols run at optimal length, mate 1 carries the barcode+UMI (28 bp in v3) while mate 2 carries the cDNA insert. |
| seqdetective.judgement.mate1 | Run | Seq-Detective | Seq-Detective read-type classification for mate 1 (or the sole file for single-end runs). 'B' = biological (transcriptomic reads from the host); 'T' = technical or non-mapping. Null when the run was not evaluated. |
| seqdetective.judgement.mate2 | Run | Seq-Detective | Seq-Detective classification for mate 2. Null for single-end runs. |
| seqdetective.judgement.reason | Run | Seq-Detective | Human-readable reason supporting the B/T judgement. Examples: 'usable mapping rate', 'biological fallback assumption', 'barcode-length mate'. Applies to both mates jointly. |
| platform_family | | Curated | Sequencing platform family resolved from instrument_model (lookup table) with platform field as fallback. Values: illumina, bgi, ont, pacbio, ion_torrent, legacy (454/SOLiD/Complete Genomics), element, unknown. |
| instrument_generation | | Curated | Instrument generation within platform family from instrument_model lookup. Representative values: early_illumina, hiseq_era, novaseq_era, nextseq, nextseq_v2, miseq, ont, pacbio_early, pacbio_modern, bgi. 'unknown' for unrecognised models. |
| read_bias | | Curated | Transcript coverage bias inferred by ugrep pattern matching on free-text protocol and title fields. Values: 3prime, 5prime, full_length, unknown. May be missing or incorrect if the submitter did not describe the bias in text. |
| selection_class | | Curated | Library selection method. Values: poly_a, rrna_depletion, random_priming, small_rna, cage, cdna_unspecified, dnase, size_fractionation, other, unknown. Primarily from experiment.library_selection; free-text hints used as fallback when that field is 'unspecified', 'cDNA', or 'other'. |
| prep_kit | | Curated | Library prep kit detected by ugrep on free-text fields. Values: trueseq, nebnext, nextera, lexogen, ribozero, smarter, unknown. 'unknown' even when a kit was used if its name is absent from the text. Lexogen is detected via both 'lexogen' and 'QuantSeq' mentions. |
| sc_or_bulk | | Curated | Single-cell vs. bulk classification. Values: sc (TRANSCRIPTOMIC SINGLE CELL library_source or named SC technology), sc_generic (generic scRNA-seq language only), bulk (TRANSCRIPTOMIC/METATRANSCRIPTOMIC with no SC signal), unknown. Resolution hierarchy: library_source > named SC technology score > generic scRNA-seq score. |
| tech_class | | Curated | Coarse technology grouping. Values: single_cell_droplet (10x, Drop-seq, inDrops, DroNc-seq), single_cell_plate (Smart-seq, CEL-seq, Fluidigm, and most plate-based methods), single_cell_generic (generic scRNA-seq signal only), bulk, clip, other_seq, unknown. Derived deterministically from technology. |
| technology | | Curated | Dominant sequencing technology resolved by weighted ugrep pattern matching across run (×1000), sample (×100), experiment (×10), and study (×1) free-text fields. Values include 10x, smartseq, dropseq, bulk, generic-scrnaseq-only, and ~15 others; 'unknown' when no pattern scores above zero. A named SC technology scoring ≥10 (experiment-level or stronger) suppresses bulk signal before resolution. |
| tech_variant | | Curated | Sub-variant of the detected technology. Currently only populated for sort-seq (tech_variant='sort_seq', technology='celseq'). Null for all other technologies. |
| submission.bioprojectsource.country | | Curated | Country of origin parsed from BioProject XML |
| earliest_date | | Curated | Earliest date associated with the run/sample, derived by taking the minimum across multiple SRA date fields |
| devstage_curation | | Curated | Curated developmental stage (fine-grained) assigned from metadata text |
| devstage_curation_coarse | | Curated | Curated developmental stage collapsed into broader categories |
| tissue_curation | | Curated | Curated tissue/organ type (fine-grained) assigned from metadata text |
| tissue_curation_coarse | | Curated | Curated tissue type collapsed into broader categories |

### Supplemental Table S4. RNAquarium System Components and Technologies, related to STAR Methods.

The RNAquarium portal follows a microservice architecture, with twelve independently developed and deployed services that share a common spine: a Next.js portal shell aggregates content and navigation, four independent visualisation apps render the scientific plots, three backend APIs serve the underlying data, three Model Context Protocol (MCP) servers expose that same data to the integrated AI chatbot, and one ETL pipeline transforms raw pipeline outputs into the DuckDB and Parquet files that the metadata and MCP services consume. The chatbot reaches the RNAquarium backends exclusively through these MCP servers, so natural-language questions are answered against the same live metadata and visualisation APIs that power the web viewers rather than a static snapshot.
The role and tech stack of each individual repository are described below.

**Web portal and interactive viewers (user-facing front-ends)**

| Service | Role | Technologies |
|---|---|---|
| rnaquarium-portal | Unified web entry point; landing page, per-gene dashboards, navigation to downstream viewers and data tools. | Next.js (App Router), React/TypeScript; MUI + @czi-sds/components (UI); Plotly.js (plots); Cytoscape.js (graphs) |
| rnaquarium-phylo-viewer-frontend | Interactive phylogenetic-tree and per-node transcript-count explorer | Next.js, React/TypeScript; MUI (UI); d3 + phylotree (tree rendering); newick (parsing) |
| rnaquarium-gene-embedding-viewer-fe | Gene nearest-neighbor search across three embedding spaces, rendered as ranked tables and interactive network graphs | Vite, React/TypeScript; MUI (UI); Cytoscape.js (graphs) |
| rnaquarium-gene-expression-viewer | Multi-gene expression comparisons (violin, heatmap, scatter, virus-conditioned robust-shift plots) | Streamlit; Scanpy + AnnData (expression matrices); Plotly (plots) |
| rnaquarium-sample-umap-viewer-fs | Interactive UMAP of the reprocessed SRA samples, with panels colorable by developmental stage, tissue, gene expression, or viral-read abundance | Streamlit + FastAPI sidecar; Scanpy + AnnData (expression matrices); Plotly (plots) |

**Backend APIs**

| Service | Role | Technologies |
|---|---|---|
| rnaquarium-phylo-viewer-backend | Prunes the NCBI taxonomy to organisms with detected transcripts and returns tree structures and counts to the phylo front-end | FastAPI; ETE3 (taxonomy, over the NCBI SQLite database) |
| rnaquarium-gene-embedding-viewer-be | Computes nearest-neighbor similarity across the RNAquarium GeneFormer, Zebrahub-fine-tuned GeneFormer, and ESM2 embedding matrices | FastAPI; h5py (matrix IO); scikit-learn + scipy (nearest-neighbor math); NetworkX (graph build); Plotly (plots) |
| rnaquarium-data-service-backend | Streams contig FASTA sequences by NCBI taxonomy ID, using closure tables to resolve sub-tree queries over the full taxonomy | FastAPI; PostgreSQL (storage) with closure tables over the NCBI taxonomy; ETE3 (taxonomy) |

**Data access and AI integration**

| Service | Role | Technologies |
|---|---|---|
| rnaquarium-metadata-server | Web-based SQL and faceted-search interface over the project's contig and SRA-run metadata | Dataset (web UI) + dataset-parquet (DuckDB reader) over DuckDB (storage) |
| rnaquarium-metadata-mcp-server | Exposes the DuckDB metadata as a SQL-query MCP tool The same codebase is deployed twice: as rnaquarium-db (RNAquarium contig/SRA DuckDB) and as zfin-db (ZFIN DuckDB snapshot) | Python MCP server over DuckDB (fork of the MotherDuck MCP server) |
| rnaquarium-viz-mcp-server | Wraps the four visualisation backends as 22 MCP tools (UMAP, gene-expression, embedding, phylo) for the AI chatbot | Python MCP server |
| rnaquarium-mcp-data-transformer | Offline ETL that converts raw SRA, contig-classification, gene-nomenclature and virus-sample tables into the DuckDB and Parquet snapshots consumed by the metadata servers | Python ETL; pandas (transforms); DuckDB (output) |
| biohub-chatbot | Multi-agent chat interface whose RNAquarium agent lets users query the project's data and run visualisations in natural language by delegating to the three MCP servers (rnaquarium-db, zfin-db, rnaquarium-viz) | Next.js, React/TypeScript; Vercel AI SDK (LLM orchestration, Anthropic); @modelcontextprotocol/sdk (MCP client) |
